# Implementation of an adaptive-optics assisted isoSTED nanoscope

**DOI:** 10.1101/2025.04.02.645322

**Authors:** Yang Li, Dong-Ryoung Lee, Edward S. Allgeyer, Lena K. Schroeder, Shijie Tu, Joerg Bewersdorf, Xiang Hao

**Affiliations:** Department of Cell Biology, Yale School of Medicine, New Haven, CT, USA; School of Mechanical Engineering, Soongsil University, Seoul, Republic of Korea; The Gurdon Institute, University of Cambridge, Cambridge, UK; Cellular Imaging Shared Resource, Fred Hutchinson Cancer Center, Seattle, WA, USA; State Key Laboratory of Extreme Photonics and Instrumentation, College of Optical Science and Engineering, Zhejiang University, Hangzhou, Zhejiang, China; Department of Biomedical Engineering, Yale University, New Haven, CT, USA; Department of Physics, Yale University, New Haven, CT, USA; ZJU-Hangzhou Global Scientific and Technological Innovation Center, Hangzhou, Zhejiang, China

## Abstract

The highest level of three-dimensional (3D) resolution in stimulated emission depletion (STED) nanoscopy involves harnessing a 4Pi architecture using two opposing objectives. This protocol describes the construction and alignment of a 4Pi-STED nanoscope, commonly referred to as an isoSTED nanoscope. It guides interested researchers through assembling optomechanical components, configuring electronic and control devices, aligning the optical beam path, and assessing the instrument’s performance. Designed for adept optical instrument builders, this protocol offers a detailed roadmap for constructing an isoSTED nanoscope with adaptive optics (AO) in approximately 12 months. With this finely calibrated instrument, researchers can achieve 3D biological images with isotropic sub-50-nm resolution in thick samples up to 35 µm in depth.

## 1. Introduction

Stimulated Emission Depletion (STED) nanoscopy, stemming from innovations first described in 1994^1^, stands as a pivotal technique that has redefined fluorescence imaging. Its first experimental demonstration in 1999^2^ and subsequent application in biological samples in 2000^3^ marked a turning point by showing that the diffraction limit of far-field light microscopy can be broken. Operating in a targeted switching mode, STED nanoscopy confines fluorescence to sub-diffraction-sized regions by depleting surrounding excited fluorophores via stimulated emission.

Originally limited to two-dimensional (2D) imaging, technical advances over the past ∼15 years have propelled STED nanoscopy into a new realm—3D imaging. In particular, using a so-called 4Pi geometry in which light illuminating the sample coherently through two opposing objectives facilitates a sharp central depletion minimum in the axial direction, affording a remarkable 33-nanometer *z*-resolution. This breakthrough, termed ‘STED-4Pi microscopy’^4,5^ laid the groundwork for the subsequent development of isoSTED nanoscopy^6^ which demonstrated 3D isotropic resolution by combining fluorescence confinement in all three dimensions. isoSTED nanoscopy yields an effective point spread function (PSF) 1,500 times smaller in volume than that of conventional confocal microscopy, achieving a striking 20–50-nm isotropic resolution. isoSTED nanoscopy has been demonstrated in material science applications for imaging self-assembled nanostructures^7^ and in cell biology, revealing sub-cellular structures in both fixed cells^8,9^ and living cells^10^.

However, despite its remarkable resolution, the original isoSTED implementation has limitations, particularly in the form of axially shifted side-lobes in its effective PSF that can generate axially repetitive artifacts. The adaptive-optics (AO) assisted isoSTED nanoscope described in this protocol, and hereafter is referred to exclusively, addresses these artifacts by introducing moderate defocus on the depletion beam^11^ coupled with AO corrections. These improvements facilitate isoSTED imaging in thick specimens^12^, greatly expanding the range of samples that can now be imaged, like brain tissue sections and drosophila egg chambers. These isoSTED nanoscopy developments were made possible through the collaboration of research laboratories at Yale University, University of Oxford, and Tufts University. Collaborative efforts continue to evolve and expand, now encompassing more institutions, such as ShanghaiTech University and Zhejiang University, in ongoing developmental endeavors.

While isoSTED nanoscopy has emerged as a potent imaging tool, its complexity requires considerable expertise and efforts across multiple disciplines for both construction and operation. In a bid to democratize access to this instrument, our protocol encapsulates a comprehensive guide—from fundamental principles to troubleshooting—enabling researchers to construct and maintain an isoSTED nanoscope. Essential resources, including part lists, schematics, computer-aided design (CAD) models, and software, are available on https://zenodo.org/uploads/10574469. We estimate that a skilled builder focusing full-time on this project would take ∼12 months to create the described instrument. We also encourage readers to contact this manuscript’s authors for updates and questions before or during the building process.

### 1.1. Conceptual description of the AO-isoSTED setup

An AO assisted isoSTED nanoscope features two opposing objectives in a vertically oriented 4Pi interference cavity. Similar to other STED modalities, in isoSTED nanoscopy, the fluorescence is restricted in 3D to a volume smaller than the diffraction limit by depleting excited fluorophores in the vicinity of the excitation volume using stimulated emission. To achieve this, the depletion laser generates a hollow, sphere-shaped focus to superimpose on the excitation focus (**Fig. 1**). To achieve uniform compression of the fluorescence emission volume, two depletion patterns, *i.e.* STED*_xy_* for lateral and STED*_z_* for axial depletion, with a common focal zero are combined. Specifically, the STED*_xy_* pattern is generated by the constructive interference of two ring-shaped foci, whereas the STED*_z_* beams interfere destructively along the z-axis in the focal plane by tightly focused Gaussian-like beams. The incoherent summation of these two patterns creates a ‘zero’-intensity minimum at the center surrounded by a high-intensity crest in all directions.

**Fig. 1.**
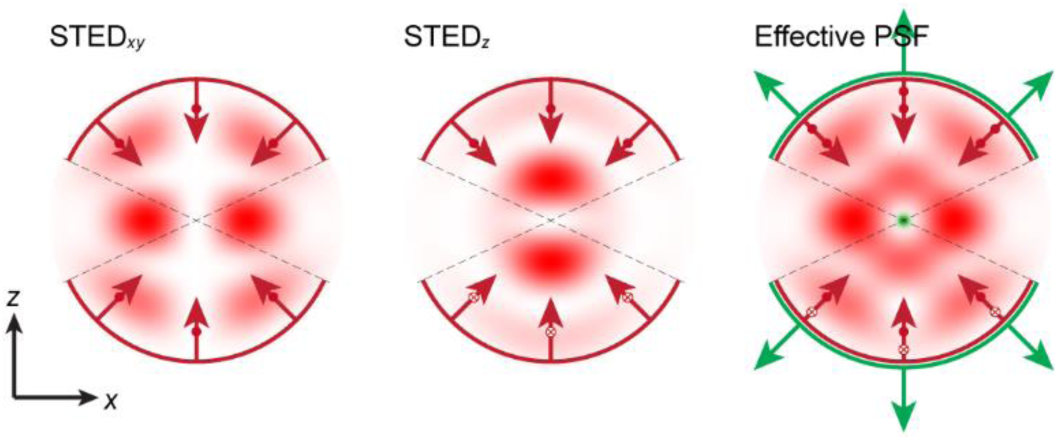
PSFs in AO-isoSTED nanoscopy in axial (*xz*) plane. From left to right: STED*_xy_*, STED*_z_*, and the combined depletion (red) and effective (green) PSFs. The small hollow arrows beside the large arrows denote the polarization state of the depletion beam when generating the respective PSFs. Specifically, the STED*_xy_* PSF is created through constructive interference of the two counterpropagating beams, whereas the STED*_z_* PSF is formed through destructive interference.

For optimal performance, it is crucial to maintain stable interference patterns throughout the imaging procedure. Addressing and mitigating potential disruptive factors, notably aberrations, is imperative. As the isoSTED nanoscope images deeper into specimens, refractive index variations can lead to aberrations that make imaging impossible. To facilitate super-resolution imaging of thick specimens, the isoSTED nanoscope implements a sensorless AO architecture, aimed at correcting aberrations induced by the specimens.

#### Optical design overview

The AO-isoSTED nanoscope uses two opposing 100×/1.35 numerical aperture (NA) silicone-oil immersion objectives (Olympus, UPLSAPO 100XS), designated as OBJ1 and OBJ2 in **Fig. 2**, installed on a vertical optical breadboard. Stable interference across the entire field of view (FOV) relies on precise alignment of the two objectives with nearly identical magnification (±0.3%). A quarter-wave plate (QWP) (Thorlabs, AQWP05M-600) set at 45° with respect to the vertical breadboard plane is positioned immediately behind each objective (QWP4 for OBJ1 and QWP5 for OBJ2). The QWPs convert the linearly polarized incident beams to circularly polarized ones, minimizing excitation and depletion selectivity regarding fluorophore dipole orientation and reducing polarization-induced background^12^ in the recorded images. The back pupil plane of each objective is conjugated to a deformable mirror (DM1 for OBJ1 and DM2 for OBJ2) (Boston Micromachines, Multi-DM5.5) using a 4f telecentric system (L10, L11 for OBJ1 and L12, L13 for OBJ2). The DMs function as the main devices for correcting aberrations, particularly those induced by the specimen. To ensure that the reflection angles of the beams stay within acceptable limits for the DMs (±15°), a pair of mirrors folds the beam path inside the 4f telecentric system (M11, M12 for DM1 and M13, M14 for DM2). Notably, M11 is mounted on a single-axis piezo translation stage to adjust the optical path difference for the zero phase retardance of interference.

**Fig. 2.**
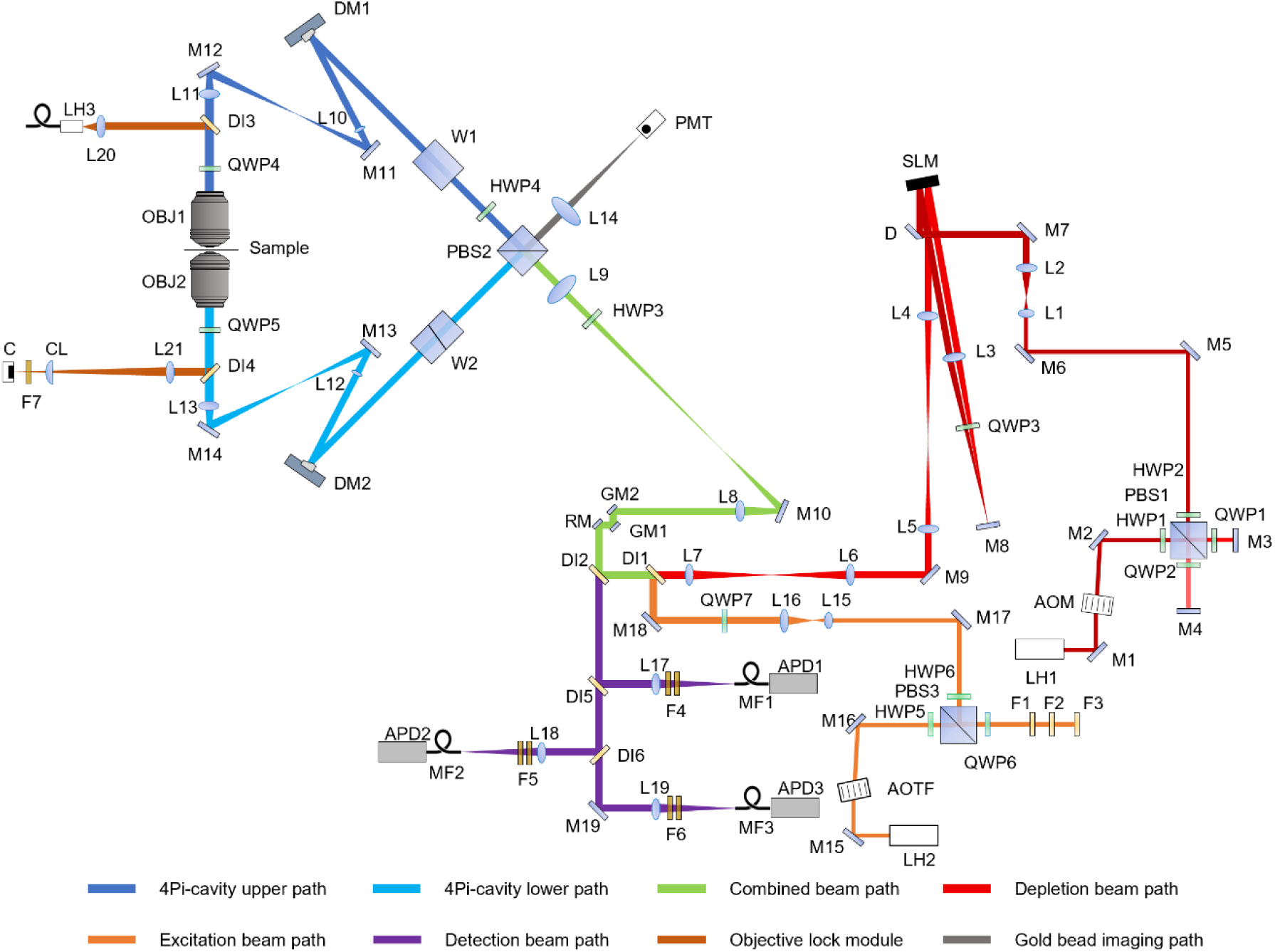
Optical beam path overview. Abbreviations: AOM, acousto-optic modulator; AOTF, acousto-optic tunable filter; APD, avalanche photodiode; C, camera; CL, cylindrical lens; D, D-shape mirror; DI, dichroic mirror; DM, deformable mirror; F, filter; GM, galvanometer mirror; HWP, half-wave plate; L, lens; LH, laser head; M, mirror; MF, multimode fiber; OBJ, objective; PBS, polarizing beam splitter cube; PMT, photomultiplier tube; QWP, quarter-wave plate; RM, resonant mirror; SLM, spatial light modulator; W, BK-7 glass wedges/optical flat.

Before being reflected by DM1, laser light passes through a stationary BK7 optical flat (W1) (UVISIR, custom designed) and a half-wave plate (HWP4) (Thorlabs, AHWP10M-600). In the other beam path, a BK7 wedge pair (W2) (UVISIR, custom designed), with one wedge mounted on a translation stage (Thorlabs, MTS25/M-Z8), replaces the optical flat, and the HWP is absent. A polarizing beam splitter cube (PBS2) (CVI Laser Optics, PBSH-450-1300-100) splits the laser light into these two beam paths. This PBS, along with all the previously described components, constitutes the core part of the isoSTED nanoscope known as the 4Pi interference cavity. Adjusting the optical path length difference between the upper and lower arms in this cavity is possible either by translating M11 or by adjusting the variable wedge in W2. Ideally, the optical path length difference is zero, eliminating any phase retardance for interference.

The coalescent illumination beams, comprising both depletion and excitation, enter the 4Pi interference cavity through the bottom-right side of PBS2. A white-light laser (LH2) (NKT Photonics, SuperK EXTREME EXR-20) provides the 78-MHz pulsed excitation light. Immediately after the laser, an acousto-optic tunable filter (AOTF) (Quanta-Tech, AOTFnC-400.650-TN) selects for the desired excitation components at wavelengths of 650 nm, 590 nm, and 488 nm. Following compensation for the chromatic dispersion within a pulse (see ‘Chromatic dispersion compensation module’ for details) and beam size expansion (using L15 and L16), the excitation beam is merged with the depletion beam path via a dichroic mirror (DI1) (Chroma, T740lpxr). A pair of wave plates (HWP6 and QWP7) tunes the polarization orientation of the excitation beam to 45° relative to that of the depletion beam so that the entire excitation beam can propagate through the upper arm in the 4Pi interference cavity.

A pulsed 775-nm wavelength laser (LH1) (MPB Communications, PFL-80-3000-775-B1R) operating at 78 MHz, with a pulse duration of approximately 0.9 ns and synchronized to the excitation laser, serves as the depletion source. The beam intensity is tunable via the acousto-optic modulator (AOM) (Quanta-Tech, MTS40-B2A3-750.850) mounted after LH1. To mitigate interference between the two polarization components later in the common focus between the objectives (**Fig. 1**), one polarization component of the laser is delayed with respect to the orthogonally polarized component via a polarization separation module (see ‘Polarization separation module alignment’ for details). Following beam size expansion using a 4f telecentric system (L1 and L2), the beam emerging from the polarization separation module illuminates a spatial light modulator (SLM) (Hamamatsu, X10468-02). A double-pass configuration^13^ encodes a vortex and a moderate defocus phase pattern (hologram) onto the two orthogonally oriented polarization components of the depletion beam. These two holograms are optically conjugated to each other and are further imaged onto a 16-kHz resonant mirror (RM) (EOPC, SC-30) through two additional 4f telecentric systems (L4 and L5 for the first 4f; L6 and L7 for the second). A mirror (M9) is placed between these two 4f telecentric systems to fold the beam path.

After merging, both the depletion and excitation beams illuminate the scanner, where the resonant mirror scans the combined laser beams along the fast-scanning axis (*x*), while two synchronized galvanometer scanning mirrors (GM1 and GM2) (SCANLAB, dynAXIS XS) scan the beams along the slow-scanning axis (*y*). Collectively, the two galvanometer mirrors behave as a single scan mirror, allowing the three scanning mirrors to act as a fast, dual-axis scanning system conjugated to the back pupil planes of the objectives. Imaging the resonant mirror onto the DMs in the 4Pi interference cavity via a 4f telecentric system (L8 and L9) guarantees the conjugation. A mirror (M10) between L8 and L9 folds the beam path from the optical table to the vertical breadboard plane. After this mirror, HWP3 rotates the two polarization orientations of the incident depletion beam to ±45° with respect to the polarization axis of PBS2, enabling precise 50/50 power allocation of both polarization components between the two arms of the 4Pi interference cavity. Simultaneously, because of its incident 45° linear polarization, the excitation beam is rotated to 0° by HWP3 and is therefore only coupled into the upper arm. HWP4 further rotates the polarization of the beams in the upper arm for subsequent interference in the combined focus.

The sample is mounted at the common focal plane of the objectives. Fluorescence from the sample is collected by both objectives, combined at PBS2, de-scanned by the scanner, and then separated from the common beam path via a custom-made, 5-mm-thick quad-bandpass dichroic mirror (DI2) (Chroma, ZT485/595/640/775rpc). Two additional dichroic mirrors (DI5 and DI6) (Chroma, ZT640rdc and ZT568rdc) split the fluorescence into three detection bands for multi-color imaging. For each band, two filters of the same type (F4, F5, and F6) reject stray excitation and depletion laser light (Semrock, FF03-525/50-25, FF01-624/40-25, and FF02-685/40-25). The filters in each detection band are separated by ∼50 mm and mounted at a small angle relative to each other to avoid interference between them. Fluorescence in each detection band is coupled into a multimode fiber (Thorlabs, FG105LCA) acting as a confocal pinhole, with a pinhole size of around 0.8 Airy units. Each fiber is connected to a single-photon counting avalanche photodiode (APD1, APD2, and APD3) (Excelitas, SPCM-ACRH-13-FC).

The upper-right side of PBS2 is not used for acquiring STED fluorescence imaging; instead, it serves as the outlet for the intense depletion laser after passing through the 4Pi interference cavity. A photomultiplier tube (PMT) (Hamamatsu, H10682-01), roughly conjugated to the objective focal plane, detects the non-de-scanned photons without going through a confocal detection pinhole. This allows the instrument to acquire illumination PSFs of the excitation or depletion laser foci without the influence of the detection PSF of the optical system^14^. This channel is utilized to assess the laser foci for aberration corrections.

The optical components installed at the back of the vertical breadboard are used to lock the positions of both objectives. A near-infrared laser (LH3) (Thorlabs, LP980-SF15) is collimated by a lens (L20) and directed towards the objectives by a dichroic mirror (DI3). When the two objectives share the focal plane, the laser beam passing sequentially through OBJ1 and OJB2 is collimated. This beam is then reflected by DI4 and focused by L21 onto a camera (C) via a cylindrical lens (CL), intentionally introducing astigmatism to allow for easy detection of the collimation status.

The isoSTED nanoscope function requires meticulous optical engineering and precise alignment, especially aligning multiple planes to be conjugated and co-centered, including: (i) the apertures of the objectives, (ii) the deformable mirrors, (iii) the resonant scan mirror, and (iv) the two circular holograms on the SLM for STED*_xy_* and STED*_z_* beams, respectively. Moreover, the conjugation of the detection pinhole with the common focus of the objectives in 3D is crucial. Detailed optical design specifications generated using Zemax (Ansys) can be accessed at: https://zenodo.org/uploads/10574469.

#### Mechanical Design Overview

The mechanical design (**Fig. 3a**) presented here prioritizes: (i) mechanical stability, (ii) easy maintenance, (iii) precise alignment, and (iv) simple use. For these purposes, the optical system was initially designed using Zemax, and then the mechanical structure was crafted around it using SolidWorks (Dassault Système) to secure all optical components. According to this design blueprint, the system was assembled using commercially available optomechanical elements as well as custom-designed, precision-machined parts to allow for precise alignment and minimize drift and vibrations. This strategy has been used to build working systems at Yale University and ShanghaiTech University. We provide an overview of the mechanical design in the following paragraphs. For more specific information regarding individual components and their assembly, we refer the reader to Section ‘Equipment setup’.

**Fig. 3.**
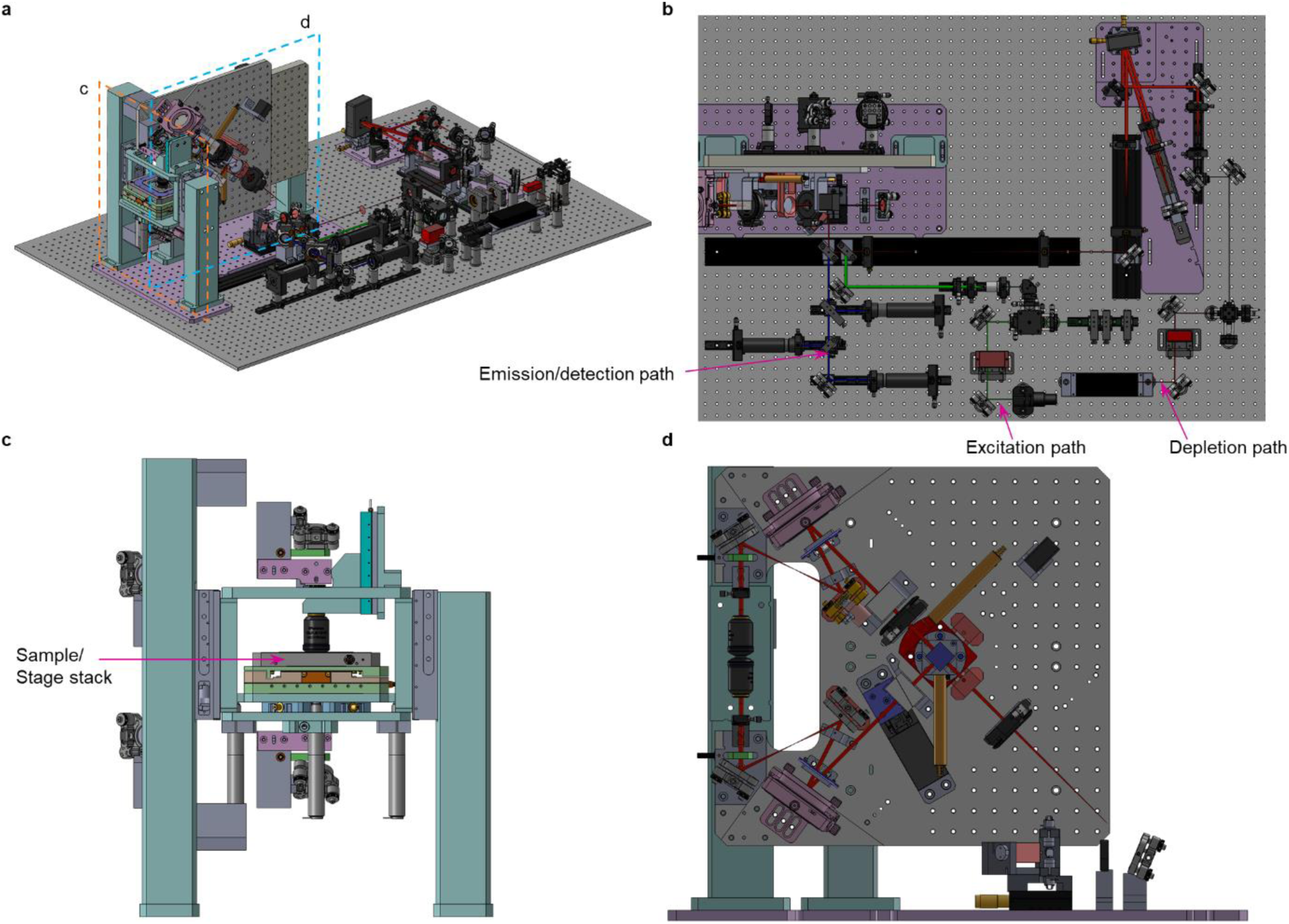
Mechanical design overview. **a**, CAD rendering of isoSTED nanoscope. **b**, Top view. **c**, side view from the left showing the stage stack. **d**, side view from the front showing the 4Pi interference cavity.

The mechanical design of the instrument is composed of two main bodies: components arranged horizontally on an optical table and those on a vertical optical breadboard. The components on the optical table can be categorized into depletion, excitation, and detection paths, as shown in **Fig. 3b**. The components on the vertical breadboard can be categorized into 4Pi interference cavity beam path and a so-called tower assembly which holds the sample and objectives, as shown in **Fig. 3c-d**. The vertical beam path configuration allows biological samples to be positioned in a conventional horizontal orientation, where the sample is sandwiched by two objectives in the direction of gravity.

The key part among the optics built horizontally on the optical table is a double-pass SLM configuration. The SLM is held by an *xy*-linear stage and a custom mount which can be rotated along *z* axis. The oblique 4f telecentric system is placed on a custom rail and a custom board which defines the SLM’s angle. The remaining optics on the optical table are easily adjustable because they are mounted by optical cage and rail systems. However, a primary design constraint is that the height of all components should be centered on the midpoint of the SLM’s active area. Additionally, within the depletion beam path, custom mounts are specifically designed to hold translation lens mounts perpendicular to and kinematic mirror mounts at a 45° angle to the optical axis. This design ensures precision, as the alignment accuracy of the depletion optics is crucial for achieving high-quality STED imaging.

The vertical optical breadboard is mounted to the optical table with two pillar legs. The breadboard has tapped holes for dowel pins based on the optical design, and the optical components attached to it are held by custom mounts which are screwed down to the breadboard. Therefore, the components can be fixed in the designated position with high precision. This approach also allows high repeatability for the removal and reinstallation of components. The key movable part on the breadboard is the PBS. The tilt and azimuthal angles of the PBS are controlled by a motorized goniometer and a motorized rotation stage, respectively. In the upper arm of the 4Pi interference cavity, a piezo actuator (Physik Instrumente (PI), P-841.10) serves for translating the attached mirror at sub-nanometer resolution, thereby optimizing the phase difference of interference at the common focal plane by tuning the optical path length difference between the upper and the lower arms. Additionally, two DMs for aberration correction are aligned to be co-centered with both the SLM and each objective back pupil. The second section of the vertical configuration is the stage stack in the tower assembly. In this assembly, the upper objective is translated along the optical axis through a long-range piezo linear stage (PI, N-565.260), enabling nanometer-level precision in positioning, while affording 26-mm travel range to accommodate easy sample insertion and removal. The lower objective is mounted on an *xy* piezo stage (PI, P-612.2SL). In combination, these two piezo actuators allow for the relative alignment of the two objectives in all three dimensions. The sample is secured atop a pair of piezo stages, comprising of an *xy* stage (PI, U-751.24) and a *z* stage (PI, P-541.ZCD). This assembly is supported by three high-resolution linear actuators (PI, M-227.10) that provide coarse sample *z*-positioning and tip/tilt leveling. The alignment strategy for this stage stack involves positioning the sample in the focal plane of the lower objective, followed by positioning the upper objective to ensure the convergence of both objectives onto the same focal plane. Lateral alignment is then achieved by optimizing the *xy* position of the lower objective. After precisely aligning both objectives, their positions are optically locked during the imaging process (see Section ‘Focus-lock module’).

Detailed mechanical drawings and CAD models generated using SolidWorks can be accessed at: https://zenodo.org/uploads/10574469. **Fig. 4** shows the optical beam path overlaid onto the mechanical layout of the instrument. This model is drawn to scale and intended to show the relationship between the optical components and their position within the instrument.

**Fig. 4.**
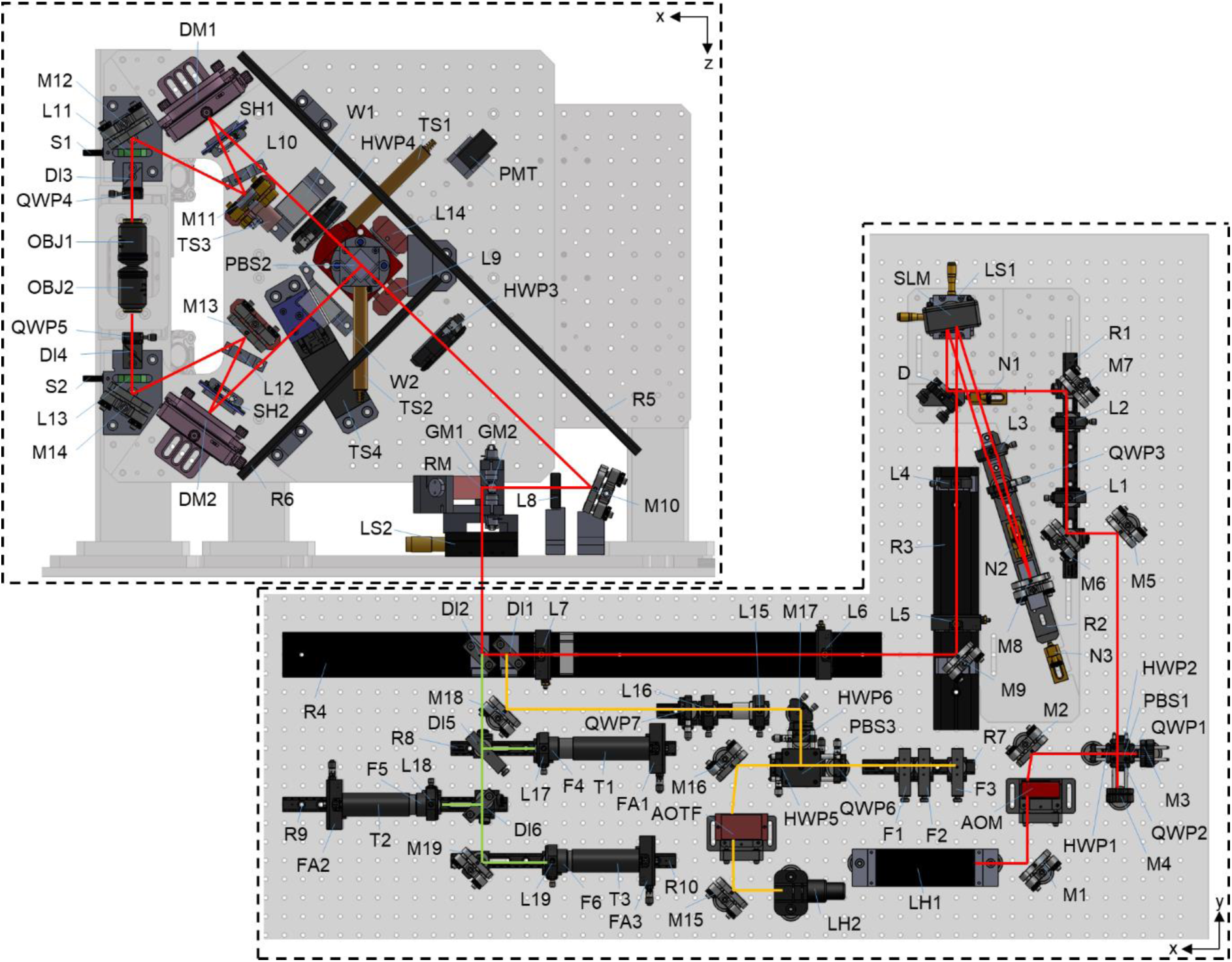
Labeled diagram of isoSTED optical beam path. Abbreviations: FA, fiber adapter; LS, linear stage; N, nudger; R, optical rail; S, screws; SH, shutters; T, tube; TS, motorized translation stages. The remaining labels are the same as those presented in Fig. 2. Note that the vertical optical breadboard shown on top-left is rotated in this diagram to a horizontal orientation for displaying purposes.

#### Control design overview

The isoSTED instrument control is centered around a single Microsoft Windows-based personal computer (PC) for data collection, synchronization, and device control (**Fig. 5**). Specifications for the instrument control PC are discussed in ‘Computer and data storage’. Notably, the computer motherboard must have at least two peripheral component interconnect express (PCIe) slots for the use of two National Instruments (NI) field programable gate array (FPGA) data acquisition (DAQ) boards that form the core of the image acquisition hardware. Specifically, an NI PCIe-7852R DAQ FPGA board is employed for data collection while an NI PCIe-7841R DAQ FGPA board is employed for waveform generation and synchronization. The details and connections of these FPGA boards are discussed in ‘FPGA Control’. The control and image acquisition are coordinated through an instrument control interface programmed in the LabVIEW environment (freely available on Zenodo at https://zenodo.org/uploads/10574469) and specifically tailored for controlling the isoSTED nanoscope. Modular control is employed for the SLM, deformable mirror, synchronization, time gating, and focus lock while additional device control is integrated into a single, easy to navigate, unified user interface accessible to non-experts.

**Fig. 5.**
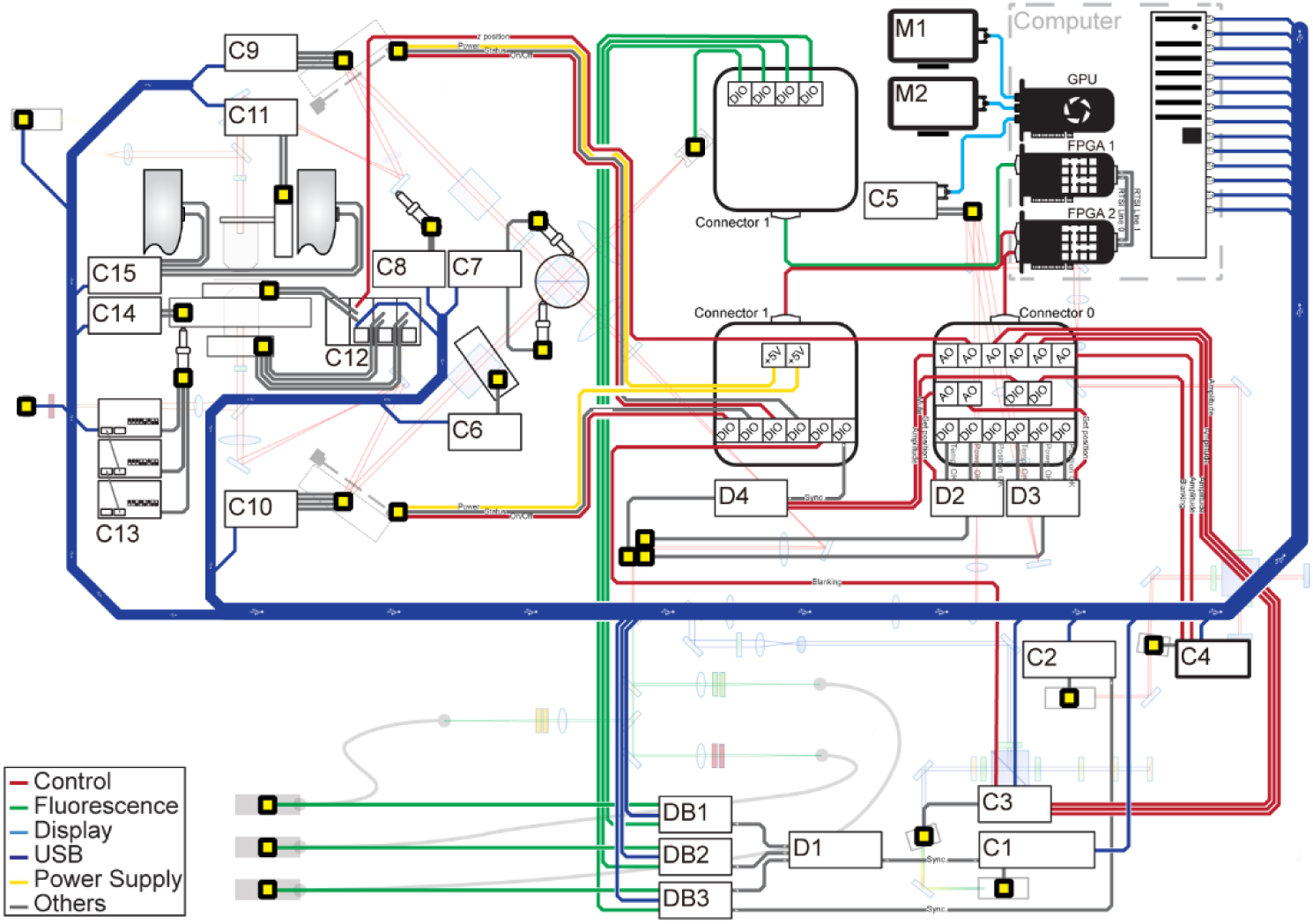
Control design of isoSTED nanoscope. Abbreviations: AO, analog output; AOM, acousto-optic modulator; AOTF, acousto-optic tunable filter; C1, controller of white-light excitation laser (NKT, SuperK EXTREME EXR-20); C2, controller of 775-nm depletion laser (MPB Communications, PFL-80-3000-775-B1R); C3, controller of AOTF (Quanta Tech, AOTFnC-400.650-TN); C4, controller of AOM (AA Opto Electronic, MTS40-A3-750.850); C5, controller of SLM (Hamamatsu, X10468-02); C6, controller of motorized linear stage (Thorlabs, MTS25/M-Z8); C7, controller of vernier micrometers (SM-25, Newport); C8, controller of piezo actuator (PI, P-841.10); C9-10, controller of DM (Boston Micromachines, Multi-DM5.5); C11, controller of objective mount (PI, N-565.260); C12, chassis of E-621 modules; C13, controller of linear actuator (PI, M-227.10); C14, controller of *xy* piezo stage (PI, U751.24); C15, controller of linear stage pair (Applied Scientific Instrumentation (ASI), LS-50); D1, fanout line driver; D2-D3, driver of galvanometer mirror (CTI, dynAXIS XS); D4, driver of resonant mirror (EOPC, SC-30), DB1-DB3, delay board (Opsero, custom designed); DIO, digital input/output; DM, deformable mirror; M1-M2, monitor; USB, universal serial bus.

### 1.2. Applications

The isoSTED nanoscope has achieved isotropic 3D sub-50 nm resolution in a variety of biological samples. One-color isoSTED imaging has shown up to 3-µm thick images of subcellular structures including microtubules, synaptonemal complexes, and mitochondria demonstrating high resolution isotropic renderings of a singly labeled targets^12^. Two-color isoSTED imaging has shown relationships between structures such as mitochondria and ER^12^, Golgi apparatus and Coatomer protein complex I (COPI)^12^, as well as coilin and survival of motor neuron proteins (SMN) in a Cajal body^15^. Implementing adaptive optics into isoSTED nanoscopy enabled two-color imaging in tissues like Drosophila egg chamber ring canals and 30-μm-thick mouse hippocampal brain sections^12^.

### 1.3. Advantages and limitations

isoSTED nanoscopy has so far achieved the highest 3D isotropic resolution among all STED technologies, thus providing an appealing tool for imaging biological structures. Like other far-field super-resolution fluorescence microscopes, the isoSTED can image biological structures with the high specificity enabled by immunofluorescence labeling. Similar to other STED approaches, isoSTED offers super-resolution imaging without post-processing and straightforward multi-color imaging. Two-color STED imaging using the same depletion laser, is particularly suitable for biomolecule co-localization studies as the two channels are intrinsically free of chromatic mismatch. Thanks to detection pinholes, which suppress out-of-focus fluorescence, isoSTED is better suited for thick-tissue imaging compared to other 3D super-resolution microscopy techniques such as single molecule localization microscopy (SMLM) or structured illumination microscopy (SIM). Utilizing AO, the isoSTED instrument can correct aberrations induced by the system as well as the sample to enable optimal imaging conditions even in thick and complex tissues.

However, the use of isoSTED nanoscopes has been limited by the complexity of the optical apparatus. Installation and operation of the instrument necessitates mastery of specialized alignment techniques and complex control software. Samples require sub-diffraction limit gold nanoparticles and/or fluorescent beads added before imaging to enable aberration correction by the sensorless AO, which can constrain or complicate sample preparation. As for all STED nanoscopys, specific fluorescent dyes are required for optimal image resolution, limiting the choice of reagents. Additionally, the use of high-NA objectives requires refractive-index matching during sample preparation, adding yet another layer of complexity to the experimental protocol. These instrument and sample challenges underscore the need for a detailed protocol, such as this, to successfully navigate and utilize isoSTED’s capabilities.

In summary, this nanoscope offers exceptional 3D resolution below 50 nm isotropically in a wide range of samples. The guidance provided in this protocol makes this complex technology more accessible.

### 1.4. Procedure overview

Building an isoSTED nanoscope requires meticulous attention to detail. While the order of some steps is flexible, many are interdependent and require a linear process. This protocol operates under the presumption that all essential components are on hand.

Prior to starting this protocol, it is highly recommended to thoroughly review the recommendations provided in Section ‘Experimental design’. This section describes fundamental prerequisites, including the laboratory space, temperature, and vibration requirements as well as the optical table selection. Additional information can be found in Section ‘Equipment setup’, which discusses procedures such as the installation and alignment of individual optical components.

The experimental setup begins with the installation of software (Steps 1**–**4), followed by the assembly of all optomechanical components (Steps 5**–**6) and all electronic connections (Steps 7**–**8). Subsequently, the depletion laser is mounted on the optical table to initiate the alignment of its beam path (Steps 9**–**41). Given the critical function of the depletion laser, its optical path serves as the reference for aligning the entire 4Pi interference cavity. Upon successful depletion path alignment, the excitation light path alignment is performed (Steps 42**–**62). Notably, the excitation and depletion beams should travel horizontally at the same height across the horizontal board referring to the center of the SLM. More precise tuning is available at each specific site to meet the designed requirement.

After combination of the excitation and depletion beams, the 4Pi interference cavity components are mounted on the front side of the vertical breadboard, and the 4Pi interference cavity is aligned (Steps 63**–**118). Once this is accomplished, the focus-lock module is installed on the back side of the same vertical breadboard (Steps 119**–**125). For optimal focus-lock results, a sample is inserted between the two opposing objectives during this process. The sample preparation procedure is described by Steps 126**–**141.

Next, using this test sample, the detection path is aligned (Steps 142**–**157), which is the final step in setting up the hardware. Subsequent procedures, including adjustments to the pulse gate/delay module and the AO module settings (Steps 158**–** 182), are performed exclusively within the control software interface.

Since the isoSTED nanoscope is a laser scanning microscope, the scanner, *i.e.* the combination of resonant mirror and galvanometer mirrors, needs to be calibrated to avoid image distortions and quantify the magnification of the instrument (Steps 183**–** 190). Finally, both fluorescent beads and biological samples are imaged to assess the achievable resolution and evaluate the performance of the instrument under typical imaging conditions, respectively (Steps 191**–**201).

#### General beam alignment

To facilitate easier construction of the isoSTED nanoscope, we have implemented a rail system designed to be a fixed reference for beam direction. Dovetail rails (*e.g.*, Thorlabs, RLA450/M) are frequently used for assisting the beam alignment. An alignment camera (Thorlabs, DCC1545M) that can be moved along the rail is used to monitor the laser beam’s position along this path. Notably, three common beam alignment procedures are regularly used while building the isoSTED nanoscope and are detailed below:

(i). Box 1: Single-mirror beam direction alignment;
(i). Box 2: Dual-mirror beam path alignment;
(i). Box 3: 4f telecentric system alignment.

These methods underscore the importance of precise beam control and alignment procedures in the construction process of the isoSTED nanoscope, ensuring optimal performance and reliability. For the other beam alignment that the precision requirements are not discussed explicitly, a ±0.5-mm-precision suffices in general. ▴ Critical The repeatability of mounting the alignment camera on the rail affect the alignment’s precision. Make sure to mount the camera on the rail properly during the process. The accuracy to identify the beam center affects the alignment precision as well. The methods in Box 1 and 2 are only compatible with collimated or slightly focused beams and may not work well with tightly focused laser beams.

###### Box 1 Single-mirror beam direction alignment

1. Mount the alignment camera on the rail near the beam source. Secure it in place and mark the current beam position with the cursor as the initial target position in the camera software.
2. Move the alignment camera along the rail in the beam propagation direction. If the beam is tilted relative to the rail axis, the beam position on the camera will shift.
3. Adjust the angle of the mirror to move the beam to the cursor position.
4. Move the alignment camera back to the near end of the rail (towards the mirror) and check if the current beam position is shifted off the target (cursor) position.
5. If yes, update the cursor position to the current beam position. Repeat Steps 1–4 until the beam position does not change on the camera anymore when translating the camera along the rail.

**! Note #1** The procedure in Box 1 aligns the beam direction along the rail axis while the beam position has been determined by the beam position when it reaches the mirror. Box 2 describes a procedure designed to align both.

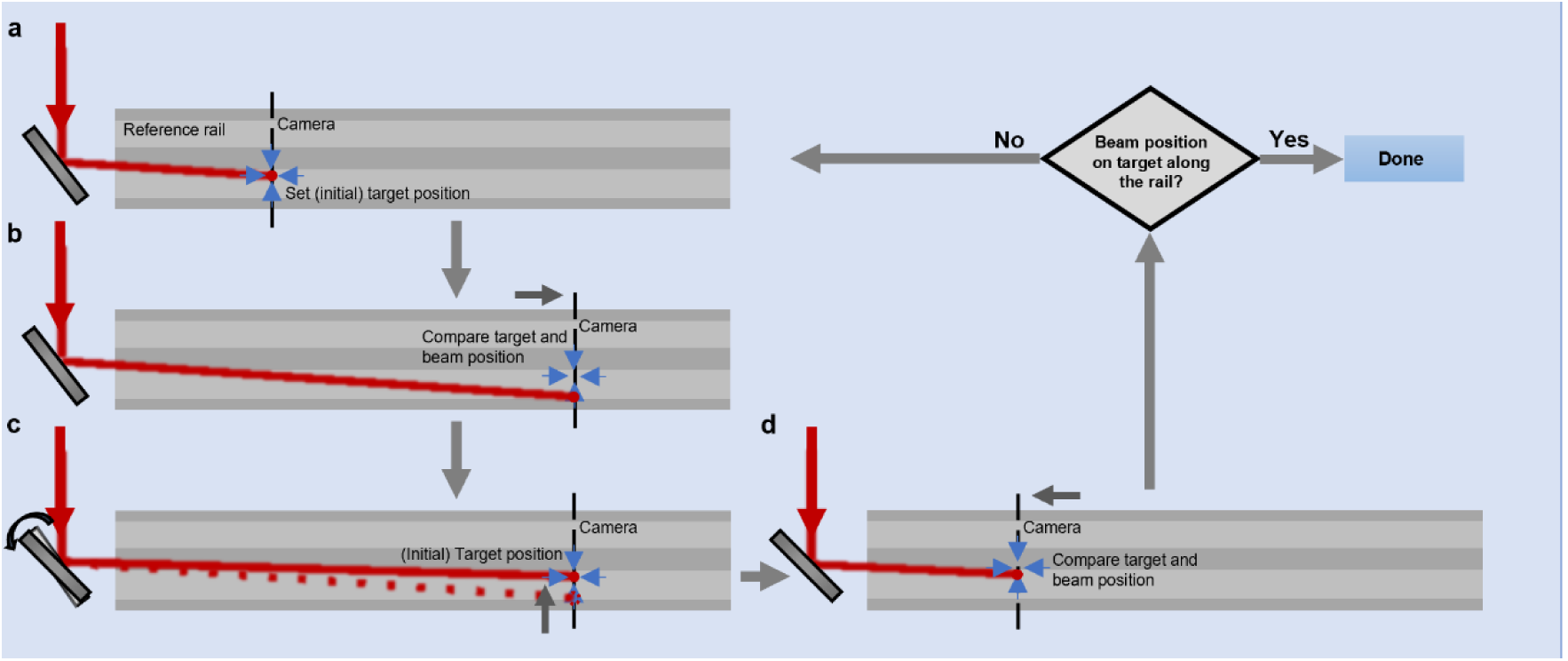

###### Box 2 Dual-mirror beam path alignment

1. Position the alignment camera at the near end of the rail toward the light source. Secure it in place. The ideal position the beam should target on the camera with the cursor in the software.
2. Adjust the angle of the first mirror (M1 in drawing below) to move the beam to the cursor (target) position.
3. Move the camera to the far end of the rail.
4. If the beam deviates from the cursor position, adjust the angle of the second mirror (M2 in drawing below) to compensate for the observed shift.
5. Move the camera back to the near end of the rail.
6. If the beam position is shifted off the cursor (target) position on the camera, repeat Steps 1–5 until the beam position is at the target position on the camera along the rail.

**! Note #1** It is unlikely that the manufacturer precisely centered the camera chip in its case. The ideal position the beam should target is therefore not necessarily the center pixel of the chip. This position needs to be determined independently, *i.e.*: by an iris on the optical mount.

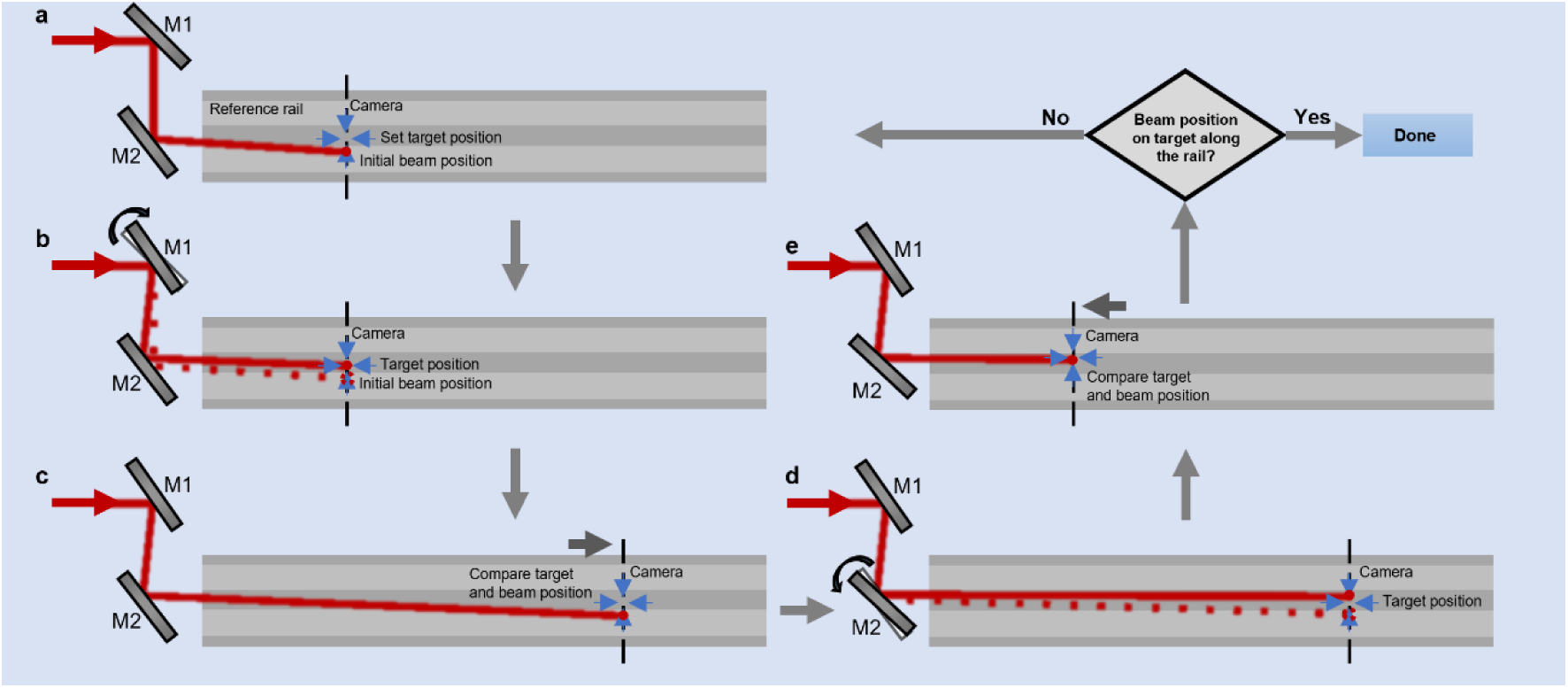

###### Box 3 4f telecentric system alignment

The alignment of three factors is imperative for each 4f telecentric system: (i) collinearity of the optical axes of both lenses; (ii) distance between the two lenses adjusted to achieve beam collimation; (iii) beam centered on both lenses. Note that the drawing emphasizes the beam center tracing and ignores the focusing effect on the beam size by the lenses. The following alignment procedure, suitable for systems where both lenses are mounted in *xy* translation mounts, is designed to achieve all three targets:

1. Place the camera at the far end of the rail, well past the second lens position, and secure it in place. With the two lenses removed, mark the center of the beam on the camera using the cursor in the camera software.
2. Insert the first lens and verify that the beam is incident to the lens at a normal angle. Translate the lens in its lateral direction until the beam is centered on the cursor position.
3. Insert the second lens and verify that the lens mount is oriented at a normal angle with respect to the outgoing beam. Translate the second lens in the direction perpendicular to the optical axis until the beam is centered on the cursor position.
4. Adjust the distance between the two lenses until the outgoing beam achieves collimation by either checking with a shear plate for sufficient coherent (depletion) beams, or by checking that the beam size not changing when moving a camera along the propagation direction.

**! Note #1** Enhancing the localization precision of the depletion beam center post SLM modulation involves incorporating a top-hat shaped 0/π hologram onto the SLM. This addition leverages the Talbot effect, resulting in a periodic focusing phenomenon along the propagation axis of the beam. Upon transmission through a lens, the absence of positional changes in the focal spot serves as confirmation of co-centered alignment between the beam and the lens.

**! Note #2** In our isoSTED design, the angle of the lenses is inherently fixed and non-adjustable. To ensure precise alignment, we have implemented a dedicated datum plane within the lens mount (**Supplementary Fig. S1**). This datum plane serves as a reference to define the orientation of the lens mount, effectively determining the alignment of the lens within the optical system. This design approach guarantees stability and accuracy in the positioning of each lens, contributing to the overall reliability and performance of the isoSTED nanoscope.

**! Note #3** To assess the collimation of the depletion beam in the isoSTED nanoscope we use a shear plate interferometer. However, this approach is not suitable for evaluating the excitation beam originating from the white-light laser due to its limited coherence length. In lieu of interferometry, the collimation of the excitation beam is gauged by measuring the beam size. A perfectly collimated beam is expected to exhibit minimal dispersion with distance, rendering beam size measurements a practical and reliable method for assessing the collimation of the excitation beam in the isoSTED nanoscope.

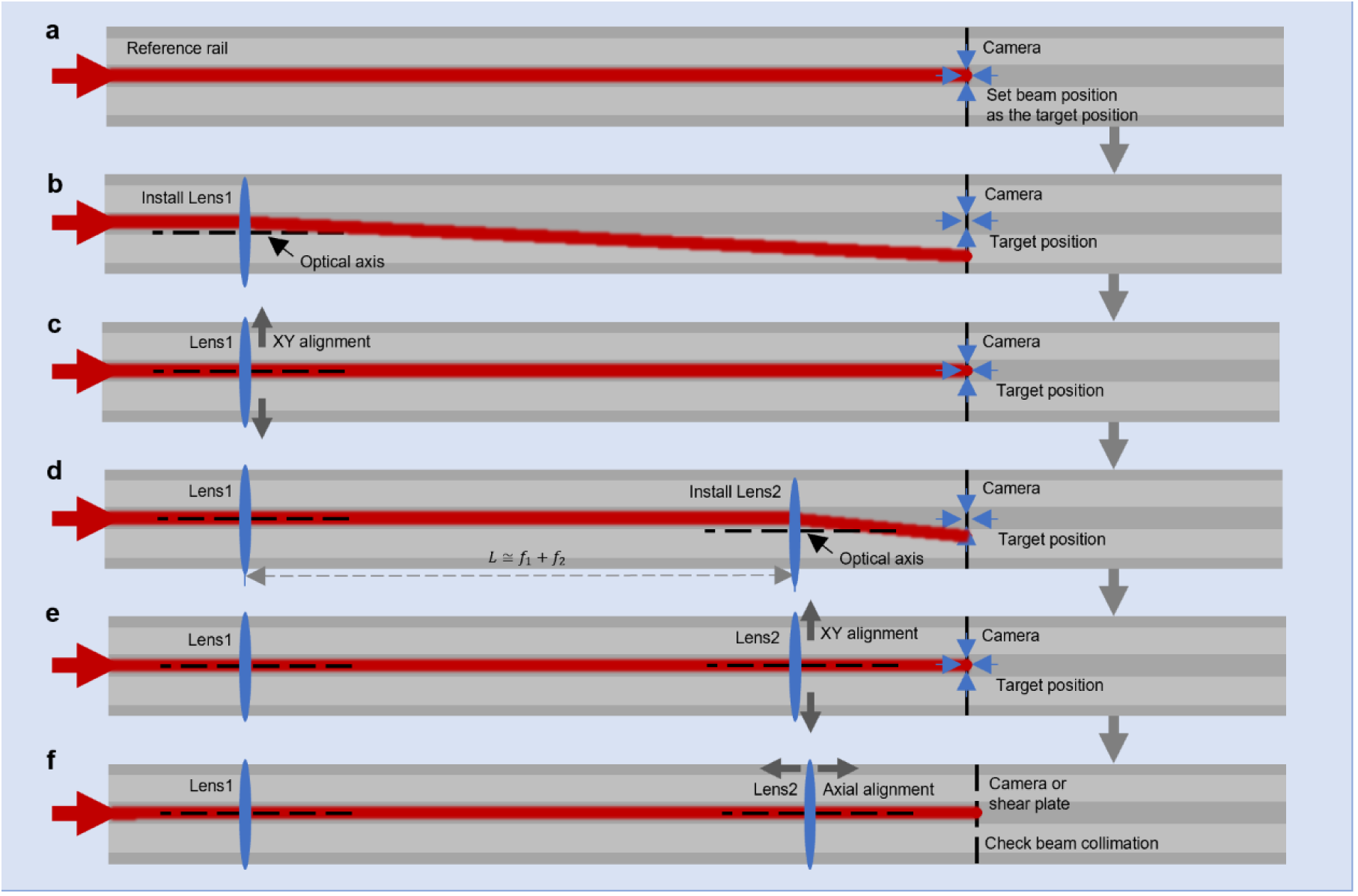

#### AOM and AOTF alignment

The goal for aligning an AOM or AOTF is to optimize its diffraction efficiency, specifically maximizing the power of the output beam in the +1 order. To enhance this efficiency, the incident beam collimation, polarization, and incident beam angle need to be aligned and the frequency and amplitude of the radio frequency (RF) signal need to be adjusted.

#### Polarization separation module alignment

Two beams with orthogonal linear polarization, co-aligned as STED*_xy_* and STED*_z_* depletion beam respectively, are constructed with a single 775-nm pulsed laser. The STED*_xy_* and STED*_z_* beams become circular polarizations at the common focus out of the objectives, as a result, they can potentially interfere between each other so as to destroy the designed depletion pattern. To avoid interference between the STED*_xy_* and STED*_z_* beams at the common focus of the two objectives, a polarization separation module, inspired by the layout of a Michelson interferometer (**Fig. 6**), has been introduced. In this module, one polarization component of the laser undergoes a controlled delay relative to its orthogonally polarized counterpart. The optical path length difference between the two arms of the module is carefully optimized to introduce a critical delay: it must be longer than the coherence length of the laser pulses, thereby avoiding interference between the two beams, and sufficiently small so that both depletion pulses arrive shortly after the excitation pulse to allow for efficient depletion (see Section ‘Pulse gate/delay modules’). This synchronization feature is pivotal as it significantly influences depletion effects and, consequently, resolution. For our system, a suitable construction is when one arm is 12.5 mm longer than the other, corresponding to a 25-mm difference in optical path length or a temporal delay of 0.83 ns. At this difference, interference between the two polarization components is negligible while depletion is still highly efficient.

**Fig. 6.**
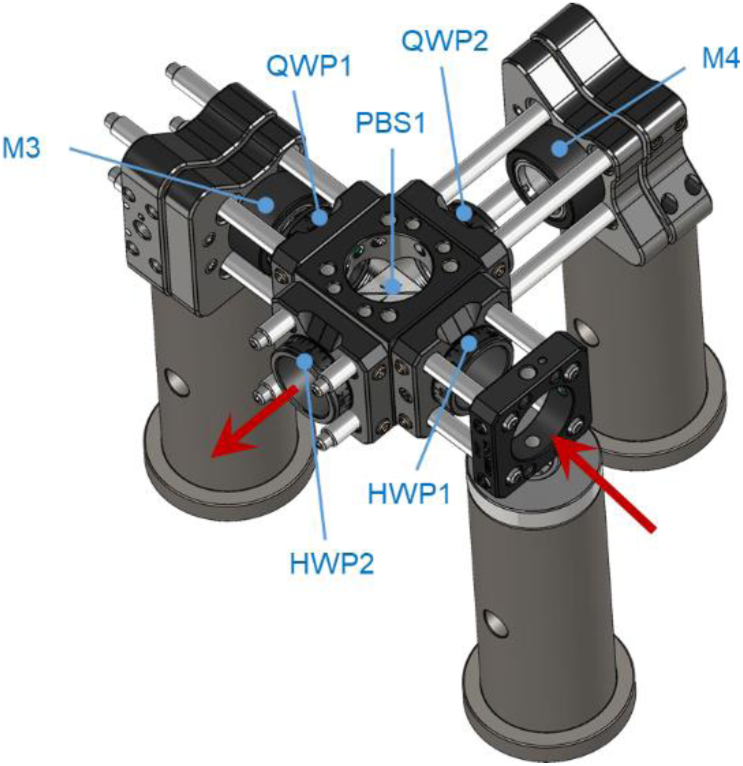
Polarization separation module. This module consists of an optical cage system centered around PBS1. HWP1, HWP2, QWP1, and QWP2 are mounted in rotation mounts. The kinematic mirror mounts can slide along the cage rod, allowing for easy adjustment of the lengths of two arms. The red arrows indicate the propagation direction of the depletion beam.

#### SLM alignment

A single reflective SLM (Hamamatsu, LCOS-SLM X10468-02) is used to generate two adjacent circular holographic grating patterns, which modulate the STED*_xy_* and STED*_z_* beams, respectively. As a polarization-selective device, the SLM exclusively modulates the *p*-polarized beam while the *s*-polarized component remains unaltered. A virtual 4f telecentric system is established by a single lens and a mirror at its focal plane. The mirror reflects the light back into the lens so that the 4f assembly projects the first holographic pattern onto the second one. The depletion beam, coming from the polarization separation module, consists of two incoherent beams with orthogonally linear polarization. Upon the first reflection off the SLM, one of the beams with *p*-polarization undergoes 0-2π vortex phase modulation programmed in the SLM, rendering it into the STED*_xy_* beam. The reflected beams go through the virtual 4f telecentric system and project onto the second holographic pattern. As the beams passing a QWP (QWP3) twice, the polarization of both beams is rotated by 90°. At the second reflection on the SLM, the STED*_xy_* beam is therefore unaffected, while the other, previously unaffected beam experiences the SLM modulation for the axial depletion (STED*_z_*). In the 4Pi-STED mode, the STED*_z_* beam is a normal Gaussian beam profile modulated by a moderate defocus phase pattern to reduce sidelobes in the microscope’s PSF^11^.

Two factors in SLM alignment are critical for achieving high image quality in the isoSTED nanoscope later: (i) The beams need to illuminate the SLM holograms uniformly (**Fig. 7a–c**), as non-uniform illumination causes the focus to tilt along the *z*-axis causing asymmetry of the PSF (**Fig. 7d–f**). (ii) The 4f telecentric system needs to be aligned for accurate image conjugation between two holograms so that both holograms can simultaneously be conjugated to the objective back pupils.

**Fig. 7.**
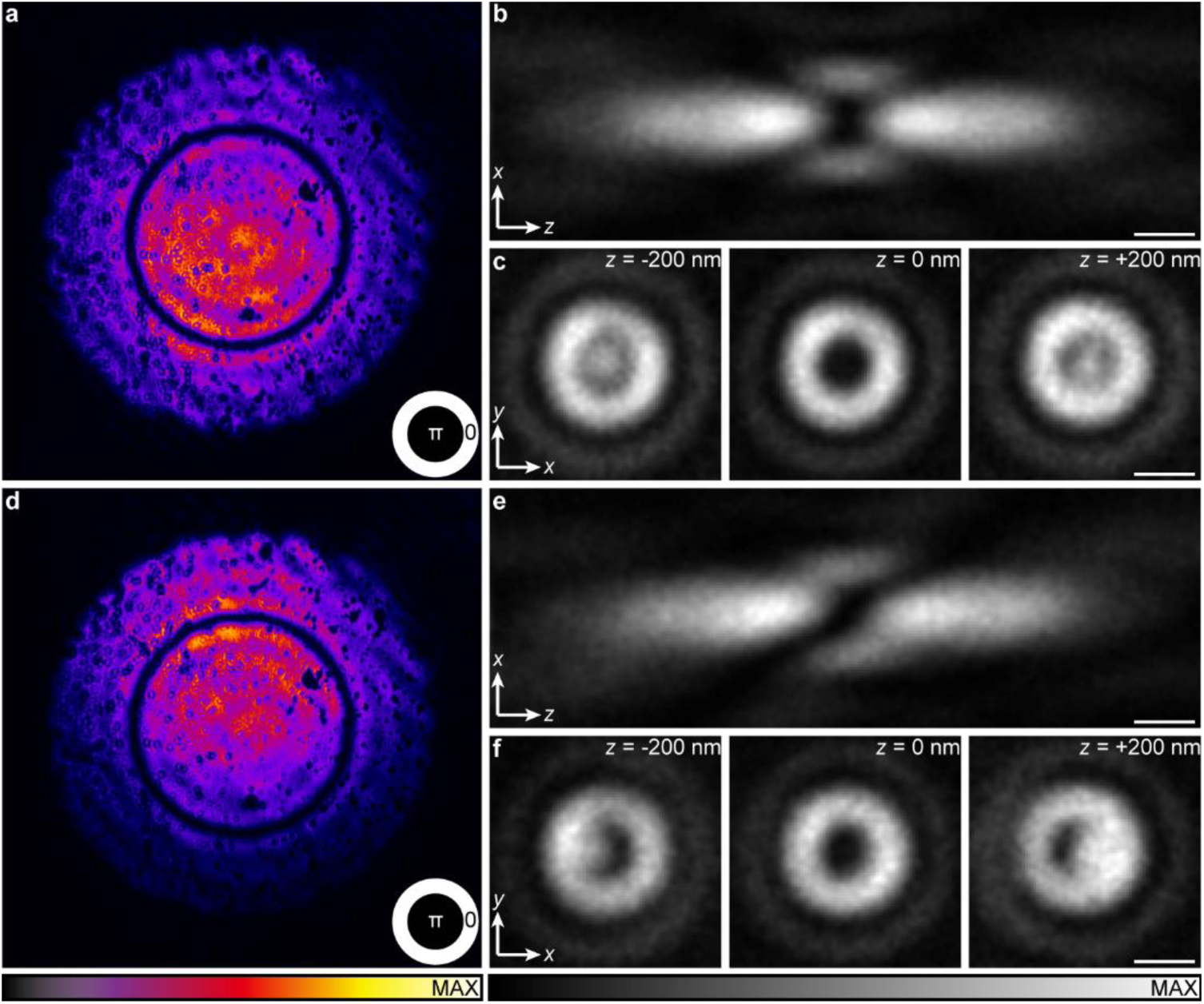
Effects of illumination uniformity on PSFs. **a**, uniform illumination on SLM. The hologram displayed on the SLM is a top-hat shaped 0/π pattern (bottom-right). This specific holographic configuration was selected due to its heightened sensitivity to aberrations, thereby amplifying the distortion within the aberrated PSF. **b**, *xz _y_*_=0_ cross section of PSF under uniform illumination. **c**, cross sections at different depths (*z*) in **b**. **d**, non-uniform illumination on SLM. The hologram added on the SLM is a top-hat shaped 0/π pattern (bottom-right). **e**, *xz _y_*_=0_ cross section of PSF under non-uniform illumination. **f**, cross sections at different depths (*z*) in **e**. Each shown figure is normalized to its own peak intensity. Scale bar: 0.5 μm.

#### Scanner alignment

To ensure uniform illumination across the entire FOV, the scanner is strategically located at an intermediate conjugation plane positioned between the spatial light modulator (SLM) and the back pupils of the objectives. It features one resonant mirror for fast-axis (*x*) scanning, complemented by two galvanometer mirrors operating synchronously to scan along the slow-axis (*y*). The two galvanometer mirrors establish a virtual tilt plane seamlessly coinciding with the resonant scanning plane. Since the resonant mirror is placed in a plane conjugated to the back pupil of the objectives, this arrangement allows for 2D beam tilting in the back pupil of the objectives (**Fig. 8a**). To achieve this, the rotation angles of the first (*α*) and the second galvanometer mirrors (*β*) satisfy the following relationship: *⍺L*_1_ = −*β*(*L*_1_ + *L*_2_), where *L*_1_ and *L*_2_ denote the distance from the resonant mirror to the first galvanometer mirror, and between the first and the second galvanometer mirrors, respectively. The negative sign in the equation indicates that the two galvanometer mirrors rotate in opposite directions during operation. Specifically, when *L*_1_ = *L*_2_, *⍺* = −2*β* (**Supplementary Fig. S2**). Additionally, synchronization between the resonant mirror scanning and laser blanking is important (**Fig. 8b** and **Supplementary Fig. S3–S4**): as the resonant mirror oscillates, the focal point scans along the *x*-axis in a sinusoidal fashion within the focal plane (**Fig. 8c**). To mitigate image distortion, only the relatively linear region in one scan direction, corresponding to 1/3 of the period of the sinusoidal scan motion, is used for imaging. The remaining difference to the ideal linear scan motion would still cause distortions in the image, and a non-uniform sampling period, where pixel dwell times are shorter when the scanner is moving faster, is therefore implemented to correct it. Further, to reduce photobleaching, the synchronization signal from the resonant mirror also serves as the timing reference for laser blanking to turn off the laser illumination when no image data is collected (**Fig. 8b**).

**Fig. 8.**
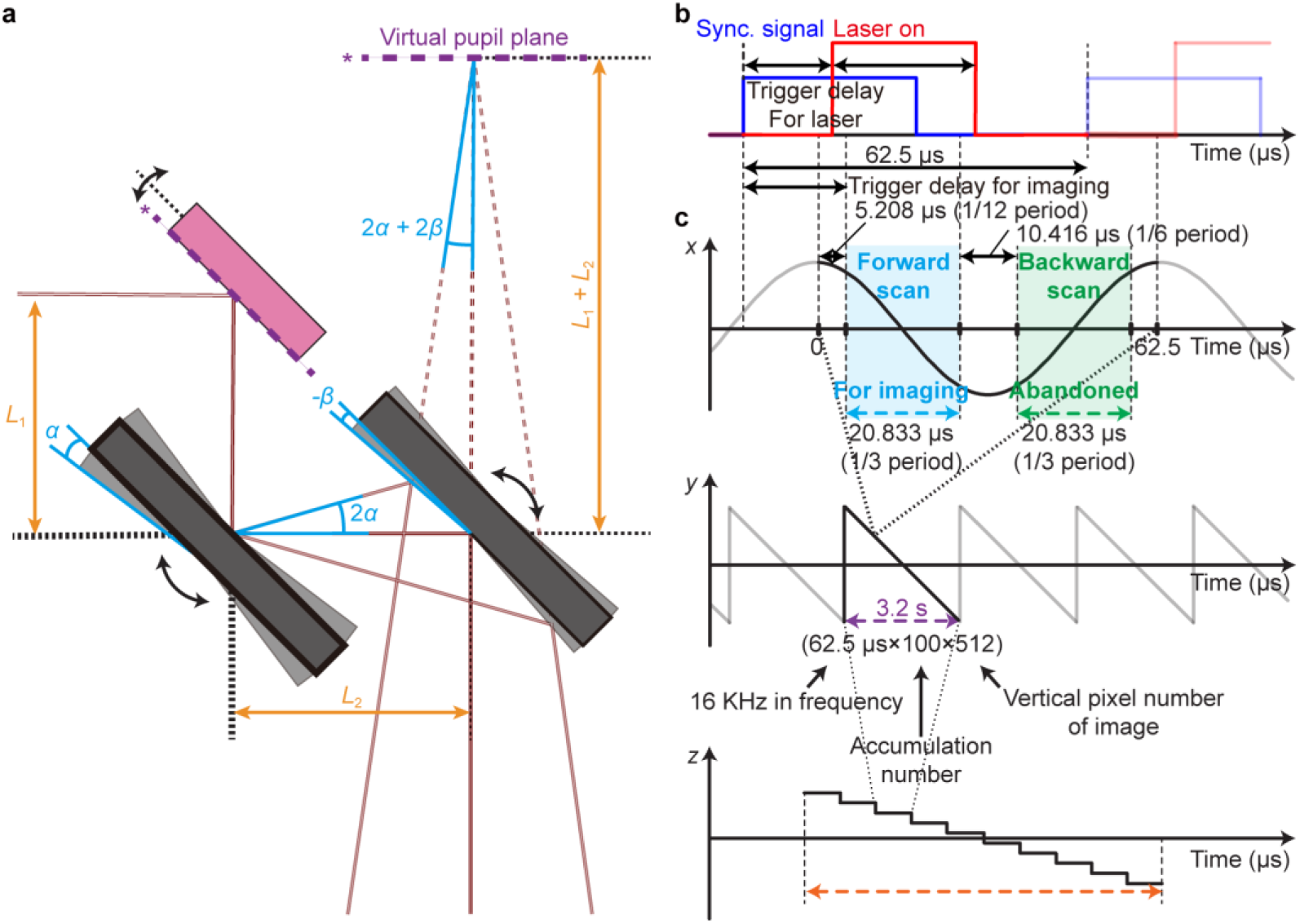
Beam scanning concept of isoSTED nanoscope. **a,** scanning mirror module (including one resonant mirror and two galvanometer mirrors). **b**, Synchronization and laser pulse trigger signals. **c**, Focus position in three dimensions during imaging. From top to bottom: the position of the focus in *x*-, *y*-, and *z*-direction. Please note the different time scales of the axes. In the top row, the amplitude of the cosine function is positively related to the control voltage of the resonant mirror. To avoid image distortion, only the relatively linear region of the motion is used to collect fluorescence signals for imaging. The backward scan motion is not used for imaging. In the middle and the bottom rows, the waveforms of the focus *y*- and *z*-positions are identical to those of the control signals for the galvanometer mirrors and the piezo *z*-axis stage, respectively.

While recording data during the forward scan is straightforward, image formation with scanning in both directions requires reversing the backward scanning in which pixels are recorded. Consequently, specialized read-write buffers must be used to invert the image data. To simplify system synchronization, we have chosen to only collect data during the forward scan and not use the backward scan for imaging.

#### 4Pi interference cavity alignment

In the 4Pi interference cavity setup depicted in **Fig. 2** and **Fig. 4**, a polarizing beam splitter (PBS2) is used to generate a counter-propagating pair of beams. To ensure correct alignment, the galvanometer mirrors and resonant mirror need to be powered on when aligning the post-scanner beam paths, especially the 4Pi interference cavity. Moreover, as at other locations in the isoSTED nanoscope, the setup incorporates two rails (*e.g.,* Thorlabs, RLA450/M, R5 and R6 in **Fig. 4**), installed along the upper and lower arms of the 4Pi interference cavity. These rails serve as fixed references for the direction of the beams, ensuring easier reproducibility of the beam alignment.

Before the PBS2, the HWP3 rotates the polarization of the incoming depletion beam to ±45° relative to the polarization axis of the PBS. This adjustment allows for precise power allocation at a 50/50 ratio between the two cavity arms. Concurrently, the polarization of the excitation beam, already polarized at 45° upon incidenceat HWP3, is rotated to be parallel to the vertical breadboard. Consequently, this beam is exclusively coupled into the upper arm of the 4Pi interference cavity. In the upper arm, another HWP (HWP4) further rotates the polarization of the depletion beams to match those of the lower arm, facilitating subsequent interference at the common focus. To minimize phase delay discrepancies between the two arms, a mirror (M11) and a BK-7 glass wedge is translated within the upper and the lower arm, respectively. Additionally, both arms of the interference cavity have a DM integrated in the plane conjugated to the back pupil of the respective objective. The DM enables aberration correction and manipulation of the PSF, as detailed in the Sections ‘DM alignment’ and ‘Aberration correction’. To ensure circular polarization of all beams at the focus, QWP4 and QWP5 are positioned right in front of the objectives. These components collectively contribute to the precise manipulation and control of beam polarization and phase, essential for optimal performance within the 4Pi interference cavity.

#### DM alignment

The alignment of the DMs includes both axial conjugation and lateral co-centering with designated optical elements. By conjugation with the resonant mirror of the scanner, the DMs in this system are at the same time conjugated to the objective back pupils and the SLM holograms. Designated positions at the end of rails R5 and R6 denoted where the DM surfaces should approximately reside (**Fig. 9a–b**). Mounting the alignment camera at the designed position allows verifying the SLM images before installing and finetuning the DMs. Following offline calibration (see Section ‘SLM and DM calibration’ for details), the two DMs are installed at their designated positions on the vertical optical breadboard, roughly co-centered and conjugated with the SLM holograms. The DM mount provides the capability for axial (*z*) translation as well as lateral (*xy*) translation and tip/tilt angle adjustment, allowing for more precise conjugation and co-centering.

**Fig. 9.**
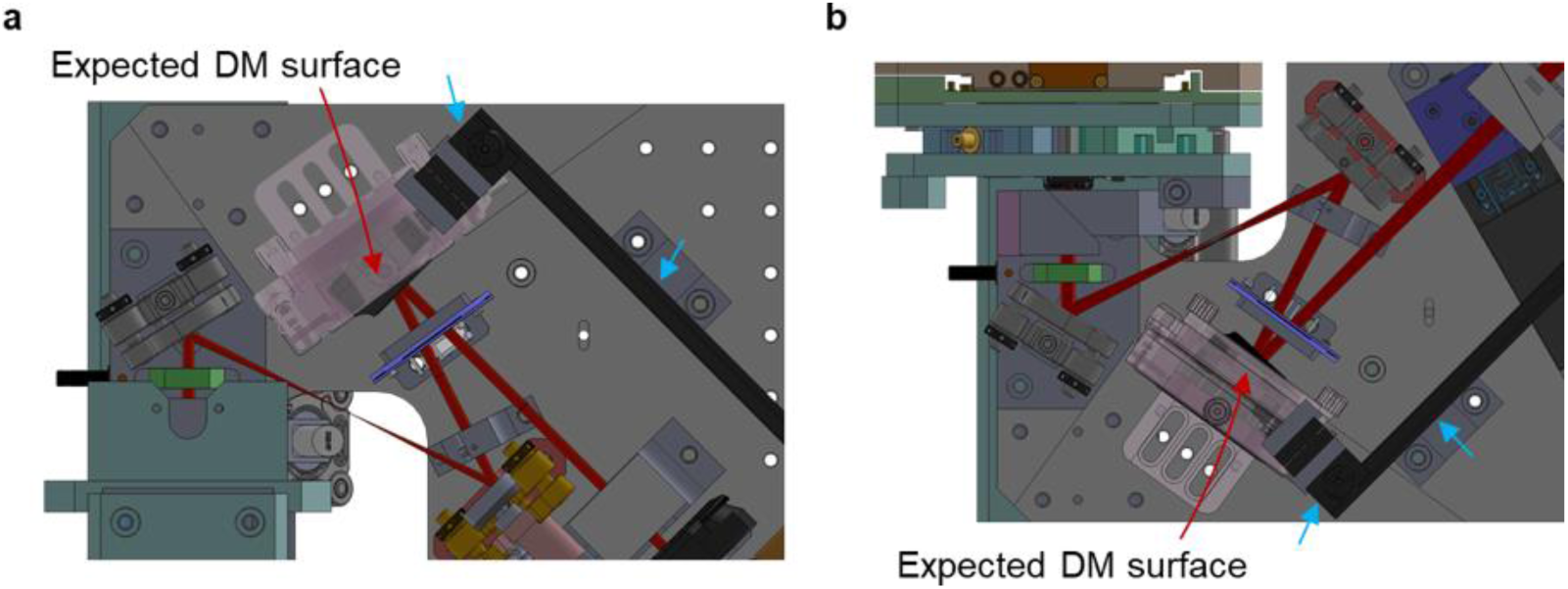
DM conjugation alignment. **a,** upper beam path. **b,** lower beam path. Blue arrows indicate the reference positions for assembly of the alignment camera. The rails R5 and R6 should be installed in accordance with the CAD model, and the end surface of the rail carrier should coincide with the edge of the rails. The alignment camera is placed at the desired DM surface with this assembly.

Each DM is conjugated to the back pupil plane of its corresponding objective through a 4f telecentric system. A successful conjugation is indicated by the absence of grid patterns stemming from DM membrane surface diffraction at the objective’s pupil plane (**Supplementary Fig. S5a–b**). For co-centering, a cross-shaped and a top-hat shaped 0/π phase pattern,are loaded onto the DM and SLM, respectively. The presence of an asymmetric PSF indicates a lack of co-centering alignment (**Supplementary Fig. S5c–d**).

#### Aberration correction

Careful adjustment of the AO elements is crucial to guarantee good imaging performance of the system. The following steps outline the process:

(i). Correction of system aberrations.
(ii). Compensation for sample-induced aberrations.
(iii). Objective alignment.
(iv). Enable focus-lock function (refer to Section ‘Focus-lock module’ for details).
(v). Translation to the region of interest.
(vi). Fine compensation of local sample-induced aberrations.

In our setup, we did not use a wavefront sensor and opted instead for image-based aberration correction to optimize the Zernike or 4Pi coefficients^16^ of the back pupil(s). This approach involves a systematic procedure where a series of images are captured while introducing varying degrees of specific aberration modes via the DMs or SLM. For each set of introduced aberrations, an image is acquired. We then use an image quality metric, such as brightness or contrast, to quantitatively assess the fidelity of each acquired image.

Following previously established procedures^17^, our correction algorithm plots for each specific aberration mode the determined image quality metric values as a function of the applied aberration mode amplitudes. Fitting a (Gaussian) curve to this data, the algorithm then determines the optimal amplitude for this aberration mode that maximizes image quality. This iterative process is completed for each of the aberration (Zernike) modes under consideration.

This approach does not need a complex wavefront sensing apparatus and offers an efficient and tractable means of aberration correction. By leveraging image-based techniques in tandem with systematic optimization algorithms, we achieve targeted aberration correction, thereby enhancing the overall imaging performance of the isoSTED nanoscope.

#### Image matching of upper and lower objectives

This step minimizes the lateral offset between the FOVs of the upper and lower objectives. To achieve the interference that is essential for isoSTED nanoscopy, the laser foci produced by the two objectives must overlap to a high degree. This alignment is achieved by translating the lower objective using the two-axes piezo stage (PI, P-612.2SL). While this procedure is regularly applied during the later routine isoSTED operation, particularly when transitioning to a new sample, the initial setup often shows such strong misalignment between the objectives that it surpasses the travel range of the piezo stage. To address this, a coarse adjustment of the two FOVs is required by adjusting the threaded-rod pushers (Newport, AJS100-0.5H-NL) underneath the piezo stage with a hex key. During this adjustment, two beam spots on the top-right side of PBS2 serve as the reference points. These spots originate from the depletion beams split by PBS2 and passing through the cavity in clockwise and counter-clockwise direction, respectively, can be observed with a camera or directly by eye using a paper screen. Adjustment is considered complete when these two spots align in position.

#### Excitation laser path alignment

When aligning the excitation laser path, it is imperative to carefully adjust it to be colinear with the depletion beam. Additionally critical elements include: (i) a filter, more specifically an AOTF in our isoSTED configuration, that selects the desired excitation wavelengths from the white-light laser beam for multicolor fluorescence excitation; (ii) chromatic dispersion correction for the selected wavelengths within a white-light laser pulse to synchronize excitation with the depletion laser pulses and thereby achieve maximum stimulated depletion for all used excitation windows (refer to Section ‘Chromatic dispersion compensation module’ for details); (iii) optimal beam size magnification to adequately fill the back aperture of the objective, thereby maximizing excitation intensity; and (iv), analogous to the depletion beam, careful attention to the polarization of the excitation beam for appropriate power allocation between the 4Pi interference cavity arms.

#### Detection beam path alignment

The detection light path involves de-scanned collection of the fluorescence through a multi-band dichroic mirror that separates it from the common light path with excitation/depletion lasers. Subsequently, the fluorescence light is split into three spectral windows by additional dichroic mirrors, and each of the three resulting beams is focused by a lens and coupled into a multi-mode fiber for collection by an APD. For the best collection efficiency, each fiber tip is conjugated to the common focus of the objectives in 3D, and the focused fluorescence is coupled to the fiber to the maximal efficiency.

#### Data analysis

In contrast to SMLM or SIM, isoSTED nanoscopy does not need post-processing for image reconstruction. Nevertheless, data analysis is still of the essence for resolution quantification. In instances where the sample exhibits sparse, small structural features, such as when imaging fluorescent beads, the resolution is assessed from the spot size by fitting a Lorentzian model function to the data (code available at https://zenodo.org/uploads/10574469). We caution against directly extrapolating system resolution from biological samples using Lorentzian fitting, as the intricate nature of such specimens may dramatically compromise accuracy. Instead, for a more comprehensive evaluation of attained resolution in images, Fourier ring correlation (FRC) methods^18^ or Nested Loop Fitting^19^ prove to be viable analytical approaches.

#### FPGA Control

Signal generation and data collection is centered around two NI FPGA DAQ boards housed in a single PC running the LabVIEW software environment in Microsoft Windows. Specifically, an NI PCIe-7852R FPGA board is used for data collection and overall synchronization (sync), while an NI PCIe-7841R FPGA board is used for waveform generation and laser power control and operates in response to triggering from the aforementioned 7852R FPGA board. This necessitates using a PC with a minimum of two PCIe slots on the motherboard (refer to Section ‘Computer and data storage’ and ‘Nanoscope control computer’ for specific requirements on computer). An overview of the key hardware connections for image acquisition is presented in **Fig. 10**. The two FPGA boards compose the core of the image collection hardware and are physically linked (internally with a Real-Time System Integration (RTSI) cable) to share trigger signals. The 7852R FPGA board coordinates data collection from the APDs for light detection with the scan motion of the 16-kHz resonant scanning mirror. The synchronization signal from the 16-kHz resonant mirror triggers waveform generation from the 7841R FPGA board for controlling galvanometer mirrors, as well as AOM and AOTF for the depletion and excitation beam blanking, respectively. Although these two FPGA boards sit inside the control PC, once running, they operate autonomously by executing code that does not rely on any timing or input information from the PC. The high data collection speed required to work with a 16-kHz mirror prevent the use of software-timed data acquisition or on-the-fly adjustments. The 7852R FPGA board collects photon time traces from each APD (color channel) with a constant time interval. Because the resonant scanning mirror moves sinusoidally, the sampling in space varies. To compensate for this effect, the time trace is processed by re-binning it into evenly spaced pixels (linearization) in space (rather than time). This process happens in real-time on the 7852R FPGA board. Furthermore, the speed of the 16-kHz resonant mirror results in pixel dwell times on the order of tens of nanoseconds and in a single scan motion of the resonant mirror very few photons are collected. Scanning a single line multiple times and accumulating the recorded signal is therefore necessary to achieve reasonable photon numbers. This accumulation happens in real-time on the 7852R FPGA board. After a single line has been scanned a user-set number of times, the accumulated signal is sent by the 7852R FPGA board to the PC where it is concatenated by the LabVIEW program to the current image and displayed. The process continues until either the live image capture is stopped by the user, or the image capture ends after a set number of frames.

**Fig. 10.**
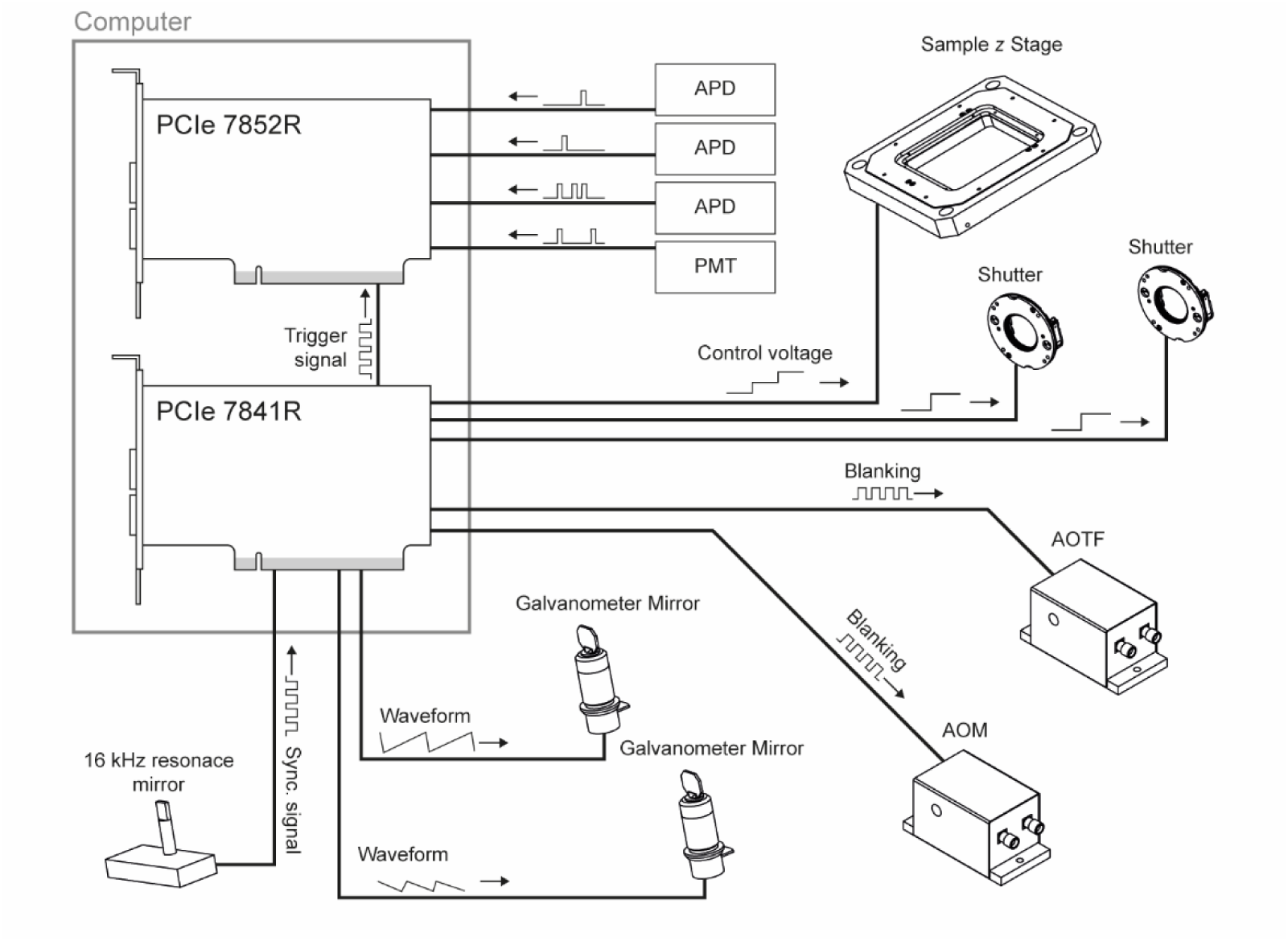
FPGA-based DAQ key connections. Two FPGA-based DAQ boards act as the core of the instrument control and image acquisition hardware. Key connections for image acquisition are presented here. Briefly, the 16-kHz resonance mirror sync signal acts as the master clock for image acquisition and is routed into the 7841R FPGA board where it triggers waveform output to the two galvanometer mirrors and is used as the reference signal for blanking the excitation and depletion beams. The 7841R board internally passes the 16-kHz sync signal to the 7852R board, which triggers data collection from the user-selected APDs or the PMT. The 7841R FPGA board also controls the sample piezo z-position and opens and closes shutters in the cavity beam path.

#### Synchronization and timing

Several carefully timed events must be coordinated for successful isoSTED imaging. They are synchronized to two master timing signals that run independently but in combination govern STED data collection. First, the 16-kHz resonant mirror acts as the master clock for image data collection. This signal regulates when the FPGA boards check for photon arrivals from the APDs/PMT, blank the AOM/AOTF, and trigger waveform generation for galvanometer mirror control. Second, the white-light laser (∼78-MHz repetition rate) acts as the master clock for laser pulse synchronization and detector gating. The electronic sync signal from the white-light laser triggers pulse generation for the depletion laser and triggers the detector gating boards for gated photon detection.

For image data collection, rising edges from the APDs, each representing a detected photon, must be registered to the current scan position of the resonant mirror scanning a line of the sample. The 16-kHz resonant mirror synchronization output transitions between high (+5 V) and low (0 V) each time the mirror changes direction. When the 7841R board detects a rising edge from the resonant mirror synchronization signal, marking the start of a new line scan in the forward direction (**Fig. 8**), the signal is passed to the 7852R FPGA board. Once detected this rising edge signal, the 7852R FPGA board waits for 5.208 µs before a ∼20-µs photon time course with 5-ns resolution from each APD is collected (**Fig. 10**), corresponding to the data collection only from the central ∼1/3 portion of the period in the scan motion for image formation. This limitation ensures that data is only collected over the relatively linear portion of the mirror scan. During the data collection time course of the 7852R FPGA board, it records the time of each APD output rising edge. Each APD output is checked every 5 ns. The deviation of the scanner motion from an ideal linear scan pattern results in the regular 5-ns temporal binning corresponding to an unevenly binned sampling pattern in space. To correct this, the temporal bins of the photon time course are re-binned to create evenly spaced pixels in sample space. Detector gating and laser synchronization are set independently from photon detection using dedicated electronics described below.

For optimal STED resolution, the depletion and excitation laser pulses must arrive in close (<<1 ns) succession at the sample. In the system presented here, the white-light laser cannot be externally triggered and, therefore, has to act as the master (laser) clock. The electronic trigger output signal from the built-in photodiode of the white-light laser is converted (Pulse Research Labs, PRL-350TTL-NIM) and split (Pulse Research Labs, PRL-350TTL-NIM with PRL-SC-104A) into three lines to act as the input, or reference signal, for three custom pulse gate/delay modules designed and built by Opsero Electronic Design. Each pulse gate/delay module has two inputs: one for the white-light laser reference signal and the other for an APD detector signal. The modules are used for two separate functions: they can (i) introduce a user-defined delay between a trigger input and a trigger output pulse, and (ii) they can be used to pass a connected APD signal only for a user-defined time window after a trigger input. For the first function, the pulse gate/delay modules can electronically delay the input signal from the white-light laser up to 20 ns in 10-ps steps. The delayed trigger signal is passed to the depletion laser via a signal translator (Pulse Research Labs, PRL-420ND), allowing their pulse arrival time at the sample to be matched with the white-light laser pulse’s arrival. In practice, the timing between the depletion and excitation lasers is optimized by continuous imaging of beads until the best resolution (if a vortex hologram has been applied to the SLM) or nearly complete depletion (if the SLM is only displaying a flat and gratings) has been achieved. The second function of the pulse gate/delay modules allows them additionally to be used for gating the signal from the APDs. Specifically, when an electronic pulse is detected, the pulse gate/delay module waits for a programmable time period (the ‘delay’) before allowing the APD signal to pass to the output and on to the connected 7852R FPGA board. The delay between the white-light sync pulse and the start of the detection window can be adjusted between 0.1 ns and 9.8 ns in 10-ps steps. Similarly, the detection window width can also be adjusted within the same range with the same precision. For window adjustment purposes, the detection window width can be set to the minimal value, and the delay relative to the excitation pulse is scanned until the strongest fluorescence signal is observed. This synchronizes the detection window with the excitation pulse. The window delay is then adjusted to be about 1 ns after the excitation pulse to allow the depletion pulse to act before fluorescence is detected for the best resolution, and the window width is opened to several ns to cover the fluorescence lifetime for a full detection.

### 1.5. Experimental design

#### Personnel

At least one full-time experienced microscope developer will need ∼12 months to build an isoSTED using this protocol.

#### Purchasing and delivery

We highly recommend purchasing parts far ahead of time. It takes in our experience 1-2 months to collect quotations and place orders. Some critical system components require up to six months, in extreme cases even longer, to be delivered after placing purchase orders and should therefore be ordered as early as possible. These parts include in our experience the deformable mirrors (Boston Micromachines), actuator stages (PI and ASI), the SLM (Hamamatsu), lasers (NKT Photonics and MPB Communications Inc.), FPGA boards (NI), and objectives (Olympus).

The pair of objectives used on the system needs to match magnification within 0.3% to ensure sufficient image overlap throughout the FOV. Therefore, we strongly recommend asking the vendor to provide a pair of magnification-matched objectives. After delivery, we recommend imaging a Ronchi Ruling with micrometer-scale for a magnification comparison. The same region should be measured using each objective on a widefield microscope stand. Fourier analysis can be utilized to measure the average distance between the line spacings.

#### Optical table and device setup

A minimum size of 1×1.5 m^2^ is required for the optical table. The optical table top should have a honeycomb core structure and dampers (*e.g.*, TMC, 784-SPECIAL). The table legs should be equipped with automatic leveling vibration isolators (*e.g.*, TMC, 14NP-417-89). After the optical table has been installed, verify its leveling and check that the vibration-isolation legs are operating properly by pushing the optical table top up and down to confirm an adequate range of motion. This quick verification process should be repeated on a daily to weekly basis, especially during system installation, since some vibration isolators require in our experience regular tuning.

Since the nanoscope presented here includes many devices with controllers and power supplies, we recommend installing an overhead shelf or a 19-inch rack for the controllers and power supplies, rather than placing them on the optical table. Devices featuring fans should under no circumstances be placed on the optical table during the final phases of system installation and later operation and cables connecting these devices to the optical instrument need to be attached firmly to the table surface or other heavy, rigid structures to isolate the optical instrument from vibrations emerging from the vibration sources.

#### Laboratory recommendations

To achieve the best performance, it is required to install the isoSTED nanoscope in a dark, temperature-controlled (temperature stable within ±0.5 °C over extended periods of time, including nights and weekends), and low-vibration (VC-C or better) laboratory. Thus, we recommend a laboratory space without windows located on the ground level or basement. In addition, an isoSTED nanoscope requires minimal environmental vibrations to preserve the axial resolution enhanced by light interference, and thus the laboratory space must be located away from heavy machinery and foot traffic. We strongly recommend that builders carefully consider air conditioning systems to prevent direct airflow onto the system, as this can also cause fluctuations in the interference pattern. The laboratory space should be large enough for the optical table, a computer desk, a chair, and either a 19-inch rack system or an overhead shelf for placing controllers and power supplies along with sufficient space for easy personnel access from at least three sides of the optical table. We recommend having an Uninterruptible Power Supply (UPS) system as a backup power option for unexpected power outages.

#### Machine shop selection

The system presented in this protocol includes many custom-made parts that require a tolerance of ±0.1 mm, an angular tolerance of ±0.5°, a surface finish of 1.6 μm, and anodization. We recommend choosing a nearby machine shop with quick turn-around times since in our experience some errors are inevitable and faster return of revised parts will save time.

#### Alignment dowel pins

Alignment dowel pins are used to assemble the custom parts in their designated positions. Using the dowel pins ensures that parts can be positioned precisely and reproducibly. Thereby, the alignment dowel pins should fit with a small tolerance; for example, the diameter of holes should be between 4.025 mm and 4.050 mm for a 4-mm-diameter dowel pin. In many cases, the diameters of holes are reduced after anodization. Therefore, we recommend that builders use a hand-reamer to enlarge the holes as needed. Dowel pins should be selected to be longer than the depth of the hole (manually confirm the hole depth before inserting any dowel pin!) to prevent pins from disappearing in a hole which makes it extremely difficult to retrieve them.

#### Laser safety

Setting up and using an isoSTED nanoscope as described in this protocol requires careful consideration of laser safety. This system utilizes high-power lasers (>3 W), posing a significant safety hazard. In particular, the 775-nm depletion laser can appear dim due to its long wavelength to the human eye. This may lead to an underestimation of the associated exposure risks. Therefore, collaboration with local laser safety personnel is essential during system construction and operation, and adherence to established safety protocols is paramount. Additionally, maintaining a fully enclosed system and ensuring laser interlocks on either the enclosure or laboratory door is crucial to prevent accidental exposure. During setup and alignment, extra precautions must be taken due to temporarily open beam paths.

#### Alignment camera

In the alignment procedure, the beam needs to be frequently inspected at specific locations of the beam path. For instance, to confirm the conjugation and co-centering of the SLM and a DM, reflections from both device surfaces need to be imaged in the position of the corresponding objective’s back pupil plane. While these tasks can be in general executed by eye with a screen made, for example, of paper, using an inexpensive camera is highly advantageous as it allows to magnify, quantify and record the beam profile at the inspected location.

Many complementary metal-oxide-semiconductor (CMOS) monochrome universal serial bus (USB) camera models can be used as an alignment camera. The main criterion is that the camera should be compact enough to fit into the optical path without colliding with existing optical elements. Additionally, the position of the sensor within the camera body should be known so that the sensor can be placed at the correct location dictated by the optical design of the instrument’s beam path.

In this protocol, commercially available cameras from IDS Imaging were used, obtained either from Thorlabs (part number: DCC1545M) or Edmund Optics (part number: 84-933). To minimize the camera size, we replaced the camera sensor housing by a custom-designed mount which ensures the mechanical centering of the sensor and establishes a known distance between the sensor surface and the mounting flange. The CAD file of this mount is available in the repository at https://zenodo.org/uploads/10574469.

Camera control software programs from IDS Imaging and Thorlabs are both fully functional and compatible with the camera. Software development kits (SDKs) for creating custom applications are also available from both companies. However, when developing our LabVIEW-based nanoscope control software, the ‘uEye’ SDK from IDS Imaging was preferred, as the ‘ThorCam’ package from Thorlabs conflicted with other hardware drivers used in the nanoscope.

#### Nanoscope enclosure

The enclosure surrounding the isoSTED nanoscope is an integral part of the system, serving several crucial functions. It effectively blocks ambient light. Moreover, it plays a vital role in eliminating the impact of air fluctuations on the system’s stability. Additionally, it is thermally insulated and accommodates heating pads and temperature sensors that can be used optionally to maintain the entire system at a constant temperature. This stabilizes the system thermally, thereby minimizing thermal drift, and makes the system, in principle, compatible with live-cell imaging by allowing heating it to 37 °C.

This protocol does not provide a specific design for the nanoscope enclosure. An appropriate enclosure will depend on the dimensions of the optical table and the arrangement of the nanoscope body, electronic cables, and drivers on or surrounding the table. When designing an enclosure, it is essential to avoid mechanical resonances that could exacerbate air vibrations.

#### Computer and data storage

A single computer is sufficient for instrument control, data collection and processing. A typical 3D stack is less than 100 MB in size and a typical experiment generates 1– 5 GB data. Two solid state drives (SSDs) should be installed for fast data write/read access, for example a ‘Samsung SSD 970 EVO 1TB’ drive for the Microsoft Windows Operating System and software installation, and a ‘WDC WDS200T2B0A-00SM50’ 2-TB SSD for data storage. A unique feature is that two FPGA boards for fast data acquisition and device control are required and need to be connected to the PCIe slots on the motherboard (ASUSTeK, WS X299 SAGE). A 14-Port USB hub is connected to the computer’s internal USB port to expand the device connections. A 12-core Intel(R) Core(TM) i7-7800X CPU at 3.5 GHz is installed as the computer’s central processing unit. A video graphic card (NVIDIA, GeForce GTX 1060 6G) is installed for graphics processing.

#### FPGA

The control architecture for the entire nanoscope system revolves around two FPGA boards (**Fig. 11**). They are connected to the PC via PCIe slots on the motherboard. **Supplementary Table 2** lists all active connectors on each FPGA board in our nanoscope, along with their corresponding functions. In brief, the analog signals from the first FPGA board (NI, PCIe-7841R) are used to actuate the piezo stages, scan the galvanometer and resonant mirrors, and adjust the laser beam intensity by modulating the transmittance of the AOM and AOTF. Digital outputs from the same FPGA board are used for shuttering and AOM/AOTF blanking. These devices also provide status monitoring signals as feedback to this FPGA board. The second FPGA board (NI, PCIe-7852R) receives and counts signal pulses from the APDs and transmits the count numbers to the nanoscope control software, which generates the fluorescence image. The two FPGA boards are synchronized via a real-time system integration (RTSI) bus.

**Fig. 11.**
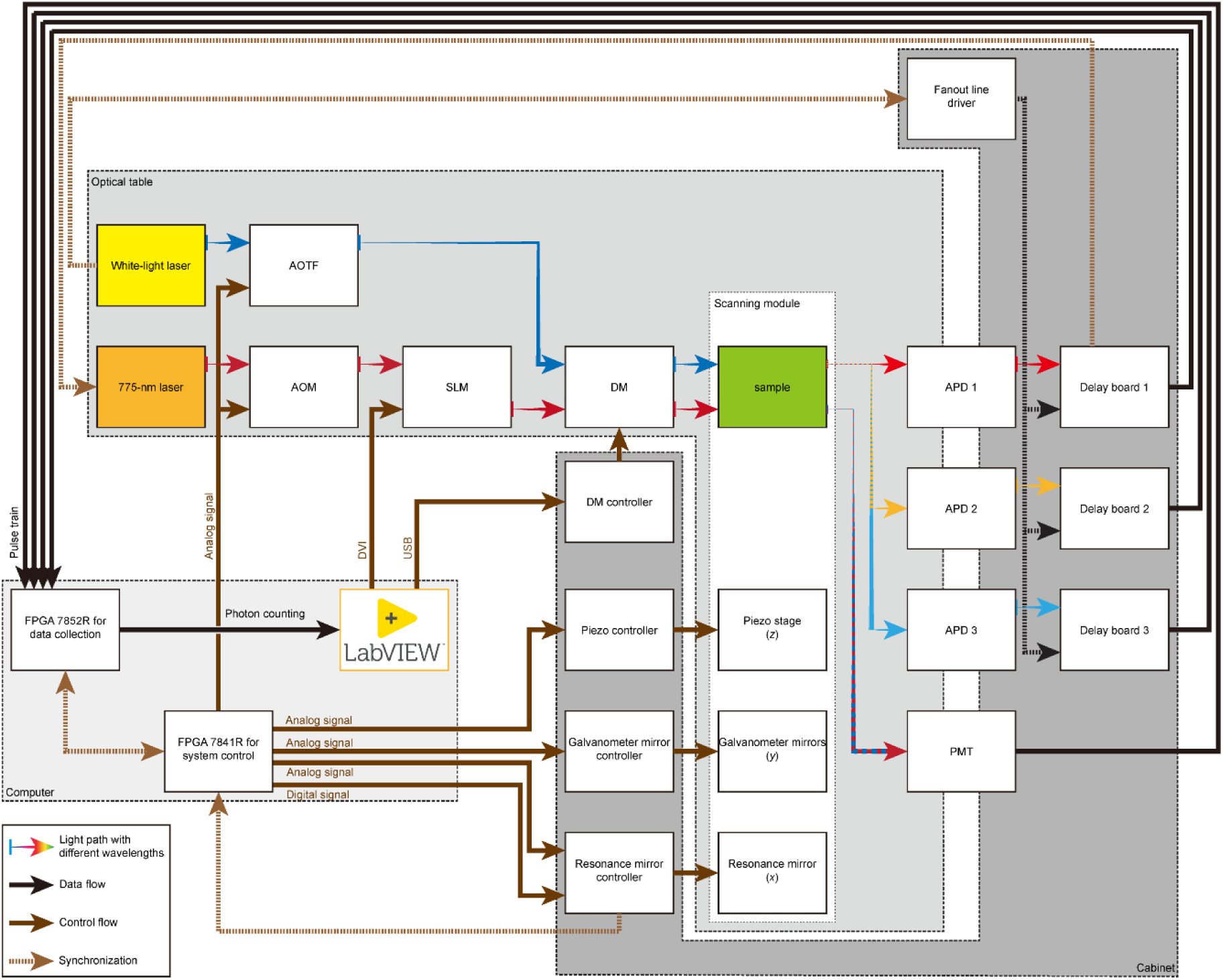
Simplified data flow, control flow, and light path. Abbreviations: AOM, acousto-optic modulator; AOTF, acousto-optic tunable filter; DM, deformable mirror; DVI, digital visual interface; SLM, spatial light modulator; USB, universal serial bus.

#### Pulse gate/delay modules

The application mechanism of the pulse gate/delay module is detailed in Section ‘**Synchronization and timing**’ and recapitulated in **Fig. 12a–c**. To achieve optimal performance, it is critical to optimize the three parameters in the LabVIEW control software, *i.e.,* ‘window width’, ‘window delay’, and ‘laser delay’. The optimal setting is characterized by imaging a fluorescent bead sample with a single objective. First, the optimal ‘window width’ is set at approximately the spontaneous fluorescence lifetime (a few nanoseconds) to collect the most fluorescence signal without adding excessive noise. Second, the optimal detection ‘window delay’ for STED imaging is set ∼1 ns after where the maximal fluorescence detection is achieved with the depletion laser off (**Fig. 12d)**. The additional delay avoids detection of ‘unwanted’ fluorescence from the peripheral excitation volume before the stimulated emission is completed. Last, one of the pulse gate/delay modules is used to activate the depletion laser at a user-defined delay. The optimal ‘laser delay’ is set at the minimized fluorescence, indicating the most effective fluorescence depletion, when the depletion PSF is switched to a Gaussian pattern (**Fig. 12e)**. Once all the three parameters are optimized, only the fluorescence from the fluorophores within the central region of the depletion focal spot is detected, leading to improved resolution and image contrast (**Supplementary Fig. S6**).

**Fig. 12.**
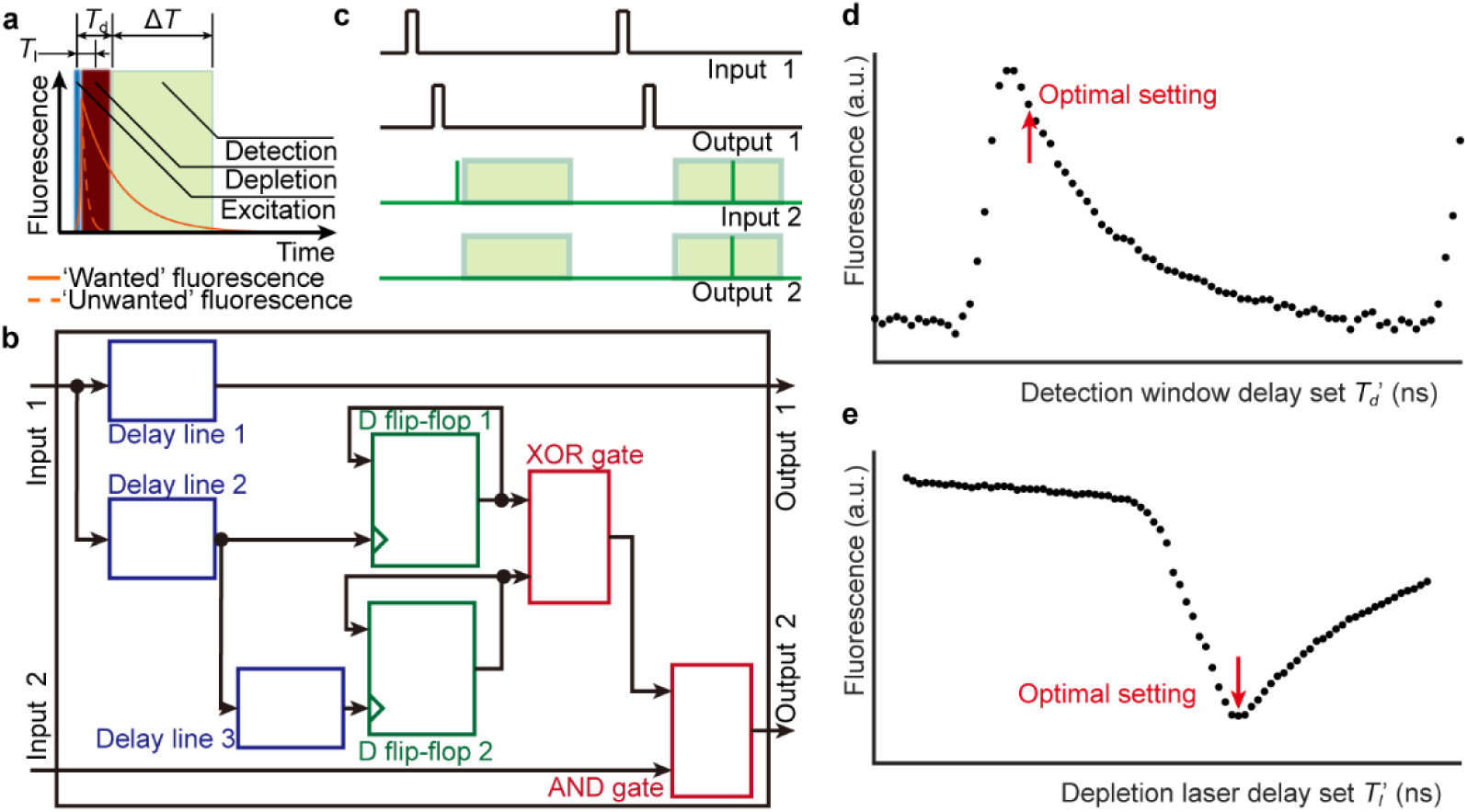
Time-gated detection. **a**, Principle of time-gated detection in isoSTED nanoscope. This diagram shows the time-correlated fluorescence photon counting on the detector under the time sequence of excitation pulse, depletion pulse, and signal detection. The time-gated detection is characterized by the depletion laser delay (*T*_l_), the detection window delay (*T*_d_), and the width of the detection window (Δ*T*). At the center of the ring-shaped depletion PSF, the intensity is zero, and the fluorophore lifetime remains unchanged (‘wanted’ fluorescence, orange line). At positions which coincide with the intensity maxima of the depletion PSF, the fluorophore lifetime is dramatically reduced (‘unwanted’ fluorescence, orange dashed line). By recording the arrival time of photons when imaging it is thus possible to increase image resolution by constructing the final image only from photons which arrive after a certain time; *i.e.* early photons from the periphery of the effective PSF are excluded from recording. **b**, Internal logic of the circuitry (namely, ‘pulse gate/delay module’) used to implement the time-gated detection. **c**, Pulse trains at each connector on pulse gate/delay module. **d**, Typical fluorescence photon counting as a function of the detection window delay setting (*T*_d_’) on pulse gate/delay module. The optimized fluorescence collection for STED imaging is set about 1 ns after the peak (highlighted by a red arrow). **e**, Typical fluorescence photon counting as a function of the depletion laser delay setting (*T*_l_’) on pulse gate/delay module. To quantify the depletion effect, the depletion PSF is switched to a Gaussian pattern. The minimal fluorescence (highlighted by a red arrow) indicates the optimal delay set for depletion laser. The optimal values of *T*_d_′ and *T*_l_′ need to be calibrated for each system.

#### Refractive index considerations

To guarantee optimal resolution across a depth of up to tens of microns within the sample, it is imperative to minimize the refractive index (*n*) mismatch between the sample, its mounting medium, and the objective immersion medium. The isoSTED nanoscope uses two silicon-oil immersion objectives (*n* = 1.406^20^) which provide a good compromise for a large variety of samples ranging from fluorescent beads and cell and tissue samples in ProLong Diamond antifade mountant (Invitrogen, *n* = 1.47 under coverslip^21^) to potential live-cell applications (n = 1.37–1.39)^22^. Additionally, the deformable mirrors in the 4Pi interference cavity compensate for residual aberrations.

#### Multi-color imaging

Multi-color STED imaging is achieved with three excitation beams, which have wavelengths of 488 nm, 590 nm and 650 nm, respectively, and one depletion beam at 775 nm. To separate the illumination and fluorescence detection beam paths, a multi-band dichroic mirror is used (**Fig. 13a**). In the detection path, two long-pass dichroic mirrors separate three channels (**Fig. 13b**). Generally, dyes with well-separated spectra are used, for example, ATTO647N or STAR635P for the red channel and ATTO594 for the orange channel. In addition, three-color STED imaging can be achieved using a dye with wide emission spectrum (*e.g.,* STAR 520SXP) because its fluorescent signal can be detected simultaneously by the two/three APDs (see **Supplementary Fig. S7** for details) and depleted by the same 775-nm laser.

**Fig. 13.**
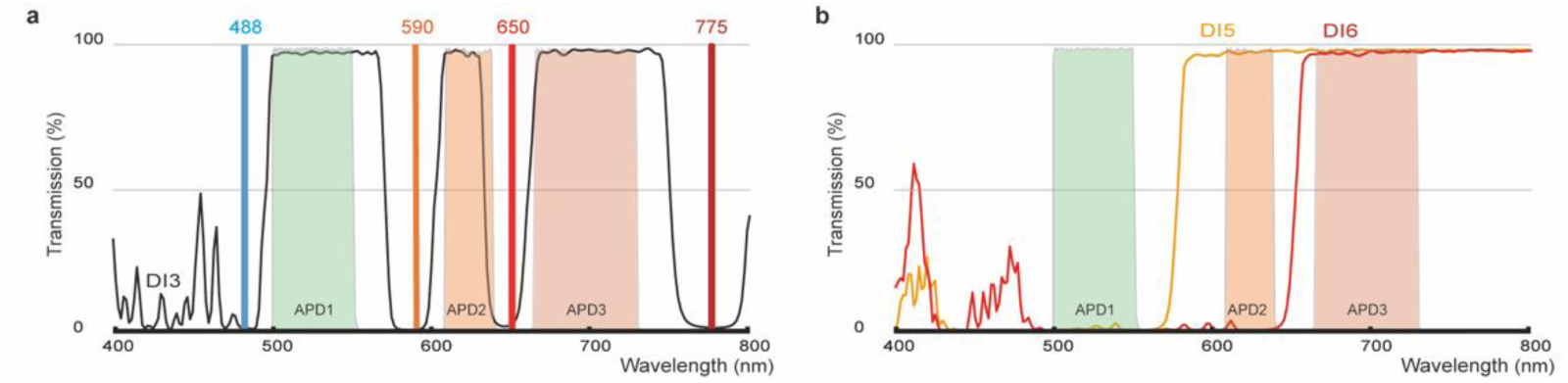
Spectrum configuration for multi-color imaging. **a**, Transmission spectrum of DI2 (black solid line), laser wavelengths (vertical lines labeled with 488, 590, 650, and 775). **b**, Transmission spectra of DI5 (orange solid line) and DI6 (red solid line) in the detection path. Both figures show additionally the three emission filter spectra of the APDs as shaded areas.

#### Focus-lock module

In isoSTED nanoscopy, optimal resolution can only be achieved when the two opposing objectives are consistently aligned to a shared focal point termed the ‘focal zero.’ Maintaining objective alignment during imaging is critical and requires lateral (*xy*) and axial (*z*) synchronization. When the PSFs fail to overlap laterally, the interference pattern that is necessary for this technique cannot be created. Axial misalignment of the objectives results in a shift/distortion of the interference fringes, thereby degrading the imaging quality.

To maintain objective alignment within ±5 nm tolerance during imaging sessions, our nanoscope incorporates a focus-lock module (**Fig. 14**). Within this setup, a continuous-wave laser operating at 980 nm is coupled into the beam path of the upper arm of the 4Pi interference cavity by a dichroic mirror (DI3). The laser beam is then focused by the upper objective and collected again by the lower objective. Subsequently, a second dichroic mirror (DI4) separates the beam from the imaging path. The focus-lock laser beam passes a 400-mm-focal-length lens followed by a weak cylindrical lens to introduce astigmatism before being projected onto a camera to observe the resulting cross-shaped astigmatic focal spot. Throughout the imaging process, both the position and the shape, more specifically the aspect ratio, of the resulting focal spot are recorded and monitored. The spot’s positional changes indicate the relative *x* and *y* alignment of the objectives, while changes in its aspect ratio indicate drift in the *z*-direction. When changes are detected, the piezo objective stages are translated accordingly for compensation using a proportional-integral-derivative feedback control loop programmed in LabVIEW.

**Fig. 14.**
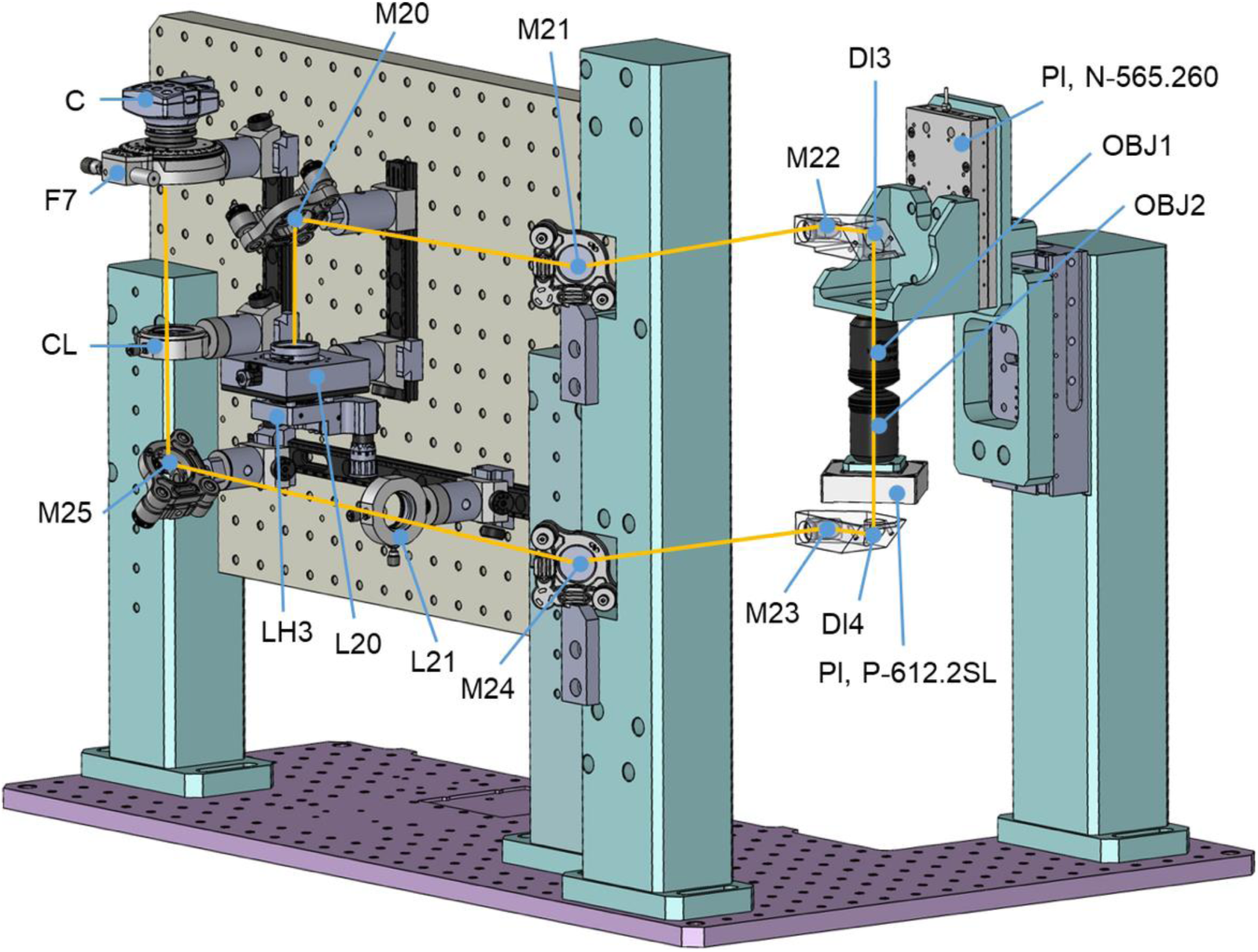
Overview of focus-lock module. A NIR laser (LH3) is launched and collimated by L20. The beam is sequentially reflected by M20, M21, M22, and DI3. The beam is then focused and collimated by OBJ1 and OBJ2, respectively, and directed to the back panel by DI4, M23, and M24. L21 focuses the beam on a camera (C) via M25 and a filter (F7), while CL deliberately introduces astigmatism into it.

This focus-lock module facilitates prolonged imaging sessions exceeding three hours. The alignment laser has negligible impact on the sample or image thanks to its low laser power (less than 5 μW) and its 980-nm wavelength being far longer than the laser and emission spectra used for sample imaging.

#### Chromatic dispersion compensation module

In the context of white-light laser pulses, chromatic dispersion describes the phenomenon that the light at different wavelengths within the pulse spreads out in time as the pulse propagates. It originates from the wavelength-dependent refractive index of various optical elements in the beam path, which causes the light at different wavelengths to travel at different speeds. In our isoSTED nanoscope, the temporal mismatch between pulses of different excitation wavelengths is primarily caused by the white-light laser itself and can be as long as several hundred picoseconds (**Fig. 15a**). Because the maximum depletion effect is only achieved when an excitation pulse is immediately followed by the depletion pulse illuminating the sample, dispersion needs to be compensated to match the delays between the depletion laser pulse and the excitation laser pulses for the three different color channels.

**Fig. 15.**
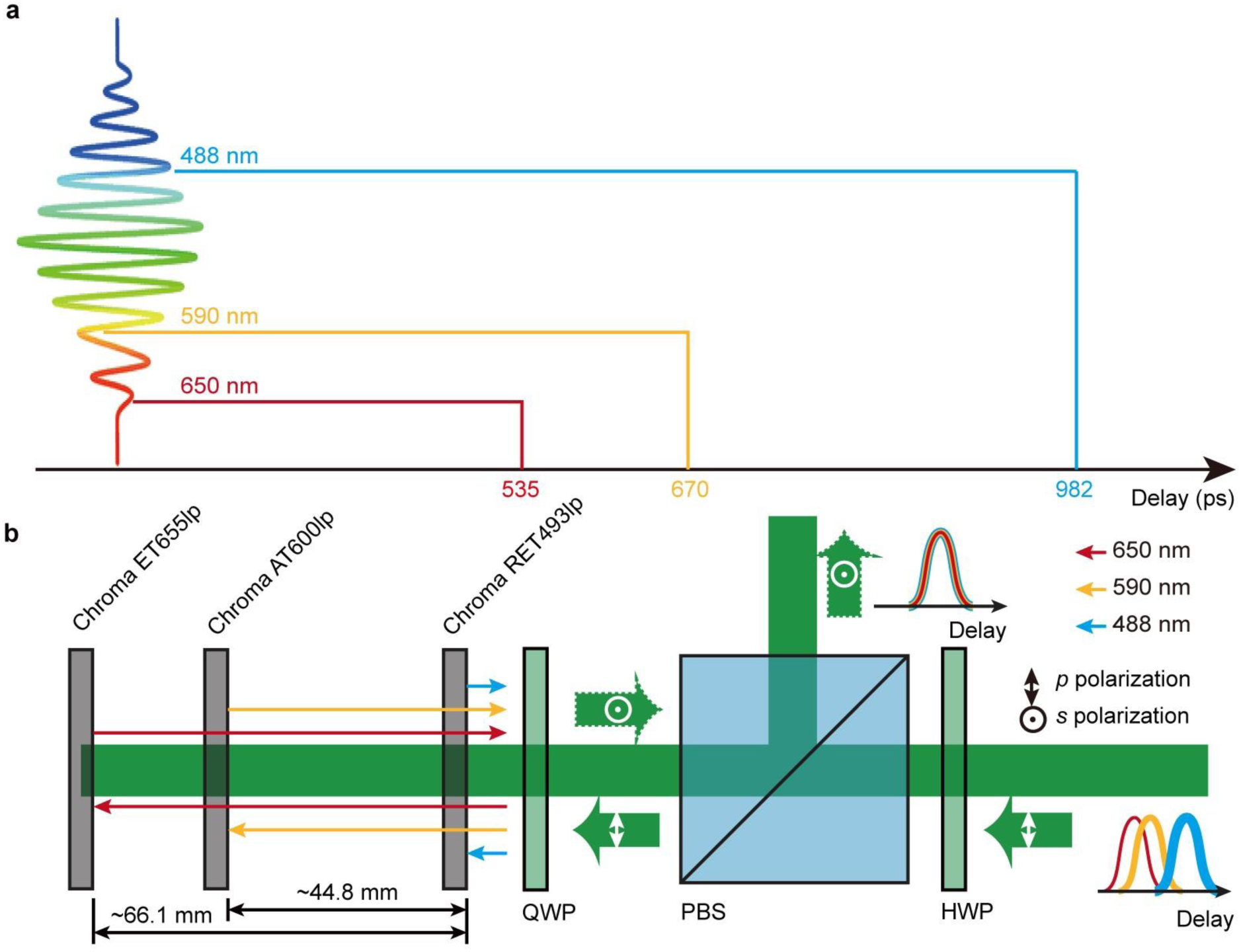
Chromatic dispersion of white-light laser and its compensation strategies. **a**, chromatic dispersion of the white-light laser. The original data was measured by the laser manufacturer (NKT Photonics). **b**, Schematic and principle of our chromatic dispersion compensation module. The incident excitation light with linear polarization passes completely through the polarizing beam splitter (PBS) after a half-wave plate (HWP) adjusts its direction of polarization. The three main components of the beam with wavelengths of 650 nm, 590 nm, and 488 nm, are respectively reflected on three adjacent filters, which produce different relative time delays due to the length of the optical path. The polarization of the reflected beam is rotated 90° as it passes through a quarter-wave plate (QWP) twice and finally exits the module from the reflecting path of the PBS. In our design, all filters are 1 mm in thickness.

To correct for the color-dependent temporal mismatch, we introduced a compensation module (**Fig. 15b**) to the excitation beam path. The linearly polarized excitation beam, which consists of three chromatically dispersed wavelength bands selected by the AOTF, enters this module through a polarizing beam splitter (PBS). Three long-pass filters (**Supplementary Fig. S8** and **Supplementary Table 3**), each reflecting one of the three wavelength bands, are placed sequentially behind the PBS. The spatial distance between adjacent filters introduces a temporal delay to ensure that the light with different wavelengths excites the sample simultaneously. The beam passes twice through a quarter-wave plate oriented at 45°. As a result, the polarization direction of the reflected beam is rotated by 90° so that it is reflected by the PBS on its way out.

#### Resolution measurement

To evaluate the resolution of isoSTED system, sub-resolution sized objects are imaged, and their full widths at half maximum (FWHM) are measured. We recommend using either 100-nm crimson fluorescent microspheres (*e.g.,* Thermo Fisher catalog #8782) or DNA-origami structures (*e.g.,* GATTA–Beads R–brightline, 23-nm DNA origami beads with ATTO 647N fluorophore). After acquiring the bead images, intensity profiles are fitted with a Gaussian or Lorentzian function for confocal imaging or STED imaging, respectively, using MATLAB analysis software (code available at https://zenodo.org/uploads/10574469). The FWHM of the effective PSF, or the resolution, can then be estimated as (**Supplementary Note 1**)

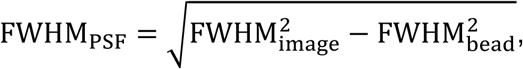

With a spherical model for the beads, 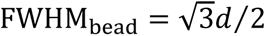, and *d* is the diameter of the beads.

When assessing the resolution of a biological specimen, such as tissue, Fourier Ring Correlation is calculated with two images that are consecutively acquired in an identical FOV. This analysis is performed using open-source ImageJ plugin. We additionally recommend considering the ‘Nested Loop Fitting’ approach^19^ that allows users to assess the resolution from tubular-shaped structures in biological samples (e.g. immunolabeled microtubules).

#### Data format and image visualization

Similar to other confocal and STED microscopes, raw images from our isoSTED nanoscope do not require image post-processing before displaying and quantification. As the sample moves incrementally along the *z*-axis, our nanoscope can acquire snapshot images of each optical section. These optically sectioned images are later compiled and stacked, directly forming a 3D representation of the sample.

To store the raw image data, we utilize the ‘.dbl’ format—a binary file containing data stacks of up to four dimensions. This format includes essential header information about the physical dimensions of the sampled volume. Access to these files is available through our nanoscope control software or via Imspector (Abberior Instruments). Moreover, our custom MATLAB code (available at https://zenodo.org/uploads/10574469) easily converts these files to the more universal ‘TIFF’ format without compromising information integrity.

Following data conversion, numerous software options can facilitate visualization. A good open-source option is, e.g., FluoRender^23^, which was used to visualize the 3D data sets in this protocol.

### 2. Materials

#### 2.1. Reagents

Precision #1.5H glass coverslips, 25-mm diameter (Thorlabs, CG15XH)

39-mm diameter anodized aluminum sample holder with 25.3-mm diameter, 500-µm deep pocket (custom-made; technical drawing see FAB-P0009)

100-nm crimson beads (625/645-nm; Thermo Fisher, custom order)

100-nm red beads (580/605-nm; F8801; Thermo Fisher)

100-nm yellow-green beads (505/515-nm; F8803; Thermo Fisher)

23-nm diameter fluorescent beads, Deoxyribonucleic Acid (DNA) origami conjugated to ATTO-647N (GATTAquant GATTA-Beads R,)

Biotinylated bovine serum albumin solution (1 mg/mL in phosphate-buffered saline; GATTAquant, GATTA-Beads R kit)

Neutravidin solution (1 mg/mL in phosphate-buffered saline; GATTAquant, GATTA-Beads R kit)

Gold Nanoparticles (150-nm diameter, Sigma-Aldrich, 742058-25 mL) ProLong Diamond Antifade Mountant (Invitrogen, P36970)

Picodent TwinSil (two-component silicone adhesive, Picodent, 1300 1000)

Poly-L-Lysine (Sigma-Aldrich, P4707)

### 2.2. Equipment

#### Tools

Mechanical tools: Hex keys set, Metric screw drivers set

Cage alignment plates (*e.g.,* Thorlabs, SCPA1 and CPA1)

Dovetail rails (*e.g.,* Thorlabs, RLA450/M)

Shim spacers (*e.g.*, McMASTER-CARR, stainless steel slotted shim, 9722K812)

Oscilloscope (*e.g.,*Tektronix, MDO3054)

Optical power meter digital console (*e.g.,* Thorlabs, PM100D) and Slim Photodiode Power Sensors (*e.g.,* Thorlabs, S130C, <500 mW) or Thermal Power Sensor Head (*e.g.,*Thorlabs, S425C, 2 mW – 10 W).

Multimeter (*e.g.,* IDEAL 61-360)

Absolute height gauge (*e.g.,* Mitutoyo, Digital ABS Height Gauge, 570-244)

Level (*e.g.,* Thorlabs, 1.25” Circular Bullseye Level, LVL01)

Micrometer (*e.g.,* Mitutoyo, CD-6” CSX)

Laser safety goggles (*e.g.,* Thorlabs, Certified Laser Safety Glasses, LG2)

IR laser card (*e.g.,* Thorlabs, VRC2 - VIS/NIR Detector Card)

Shearing Interferometers (*e.g.,* Thorlabs, SI254)

Alignment camera (*e.g.,* Thorlabs, DCC1545M)

UV lamp (*e.g.,* UVL-18 EL, 365-nm)

UV-curing optical adhesive (*e.g.,* Thorlabs, NOA81)

DC Servo Controller/Driver (Newport, CONEX-CC)

Neutral density filter (*e.g.,* Thorlabs, NE20A)

Laserline cleanup filters: Semrock, LL01-647-25, Semrock FF01-591/6-25 and Semrock LL01-488-25

Linear polarizer (*e.g.,* Thorlabs, GT15-A).

A 1”-diameter, 40-mm focal length optical lens assisting scan mirror alignment (*e.g.,* Thorlabs, LA1422-B)

#### Optomechanical components

The system is composed of numerous commercial and customized optical and optomechanical components. A full part list is available in **Supplementary Table 1** and a SolidWorks model of the system assembly is available at https://zenodo.org/uploads/10574469.

#### Nanoscope control computer

Operating system (OS): Windows 10 Pro, 64-bit

Central processing unit (CPU): 12-core Intel(R) Core (TM) i7-7800X CPU at 3.5 GHz

Random access memory (RAM): DDR4, 32 GB

Graphic processing unit (GPU): NVIDIA GeForce GTX 1060 6G

Hard drive: Samsung SSD 970 EVO 1 TB and WDC WDS200T2B0A-00SM50 2 TB

Motherboard: ASUSTeK WS X299 SAGE

#### Nanoscope control software

National Instruments, LabVIEW 2018 32-bit, device drivers, VISA, Measurement & Automation Explorer (MAX), Vision Development Module, and FPGA Development Module

Boston Micromachines, LinkUiMulti, DM Shapes

MPB Communications, Pulsed Fiber Laser Control

NKT Photonics, CONTROL

Physik Instrumente (PI), PIMikroMove

Thorlabs, Kinesis and ThorCam

Applied Scientific Instrumentation (ASI), ASI Console

IDS Imaging Development Systems, Camera Manager and uEye Cockpit

Quanta-Tech (AA Opto-Electronic), MPDSnC

Opsero, LUFA VirtualSerial.inf

Hamamatsu Photonics, LCOS-SLM Control

#### Data visualization software

MathWorks, MATLAB 2018a 64-bit or later

University of Utah, FluoRender 2.22.3 or later

Abberior Instruments, Imspector 0.13.11588 or later

National Institute of Health, ImageJ 2.3.0/1.54b or later

### 2.3. Reagent setup

#### Fluorescent bead samples

Fluorescent bead samples are required during instrument alignment and resolution measurement steps. DNA origami beads labeled with ATTO 647N with a diameter of 23 nm are particularly recommended for the resolution test. In our experimental setup, fluorescent and gold nanoparticles were deposited onto a 25-mm diameter round coverslip, which was subsequently covered with another 25-mm diameter round coverslip using the ProLong Diamond antifade mountant. This sandwiched configuration ensures optimal imaging conditions. The assembled coverslips were securely positioned within a custom aluminum sample holder and firmly secured using two-component silicone adhesive.

#### Biological test samples

ATTO 647N-labeled microtubule samples are recommended for testing the imaging performance of the system. The immunolabeling process for isoSTED nanoscopy closely mirrors that for other STED nanoscopy applications and has been previously described, e.g. in subsection ‘Microtubules’ in Ref. [12]. For isoSTED imaging, we recommend additionally depositing both 150-nm diameter gold nanoparticles and 100-nm diameter fluorescent beads on the opposing coverslip to evaluate and modify system alignment before imaging the biological sample. Start preparing the coverslip with bead samples by mixing gold nanoparticles and fluorescent beads at a 1:10,000 (vol/vol) dilution. Add 25 µL of poly-L-Lysine to the coverslip, allow to sit for 15 minutes, and then aspirate poly-L-Lysine. Add 25 µL of the bead mixture, allow to sit for 15 minutes, and then aspirate the bead mixture. Place the coverslip with beads in the sample holder and add a drop of curing mountant. Place the coverslip with the cells upside down onto the coverslip in the sample holder. Gently blot the excess liquid from the coverslip sandwich using a lint-free cleaning tissue. Seal the coverslips and fixate them to the sample holder using two-component silicone adhesive.

### 2.4. Equipment setup

#### SLM and DM calibration

For aberration correction (**Fig. 16a–c**), we utilize two DMs (**Fig. 16d**) in the 4Pi interference cavity and an SLM (**Fig. 16e**) in the depletion beam path to achieve a sensorless AO architecture. To maximize the performance of AO, both SLM and DMs require calibration.

**Fig. 16.**
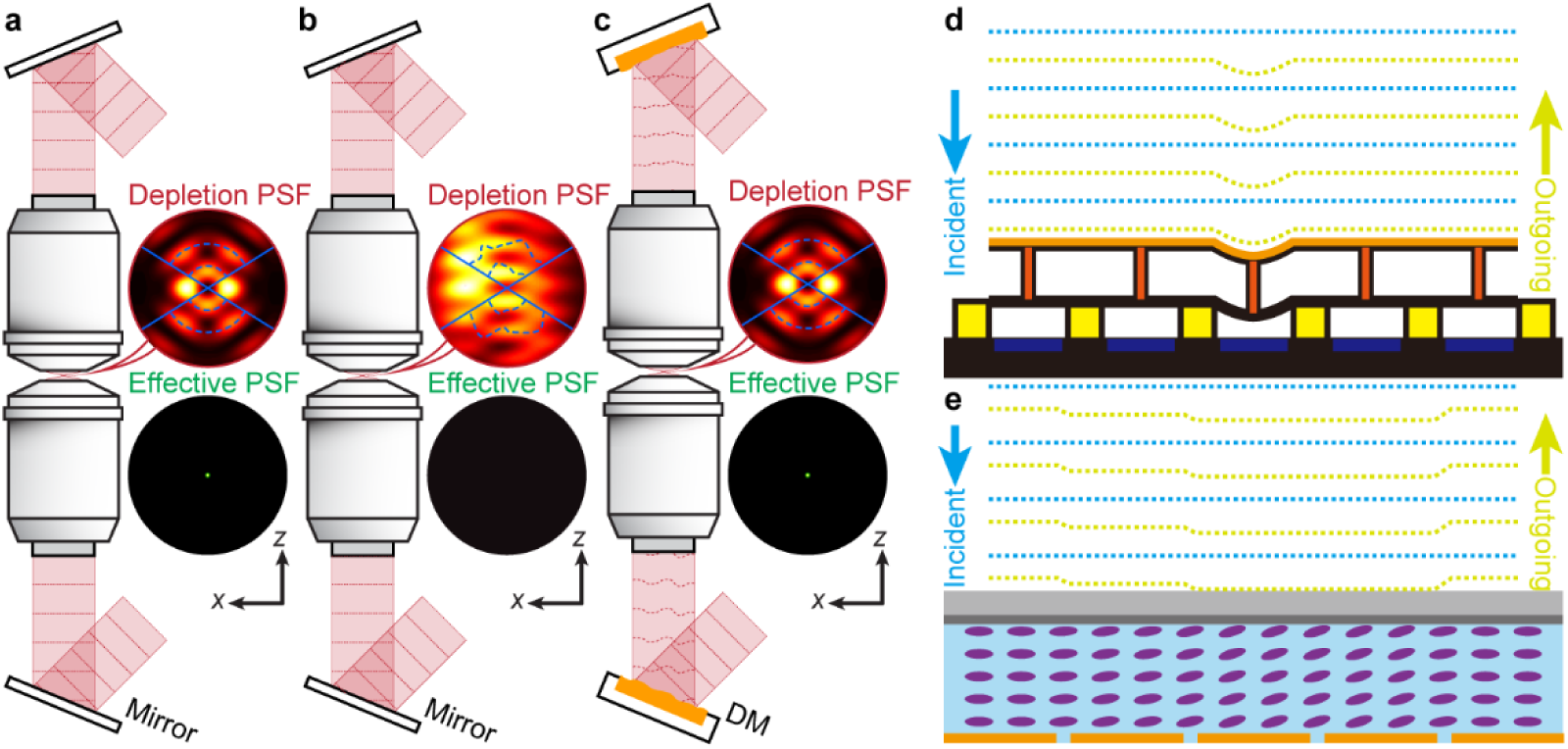
AO in isoSTED nanoscopy. **a**-**c**, Principle of aberration correction in isoSTED nanoscope. Depletion and effective PSFs with and without sample-induced aberrations are shown as **a** and **b**, respectively. **c**, The DMs in the interference cavity introduce phase patterns that counteract aberrations, restoring nearly perfect optical performance. **d**, **e**, Schematic of AO devices. Specifically, in a DM (**d**), the shape of the front reflection membrane (mirror layer) is controlled by discrete actuators. When a voltage is put on an electrode, the actuator layer above it is attracted, and it pulls the mirror layer down. Changing the shape of the mirror layer enables us to control the wavefront of the reflected beam (highlighted in yellow). In an SLM (**e**), as liquid crystals are birefringent, applying a voltage to the cell changes the effective refractive index seen by the incident wave, and thus the phase retardation of the reflected wave (highlighted in yellow). (**a**-**c** reprinted from Ref. [12]).

The DM features a light reflective membrane that can be deformed dynamically by an array of software-controllable actuators. DM calibration is vitally important to characterize how the reflective surface of the mirror reacts to the pokes of the actuators. Each DM is calibrated offline using an external Twyman-Green interferometer in which the DM is systematically controlled and analyzed by custom code programmed in Python, as described in detail in Ref. [24]. In brief, tilting the flat mirror in the interferometer’s reference arm allows for the acquisition of the phase induced by the DM. This phase is derived through Fourier fringe analysis and phase unwrapping. The pivotal task of DM calibration involves computing the control matrix, denoted as *H*. This matrix acts as the guiding map, connecting a vector **u**—comprising the voltages assigned to each actuator of the DM—to its corresponding vector **z**, containing the coefficients derived from the Zernike analysis of the phase. The computation of matrix *H* involves gathering a series of paired input and output vectors—**u** and **z**, respectively—followed by solving a least-squares problem. This method allows for the derivation of *H*, unraveling the intricate relationship between actuator voltages and the resulting Zernike phase coefficients.

Similarly, the SLM calibration is critical to determine the phase response, corresponding to the gray level of each pixel of the SLM, and construct a comprehensive look-up table (LUT). Amidst various calibration approaches, we have chosen diffractogram analysis^25^ for its distinct advantages. While offline calibration is also possible, the implementation of diffractogram analysis facilitates convenient *in-situ* calibration of the SLM by placing an additional camera immediately after the SLM. During diffractogram analysis, a binary hologram is displayed on the SLM under the illumination of a linearly polarized beam. The camera captures the diffracted beam pattern produced by the hologram on the SLM. By examining the intensity variations between the diffraction peaks and valleys, alongside any fringe shifts within the pattern, we can ascertain the phase discrepancies across distinct binary regions of the SLM, as elaborated in Ref. [25]. The principle of diffraction establishes a link between phase disparities and the corresponding gray levels of the pixels, enhancing our understanding of pixel-specific responses.

Notably, the control matrix of the DM and the LUT of the SLM may be available from the manufacturers, but we highly advocate for calibrating these devices just before integrating them into the nanoscope. The nuances of the local electronic environment can potentially impact the response of the pixels/actuators.

#### Optomechanical assembly

In general, the instrument setup follows the CAD model. Assemble all parts including custom-manufactured mechanical components, stages, optical components, commercial optomechanical components, light sources, dowel pins, SLM, and DMs following the CAD model.

**! Note** We highly recommend manually checking the dimensions of every custom-machined mechanical component before integrating it into the system (ideally already at the time of delivery). In our experience, manufacturing errors are unavoidable and can cause serious optical alignment issues if not detected and corrected early in the process.

**! Note** Pay close attention to details, especially regarding optical components such as lens mounting depth in holders, as well as the orientation of lenses, filters, and retarders.

**! Note** The *xy* piezo stage (PI, P-612.2SL) holding OBJ2 may have significant vibration issues in closed loop mode when a heavy objective is installed. Adjusting the gain settings in the controller can solve this problem. We suggest consulting the manual of the piezo stage or inquiring with the manufacturer about the specific tuning procedure to accommodate the objective and objective mount’s mass on the stage.

#### PBS cube installation

The PBS cube is glued to a custom part (FAB-IS0002, custom-designed) by using an ultraviolet (UV)-curing optical adhesive (*e.g*., Thorlabs, NOA81) with UV lamp illumination. Since the PBS is a central component of the nanoscope, it is essential to position the PBS accurately and securely glue it in place.

#### Translation stage tower installation

The tower assembly is detailed in the CAD model. A pair of ASI stages (LS-50) is mounted on the two pillar legs. They hold the assembly of the sample stages and the objectives, facilitating axial translations of this assembly as a whole. This allows operators to change the optical path difference between upper and lower arms of the 4Pi interference cavity over a long range without changing the alignment between the objectives and the sample. Precise alignment is enabled by the ASI stages’ two operational modes: synchronous movement and independent stage adjustment.

For initial assembly, we advise securing one pillar leg to the customized base plate while slightly loosening the other. Next, translate the ASI stage that can be moved independently until it reaches the same height as the other stage, verified using a height gauge. Once the heights are aligned, switch to synchronous movement of the stages.

Next, move the ASI stages up and down to the highest and lowest positions the stages can reach without colliding with other components. Adjust the position of the loose supporting leg until the stages move smoothly for the complete travel range and fix the loose supporting leg in place by tightening the screws connecting it to the optical table.

Finally, move the ASI stages vertically to the position indicated in the CAD model. Note that the ASI stages are rarely moved post-installation, underscoring the importance of proper alignment during the setup phase.

#### Sample stage leveling

The sample is mounted on a leveled stage, where the *xy* and *z* sample stage stack are held by three linear actuators (PI, M-227.10) positioned beneath. Coarse translation along the *z*-axis is achieved by simultaneously adjusting all three actuators, while individual manipulation of the actuators enables tip and tilt motions of the stage stack. A bubble level gauge, such as the LVL01 from Thorlabs, can be used to guide the leveling process. Each actuator is individually adjusted until the sample stage stack is oriented horizontally. Finally, the three linear actuators are simultaneously translated along the *z*-axis to position the sample stage to be approximately aligned with the position in the CAD model.

#### QWP setup in 4Pi interference cavity

In the 4Pi interference cavity setup, described in detail below, precise tuning of the QWPs near the objectives is paramount. The QWPs play a crucial role converting linearly polarized depletion beams into circularly polarized ones to produce beam profiles essential for generating ring-shaped depletion foci. Additionally, circular polarization of the excitation light ensures uniform excitation of fluorophores with different dipole orientations. The orientation of the QWPs dictates whether the depletion beams’ polarization becomes left-handed or right-handed circular polarization. The choice between these orientations is critical for isoSTED nanoscopy, where the circular polarization direction must align with the rotational direction of the vortex phase pattern imprinted on the depletion beam. This alignment is necessary to prevent high laser intensity residue at the center of the ring-shaped beam, which reduces fluorescence signals and negatively impacts the resolution^26^. Right-handed circular polarization corresponds to an anti-clockwise vortex phase pattern, while left-handed circular polarization serves the same purpose but matches the opposite direction for the vortex phase pattern.

#### Device drivers and software installation

Prior to initiating the alignment protocol, connect all computer-controlled devices to the computer and install the manufacturer-supplied drivers. Confirm that control software and devices are functional. The necessary software packages for installation include:

Boston Micromachines, LinkUiMulti and DM Shapes for DM control

NKT Photonics, CONTROL for white-light laser control

MPB, Pulsed Fiber Laser Control for depletion laser control

PI, PIMikroMove for piezo, sample, objective Z and DC-Mike control

Thorlabs, Kinesis for BK7 wedge motor control; ThorCam for alignment camera control

ASI, ASI Console for LS-50 linear translation stage control

IDS Imaging, uEye camera drivers and camera manager for alignment camera control

Opsero, LUFA VirtualSerial.inf file for pulse gate/delay module AA Opto-Electronic, MPDSnC for AOTF control

Hamamatsu Photonics, LCOS-SLM Controller

To operate the nanoscope control software (**Fig. 17**), ensure that various software packages from National Instruments (NI) are installed on the computer. These are:

**Fig. 17.**
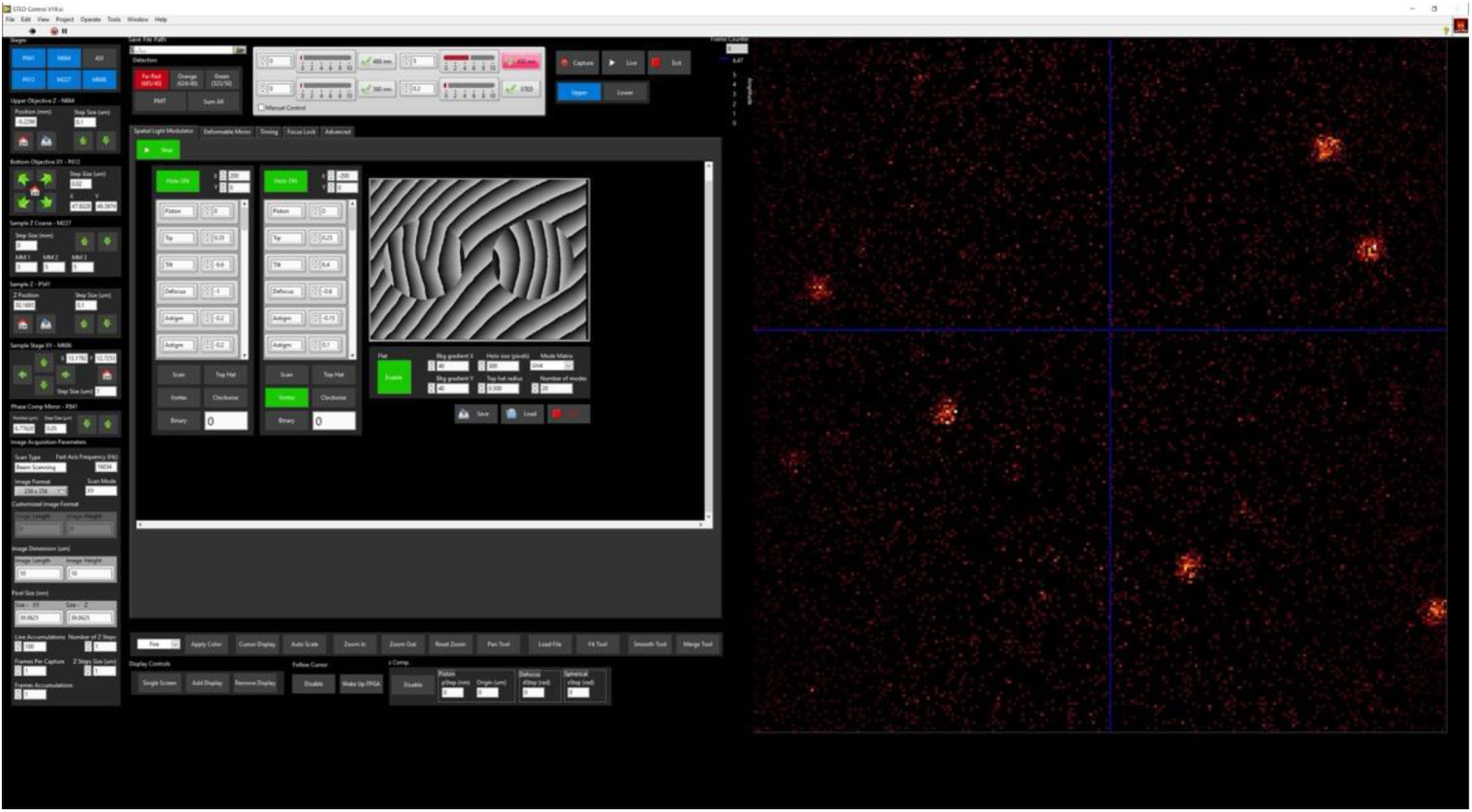
User interface of the LabVIEW-based nanoscope control software.

LabVIEW 2018 32-bit development environment

All-in-one package of NI device drivers (version January 2019) including NI-DAQ™mx, NI MAX, NI-VISA, NI Vision, NI-RIO, and NI-Serial

NI Vision Development Module

NI FPGA Development Module

To visualize the data, we further need:

MathWorks, MATLAB 2018a 64-bit or later

University of Utah, FluoRender 2.22.3 or later

Abberior Instruments, Imspector 0.13.11588 or later

National Institute of Health, ImageJ 2.3.0/1.54b or later

## 3. Procedure

### 3.1. Software installation and setup ⬤ Timing: 1 day

1. Install LabVIEW development environment and all necessary add-ons as detailed in ‘Nanoscope control software’.

2. Connect all computer-controlled devices such as stages, lasers, and FPGA boards to the computer. Install the necessary manufacturer-supplied drivers and control software. Confirm the proper functionality of all devices to ensure they are working correctly.

3. Install custom nanoscope control software.

4. Press the ‘Run’ button of the FPGA VI to initiate the compilation process. Follow the provided instructions to execute the Xilinx compiler, generating the bitfiles for the FPGA boards.

### 3.2. Optomechanical assembly ⬤ Timing: 1 week

4. To ensure accurate installation of the mechanical components and proper fixation of screws, refer to the SolidWorks model provided at the following link: https://zenodo.org/uploads/10574469. During installation, pay attention to mounts equipped with pins designed to be installed on the vertical breadboard. These pins aid in securing the components in their designed positions and ensure optimal functionality of the setup. Once the mechanical components are installed, and screws are properly fixed, verify alignment and stability to ensure the successful assembly of the apparatus.

5. Ensure that all optical components, including lenses, mirrors, dichroic mirrors, waveplates, etc., are securely fixed in their respective mounts either mechanically or using suitable adhesive. However, avoid tightening screws that directly exert force on optical elements, especially mirrors, too much since that will deform these elements and can cause severe optical aberrations. Pay close attention to the orientation of lenses and filters, as they are often not reversible. Properly fix these components in their mounts to maintain the desired functionality. Install all the assemblies in accordance with the instructions provided in the ‘Equipment setup’ guidelines.

### 3.3. Electronics assembly ⬤ Timing: 1 week

7. Connect cables between the FPGA connectors and their corresponding breakout boxes.

8. Connect all cables and wires from the electronic controlled devices, including stages, APDs, PMT, resonant mirror, galvanometer mirrors, AOM, and AOTF, to their respective terminals in the breakout boxes. Refer to **Supplementary Table 2** for the FPGA wire configurations.

### 3.4. AOM setting and alignment ⬤ Timing: 2 hours

9. Read Section ‘Procedure overview’, especially the ‘AOM and AOTF alignment’ subsection.

10. Install LH1 and measure its power using a laser power meter to confirm that the specified output laser power is available. (>2 W of average power from the depletion laser source is required.) Confirm that the beam exits LH1 at the specified beam height of the SLM screen center.

11. To adjust M1, temporarily install an optical rail and, using M1, align the laser beam to run parallel to the rail, following the general beam alignment procedure described in ‘General beam alignment’.

12. Mount the AOM on an angular alignment stage (Newport, 9071-M) and add posts and spacers as needed, so that the laser beam passes through the centers of the entrance and exit windows of the AOM. Turn on the AOM and set the frequency and amplitude using the manufacturer-provided software (MPDSnC) until a diffracted beam appears at the +1 order.

13. Adjust the angle of the AOM using the angular alignment stage to optimize the power of the +1 order diffracted beam.

14. Use the manufacturer-provided software (MPDSnC) to tune the drive frequency of the AOM to maximize the output power in the +1 order diffracted beam.

15. Repeat the optimization of Steps 13–14 to maximize the power of the +1 order diffracted spot, which should surpass 90% of the input power.

### 3.5. Polarization separation module alignment ⬤ Timing: 4 hours

16. Use M2 to reflect the +1 order diffracted light beam emerging from the AOM and align it according to the layout depicted in **Fig. 2** and **Fig. 4**. For this purpose, temporarily install a rail and align the beam according to ‘**General beam alignment**’.

17. Install the assembled polarization separation module (**Fig. 6**). Use a cage alignment plate (*e.g*., Thorlabs, CPA1) to align the polarization separation module along the beam. The input beam and the transmitted beam through PBS1 should hit the center of the cage alignment plate when placed in front of PBS1 and M3.

18. Align the reflection arm of the polarization separation module (used for the STED*_xy_* beam) by rotating PBS1 so that the reflected beam hits the center of a cage alignment plate placed behind QWP2.

19. Place the cage alignment plate behind HWP2. Block the transmitted light behind QWP1 and adjust M4 so that the reflected beam hits the center of the cage alignment plate.

20. Rotate QWP2 to maximize the output power at the exit of the polarization separation module.

21. To align the transmission arm (used for the STED*_z_* beam), block the beam behind QWP2 and adjust M3 to reflect the beam to hit the center of the cage alignment plate. Rotate QWP1 to maximize the output power at the exit of the polarization separation module.

22. Rotate the angle of HWP1 to allocate the power for the STED*_z_* to STED*_xy_*. beams to a ratio of 1:5.5. In general, for simplicity and to avoid confusion, keep using a single depletion beam (e.g., STED*_xy_* beam) and block the other beam at the respective arm of the polarization separation module for the following system alignment. Switch beams or apply both only when necessary.

23. Rotate the angle of HWP2 to set the polarization of the STED*_xy_* beam to horizontal orientation. The STED*_z_* beam will automatically be polarized perpendicularly to the STED*_xy_* beam. The polarization orientation can be checked by measuring the laser power behind a polarizer and will be validated by the modulation of the SLM once aligned.

### 3.6. SLM installation and alignment ⬤ Timing: 1 day

24. Adjust the mirrors M5 and M6 to align the beam along the center of R1 at the height of the SLM hologram center. Follow the method in **Box 3** to align the L1-L2 4f telecentric system for beam expansion.

25. Temporarily install a rail at the designated position of the D-shape mirror (DS) before installing the DS. Mount an alignment camera on the temporary rail at the designated DS position to align the beam following **Box 1**. Adjust the tip and tilt angles of M7 and translate it along R1 until the beam is aligned to the center of the rail.

26. Remove the temporary rail and install DS in the designated position. Translate M7 along R1 to have the beam centered on DS if its edges cut off the beam.

27. Temporarily install a rail at the designated position of the SLM along the designated beam direction reflected by DS. Mount the alignment camera on the rail. Translate DS using N1 and adjust its tip and tilt angles until the beam reflected by DS propagates parallel to the rail. The tip and tilt angles of DS are adjusted in this step while its translation will be adjusted later. ▲Critical There are two dowel pins located at the bottom of the DS mount (FAB-IS0050, custom-designed). These pins serve to restrict the movement of DS to the propagation direction of the incident beam. Ensure these pins are in place before translating DS.

28. Remove the temporary rail and mount the SLM on the base plate. Use a ruler to roughly place the SLM screen center at the focus of L3. Apply a 0/π top-hat hologram pattern (300 pixels in diameter) along with the blazed gratings (set ‘Tip’ = 7 on the ‘SLM’ panel in the LabVIEW-based control software) on the SLM. Ensure the hologram is positioned at the center of the SLM (‘Holo 1’ coordinates: *x* = 0, *y* = 0 on the ‘SLM’ panel, as shown in **Fig. 18a**). Slightly loosen the screws that secure the DS mount. Translate the DS until the hologram is uniformly illuminated, checked visually by a paper screen held into the beam after it has been reflected at the SLM. Re-tighten the screws. ▲Critical To minimize the stray light outside the objective apertures of the nanoscope (represented here by the size of the hologram pattern), apply another blazed grating pattern to the SLM area outside the hologram(s) (*e.g.,* ‘Bkg gradient X’ = 40, ‘Bkg gradient Y’ = 40 on the ‘SLM’ panel; see **Fig. 18**).

**Fig. 18.**
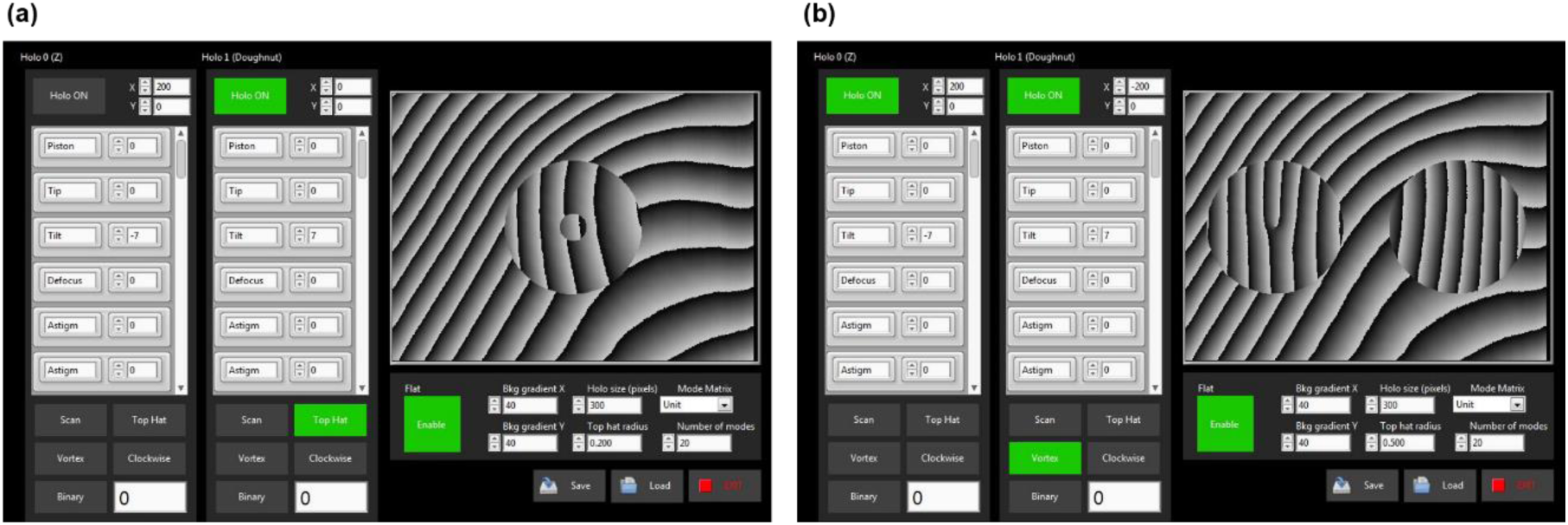
User interface of ‘SLM’ control panel. **(a)** shows the single-hologram mode at the center of the SLM used for alignment purposes only; **(b)** shows the double-hologram mode used to modulate the STED_xy_ and STED_z_ beams.

29. ▴Critical Temporarily replace R2 with a long rail (*e.g.*, Thorlabs, RLA300/M). Mount an alignment camera on this rail to check the beam. Turn on the ‘Top Hat’ hologram with ‘Top hat radius’ set to ∼0.2 to highlight a clear hologram center imprinted on the beam. Adjust the tip and tilt angles of the SLM by rotating it on its mount and utilizing spacer shims to fine-tune the direction of the reflected beam until it runs parallel to the rail. Alternate the illumination beam between STED*_xy_* and STED*_z_* beams by blocking either the reflected or transmitted arm of the polarization separation module and adjust HWP2 until only the STED*_xy_* but not the STED*_z_* beam is modulated by the applied hologram (**Fig. 18a**).

30. Replace the temporary rail with R2 and install QWP3 and the L3-mount without the lens. Add an iris to the L3 mount and place an alignment camera after QWP3 for beam monitoring. Adjust the linear stage underneath the SLM in the direction perpendicular to the incident beam to center the top-hat pattern through the iris. Translate DS for uniform illumination of the hologram on the SLM (**Fig. 18a**). Mark the current beam center on the camera in the ThorCam software. Install the lens L3 in the mount and tune its position to maintain the beam position on the camera.

31. Mount M8 on R2 at the focal plane of L3. Adjust the angle of M8 to reflect the beam back along the incident beam path through the same iris on L3. **! Note:** To determine the focal position of L3, speckle analysis^27^ may be applied.

32. Activate both holograms in the LabVIEW program and arrange them side by side as displayed in **Fig. 18b**, Set the coordinates of the two hologram centers to (-200, 0) and (200, 0), respectively. Apply blazed gratings with identical periods but in opposite directions to the holograms (*i.e.,* for the left hologram: ‘Tip’ = +7, and for the right hologram: ‘Tip’ = -7, on the ‘SLM’ panel).

33. Translate DS to uniformly illuminate the left hologram on the SLM. Ideally, with the well aligned 4f-system on R2 (consisting of passing twice through L3), the reflected beam from M8, after passing through L3, should illuminate the right hologram with both holograms now being co-centered, ready for further tuning later.

34. Mount an alignment camera on R3 to monitor the beams. Illuminate only with the STED*_xy_* beam (modulated by the left hologram). Rotate QWP3 while sequentially adding top-hat patterns to either of the two holograms until the beam modulation by the right hologram is minimized, aiming for as little modulation as possible. Then, illuminate with both STED*_xy_* and STED*_z_* beams. Ideally, the two hologram patterns are co-centered; if not, adjust the angles of M8 to correct it.

35. Move the alignment camera along R3 to verify that the beam is parallel to R3 following instructions of **Box 1**. If a beam direction mismatch is detected, tune the angles of DS to align the beam direction with R3 and adjust M7 to ensure a uniform illumination of the holograms. Install the L4 lens mount with an iris, without the lens, in front of the camera to verify that the hologram center passes through the iris center. Translate the SLM in the direction perpendicular to its incident beam to center the right hologram with the iris. Translate the SLM in the direction parallel to its incident beam to have the left and right holograms co-centered as much as possible.

36. Mount L4 and L5 on R3. Use a ruler to set L4 at a distance equal to its focal length from the right hologram on the SLM. Align this 4f telecentric system following **Box 3**General beam alignment. At the polarization separation module, block the beam modulated by the left hologram (STED*_xy_* beam) and illuminate only with the beam modulated by the right hologram (STED*_z_* beam). Install an alignment camera after L5 (approximately at the location of M9) and securely position the camera on R3 where a clear image of the right hologram pattern is visible (See **Supplementary Video S1** for an example).

37. Illuminate only with the STED*_z_* beam modulated by the right hologram. Check the hologram image on the camera whether it is in focus. Adjust N3 to move R2 along the rail direction to ensure a clear image on the camera. ▲Critical Do not change the angle of R2 while tuning its position. This can be achieved by pushing R2 against the corresponding edge of the Assembly Plate (FAB-IS0042).

38. Keep the camera at its current position and verify the following factors: (i) Ensure a clear image of both holograms is observed by switching between the two depletion beams (optimized by N3); (ii) Check for co-centering of the two holograms (optimized by M8); (iii) Verify the illumination uniformity across the holograms (optimized by M7). Optimize these factors if necessary. Once all these conditions are met, secure all components and avoid any further changes.

39. Remove the alignment camera and install M9 in its place. Adjust the angles of M9 to reflect the beam along R4. Move M9 along R3 to align the reflected beam with R4.

40. Align the L6-L7 4f telecentric system following **Box 3**.

41. Install an iris in the intermediate focal plane between L6 and L7 as a special filter to block stray light.

### 3.7. Excitation path alignment ⬤ Timing: 3 days

42. Read ‘Procedure overview’, especially the ‘Excitation laser path alignment’ subsection.

43. Install the laser head for excitation (LH2) and set the beam height the same as the depletion laser.

44. Mount the AOTF on the angular alignment stage (Newport, 9071-M) and add posts and spacers to match with the laser beam height.

45. Reflect the laser beam with M15 to pass through the centers of the entrance and exit windows of the AOTF.

46. Temporarily insert a filter stack between LH2 and AOTF, consisting of a neutral density filter (*e.g.*, Thorlabs, NE20A), a laserline cleanup filter (*e.g*., Semrock, LL01-647-25), and a linear polarizer (*e.g.,* Thorlabs, GT15-A). Ensure that this filter stack does not alter the beam direction and the orientation of the polarizer matches with the polarization modulated by AOTF.

47. In the MPDSnC software, enable Channel #1 and designate the operational wavelength of the AOTF as 650 nm. ! Note: In MPDSnC, only the driving frequency, not the wavelength, is directly adjustable. Refer to the manufacturer-provided calibration sheet to find the driving frequency matching the wavelength of interest.

48. Adjust the angle of the AOTF using the angular alignment stage it is mounted on to maximize the power of the +1 order diffracted beam with the 650-nm laser.

49. In MPDSnC, optimize the driving amplitude of the AOTF, aiming to maximize the output power of the 650-nm laser component. ! Note: The largest driving amplitude does not necessarily result in the highest power output.

50. Repeat Steps 48–49 to maximize the power of the +1 order diffracted spot.

51. Replace the laserline cleanup filter with appropriate filters for the other two desired excitation wavelengths, such as Semrock FF01-591/6-25 for 590 nm and Semrock LL01-488-25 for 488 nm, respectively. Configure the corresponding driving frequencies in Channels #2 and #3 in MPDSnC. Activate each channel sequentially while deactivating the others. Adjust the driving frequency and amplitude for each channel to maximize the power of the +1 order diffracted beam for each wavelength.

52. Remove the cleanup filters for subsequent alignment and imaging applications. Use the 650-nm laser as the default for the remaining alignment procedures.

53. Use M16 to direct the beam path towards R7.

54. Install the assembled cage system of the chromatic dispersion compensation module (**Fig. 15**). Align the cage system along the beam using the cage alignment plate (*e.g.,* Thorlabs, SCPA1). The laser beam should hit the center of the cage alignment plate in front of PBS3 and behind QWP6.

55. Adjust the angles of PBS3 to direct the reflected beam going through the hole in the center of the alignment plate on the cage system. Rotate HWP5 to maximize the power of the beam transmitted through PBS3.

56. Install F1, F2, and F3 according to the CAD model. ! Note: To minimize chromatic dispersion effects, position filter F2 and F3 44.8 mm and 66.1 mm from F1, respectively. The optimal positions may vary depending on the laser dispersion. For further details, refer to Section ‘Chromatic dispersion compensation module’.

57. Adjust the angles of F1, F2, and F3 so that their corresponding reflected beams are re-combined to the same path through the center of the alignment plate on the cage system. Note that the adjustment of F1, F2 and F3 should be using 488-nm, 590-nm, and 650-nm laser from the AOTF channels, respectively.

58. Rotate QWP6 to maximize the power of the beam reflected by PBS3.

59. Install M17, L15, L16, QWP7, and M18 according to the layout in **Fig. 4**. Align the L15-L16 4f telecentric system and ensure the beam travels along the direction of the rail.

60. Adjust M18 and DI1 to couple the excitation beam into the depletion beam path. Switch between the three different excitation beams and ensure that all three beams travel the same path as the two depletion beams along R4 after DI1. If not, adjust F2 and F3. ! Note: Tuning the angles of DI1 affects not only the reflected excitation beam path, but also slightly affects the transmitted depletion beam path.

61. HWP6 and QWP7 remain to be tuned after aligning the 4Pi interference cavity (refer to Section ‘4Pi interference cavity alignment ⬤ Timing: 1 week’) in order to maximize the excitation power in the upper arm of the cavity.

62. Keep using the depletion beam for the following alignment procedure unless specified otherwise.

### 3.8. Scanner installation and alignment ⬤ Timing: 1 day

63. Ensure all the three excitation beams and both depletion beams travel the same path along R4 after DI1 using the checking method in Step 60.

64. Temporarily mount a dovetail rail in the propagation direction of the beam emerging from DI2 at the designated position of the scan mirrors. Adjust the angles of DI2 to align the beam direction along the rail. ▲Critical: Ensure the beam reflected by DI2 is horizontal to the baseboard; otherwise, the fluorescence detection light path, which shares the same beam path but in the reversed direction, will experience large beam height changes over its long beam path.

65. Remove the temporary rail from Step 64. Mount the resonant mirror (RM) and two galvanometer mirrors (GM1 and GM2) in the custom-built mount on translational stage LS2. **! Note** Before installing RM, remove its protective case to prevent obstruction of the light beam. Position the stage at its designated location on the base plate. Translate LS2 so that the beam hits the center of RM.

66. Temporarily adjust the positions of GM1 and GM2 to ensure the beam reflected by RM is not obstructed by either of the galvanometer mirrors.

67. Temporarily mount a dovetail rail onto the base plate in the propagation direction of the beam emerging from GM2 at the designated position of L8 and M10. Loosen the screw that secures RM and adjust RM until the reflected beam travels horizontally along the rail. Once aligned, tighten the screw to secure RM in its holder. Translate the holder of RM in the direction perpendicular to the vertical breadboard within the scanner mount until the beam is about 80 mm to the vertical breadboard.

68. Temporarily install a lens with a 40-mm focal length (*e.g.,* Thorlabs, LA1422-B) in the L8 lens mount. Increase the RM scanning amplitude to approximately 20 μm in the LabVIEW control program. Translate the alignment camera along the rail to find the position where the beam image shows minimal scanning motion (i.e. no beam image elongation induced by scanning). Activate the top-hat pattern on the SLM in the LabVIEW control program. Adjust the height of the RM mount using shims until the beam center shows minimal blur during scanning, indicating that the RM scanning axis is aligned with the center of the beam hologram applied by the SLM.

69. Set the amplitude of the RM control voltage to 0. Translate the alignment camera on the rail toward the far end to monitor the position of the beam reflected by GM2. ▲Critical Keep the RM and the GMs turned on and maintain the control voltage setting for all subsequent alignment procedures.

70. Adjust the angles of GM1 and GM2 according to the instructions in **Box 2** of the ‘**General beam alignment**’ section until the beam travels along the rail after passing both galvanometer mirrors.

71. Loosen the screws (**Supplementary Fig. S9)** securing the RM mount. Translate the RM mount in the direction of the incident beam until the beam hits the center of GM2. Once aligned, tighten the screws to secure the RM mount in its new position. Repeat Step 70 as needed.

72. Mount M10 at the designated position without mounting L8. Install R5 onto the vertical breadboard and mount an alignment camera assembly (**Supplementary Fig. S10**) on R5. Adjust the angles of M10 until the beam propagates along R5. Secure the alignment camera at the designated position of the DM surface (**Fig. 9**) and mark the beam center on the camera image.

73. Mount L9. This might cause a shift of the beam position on the camera image. Mark the new beam position as the target and compare the beam positions before and after L9 installation. If there is a shift perpendicular to the vertical breadboard, loosen the screws securing the pillar legs and translate the breadboard in the same direction for compensation. If there is a vertical shift, translate M10 along the propagation direction of the beam emerging from L8. Repeat Steps 70**–**72 as needed until inserting L9 does not alter the beam center position on the camera anymore.

74. Keep the alignment camera at the designated DM surface position for subsequent alignment until otherwise stated. Mount L8 with two pins. Translate its position in the direction perpendicular to the incident beam from the scanner and use shims for beam height compensation as needed until the insertion of L8 does not change the beam center position marked in Step 73 on the camera.

### 3.9. PBS installation and angle alignment ⬤ Timing: 3 hours

75. Install the rotation stage (Newport, M-RS65) on the vertical optical breadboard.

76. Mount the goniometer spacer (FAB-IS0001, custom-designed) with the rotation stage M-RS65.

77. Mount the goniometer (Newport, M-GON40-U) on FAB-IS0001 and ensure that the orientation of the goniometer matches the CAD model.

78. Mount motorized linear actuators (Newport, CONEX-TRA12CC) as the actuators of the rotation stage M-RS65 and goniometer M-GON40-U.

79. Place the goniometer cube holder (FAB-IS0002, custom-designed) on a flat work surface. **! Note** The bottom of the cube holder is not flat. To level it, insert a 2 mm-thick spacer underneath it.

80. Apply one drop of UV curing adhesive to the surface of the cube holder and place PBS2 (CVI laser optics, PBSH-450-1300-100) on top, ensuring it is positioned that the side surfaces of PBS2 attach to the datum plane on the cube holder (**Supplementary Fig. S1**).

81. Place the cube holder about 30 cm away from the UV lamp. Turn on the UV lamp and irradiate the optical adhesive for 15 minutes.

82. Mount the cube holder on the goniometer M-GON40-U (Newport). **! Note** Confirm that the UV adhesive is fully cured to avoid PBS2 from detaching when mounted on the vertical breadboard.

83. Glue the BK-7 glass block onto its holder (FAB-IS0010, custom-designed) using optical adhesive (see Steps 80–81) and mount the holder at the designated position on the vertical optical breadboard.

84. Use the alignment camera to confirm that the beam propagates along R5.

85. Mount the motorized stage (Thorlabs, MTS25/M-Z8) on the vertical optical breadboard.

86. Glue the BK-7 glass wedges to the corresponding holders using optical adhesive (see Steps 80–81), then mount the holders on the vertical breadboard.

87. Rotate HWP5 until most of the laser power is reflected by PBS2.

88. Use the DC Servo Controller/Driver (Newport, CONEX-CC) to tune the tilt and azimuthal angles of PBS2 until the reflected beam propagates parallel to R6.

### 3.10. Power allocation wave plate alignment ⬤ Timing: 30 minutes

89. Mount HWP3 in the rotation mount in front of L9.

90. Rotate HWP3 and measure the depletion beam power transmitted and reflected by PBS2 until the power allocation equals 50:50.

91. Lock the rotation mount using the locking mechanism.

### 3.11. DM, scanners and SLM conjugation alignment ⬤ Timing: 2 hours

92. ▴Critical Set the RM scanning range to approximately 20 μm in the LabVIEW control program. If the beam image appears elongated in the RM scanning direction as a result of the scan motion, the RM surface is not conjugated to the camera plane. Activate the ‘Top Hat’ pattern on the SLM and mark the beam center position as the target in the image with the camera mounted at the designed DM1 surface position. Translate DI2 along R4 until the beam image is not elongated during scanning anymore to conjugate the RM scanning plane and the camera plane. **! Note** Translating DI2 may shift the beam position on the camera; correct the beam shift back to the target by translating LS2.

93. ▴Critical Adjust the conjugation between the RM surface and the hologram on the SLM using the camera mounted at the designed DM1 surface position. Activate the hologram by ‘Holo ON’ on the SLM and set the control voltages of the scanners to 0. Adjust the L4-L5 4f telecentric system along R3 to achieve a sharp hologram image on the camera. Use the piezo pusher (FAB P0013 and FAB P0014, custom-designed) to gently move the lens mounts of the L4-L5 4f telecentric system. Loosen the screws of the lens mounts and push the two lenses in the same direction by the same distance using identical spacers.

94. Use the ‘Live’ image acquisition mode in the nanoscope control software. This functionality enables automatic renewal of imaging within the same FOV, facilitating real-time observation and adjustment. Set the scan range to approximately 20 μm for both directions and confirm that RM scanning is conjugated to the camera plane. The GM1 and GM2 mirrors scan synchronously. Optimize the ‘Galvanometer Scale Factor’ in the ‘Initialize Waveforms V4.vi’ LabVIEW file to minimize the scanning motion of the hologram image on the camera, as demonstrated in **Supplementary Video S1**. This value is crucial for establishing a virtual scanning plane for the galvanometer mirrors that coincides with the scanning plane of the resonant mirror. The optimized alignment is shown in **Supplementary Video S2**.

### 3.12 4Pi interference cavity alignment ⬤ Timing: 1 week

95. Mount HWP4, W1, W2, SH1, SH2, DM1, and DM2 on the vertical optical breadboard in the 4Pi interference cavity according to the CAD model.

96. Secure the half-inch lens mount (FAB-IS0012 and FAB-IS0013, custom-designed for L10 and L12, respectively) at the designated position on the breadboard. Install a Ø0.5” target (*e.g.,* Thorlabs, P1000HW) in the lens mount to take the place of the lens for alignment purpose.

97. Adjust the tip/tilt angles of DM1 until the hologram center precisely propagates through the center hole of the target. Then, remove the target from the lens mount.

98. Glue the Ø1” mirror (M11) onto the actuator (PI, P-841.10) with UV glue and install the actuator on the custom-designed mount (FAB-IS0026, custom-designed).

99. Mount M12, DI3, and QWP4.

100. Remove OBJ1 and install the alignment camera with an iris in front of the it on the upper objective mount (FAB-P0007, custom-designed). Translate the upper objective stage (PI, N-565.260) to monitor the beam with the camera. Adjust the angles of M11 and M12 until the beam hologram stays centered through the iris along the stage translation.

101. Mount L10 in the half-inch lens mount (FAB-IS0013, custom-designed).

102. Mark the beam hologram center on the camera. Mount L11, then translate it parallel to the breadboard and use shims to adjust its height until the beam hologram center is back to the marked position.

103. Translate the upper objective stage until the DM1 surface is imaged onto the camera. This is verified by powering off the SLM and checking the beam pattern on the camera, as illustrated in **Supplementary Figures S5a-b.**

104. Power on the SLM and activate the holograms. Adjust DM1 axially by adjusting its mount (Newport, LP-2A) until the holograms appear sharp on the camera. Verify the conjugation by observing minimal beam scan motion on the camera, as illustrated in **Supplementary Videos S1-S2**. Repeat Steps 103**–**104 as necessary until the conjugations of DM surface, the virtual scanning plane, and SLM hologram are achieved at the same camera plane.

105. The axial translation of DM1 in Steps 103**–**104 affects the co-centering of the beam (hologram) center to L10. To compensate for the misalignment, remove lenses L10 and L11, and repeat Steps 96**–**102.

106. Flip the alignment camera board in its mount and mount it at the lower objective mount, so the camera is facing up to monitor the upper beam. Activate the ‘Top Hat’ pattern on the SLM. Launch the ‘DM Shapes’ software from Boston Micromachines to load the cross shape onto DM1. Adjust the position of DM1 using LP-2A (Newport) to align DM1 surface co-centered with the SLM-hologram, as depicted in **Supplementary Figure S5c-d**. **! Note** The cross pattern applied on DM is invisible at the conjugated plane but becomes visible far beyond the conjugation. For this reason, the camera is mounted in the position of the lower, not the upper, objective mount.

107. Close SH1. Align the lower arm from DM2 to OBJ2 using the method described above for aligning the upper arm of the 4Pi interference cavity (Steps 95**–** 106). Mount the camera on the upper objective mount and have the camera facing downwards to monitor the beam. Use the translation path of the upper objective stage as the reference to align the beam direction of the lower arm to. Mount the following components and align them accordingly: M13, L12, SH2, DM2, M14, L13, DI4, and QWP5.

108. Temporarily remove QWP4 and QWP5. Open both SH1 and SH2. Position a power meter (*e.g.,* Thorlabs, PM130D) behind the top-right side of PBS2. Rotate HWP4 to minimize the measured power value.

109. Re-mount QWP4 and QWP5. Adjust their orientations iteratively until the power measured by the power meter is maximized.

110. Remove the power meter. Extend R6 towards the top right direction. Mount an alignment camera assembly on the extended rail.

111. Use the alignment camera to locate the beams. If two spots are observable, adjust M13 until the spots coincide. Then, translate the camera to the far end of the rail and overlap the spots again by adjusting M14. Repeat this step for several rounds until both spots are consistently overlapping along the rail. Once completed, remove the camera assembly and the extended rail.

112. Rotate HWP3 in front of PBS2 to achieve an approximate 50:50 power allocation between the two arms of the 4Pi interference cavity.

### 3.13. Objective alignment ⬤ Timing: 3 days

113. Mount both objectives in their corresponding lens mounts. **! Note** the objectives for isoSTED nanoscope are Olympus UPLSAPO100XS silicone oil objectives, which feature correction collars to compensate for spherical aberrations induced by the cover glass. Adjust this ring as needed. In our case, it is calibrated for 170-μm thick cover glasses.

114. Prepare a gold nanoparticle sample as described in ‘**Test sample preparation ⬤ Timing: 3 days**’.

115. Install L14 and mount the PMT at the designed position. Add an iris and a neutral density filter (*e.g.,* Thorlabs, ND20A) in front of the PMT to reduce background noise.

116. Activate the blazed-grating hologram on the SLM.

117. Apply a drop of silicone oil (Olympus, SIL300CS) to OBJ2. Place the sample between the two objectives on the sample holder. Close SH1. Use the PMT to image the sample with OBJ2. Adjust the *z* position of the sample until the beads are in focus.

118. Apply a drop of silicone oil to the top cover glass. Open SH1 and close SH2. Lower OBJ1 using the upper objective stage piezo to get the same gold nanoparticles in focus. **! Note** If there is no overlap between the FOVs of OBJ1 and OBJ2, adjust the adjustment screws (Newport, AJS100-0.5H-NL) underneath the lower objective stage to coarsely overlap the FOVs. Further fine adjustments to achieve perfect FOV overlap can be made by translating the piezo stage P-612.2SL (PI).

### 3.14. Focus-lock module alignment ⬤ Timing: 4 hours

119. Install L20 in the *xy* translation mount following the CAD model as shown in **Fig. 14**. Adjust its axial position to collimate the 980-nm near-infrared (NIR) laser beam of the laser diode LH3.

120. Merge the NIR beam with the other beams via dichroic mirror DI3 in the top arm of the 4Pi interference cavity. Adjust M20 and M21 so that the 980-nm beam hits the back aperture center of the upper objective.

121. Let the NIR beam propagate through both objectives and separate it from the common beam path by dichroic mirror DI4 in the lower arm of the 4Pi interference cavity.

122. Focus the NIR beam onto the camera C using L21. Adjust M24 and M25 until the beam hits the center of the camera.

123. Insert the cylindrical lens CL between C and L21 to introduce astigmatism to the focus so that the focus is elliptical when the focal planes of the two objectives no longer coincide.

124. Translate the lower objective stage in both axes directions with the nanoscope control software and observe the direction the focus is moving in the camera image. Rotate C until the focus moves horizontally and vertically in the image when translating the lower objective stage.

125. Activate the focus-lock module (**Fig. 19**) in the nanoscope control software. Once alignment of the two objectives is achieved, click the ‘Locked’ button to register the position. Subsequently, the ‘Drift’ indicator turns yellow when the two objectives are misaligned from the locked position. The ‘Control Option’ button located below the ‘Locked’ button can toggle between ‘Manual’ and ‘Automatic’ misalignment compensation modes.

**Fig. 19.**
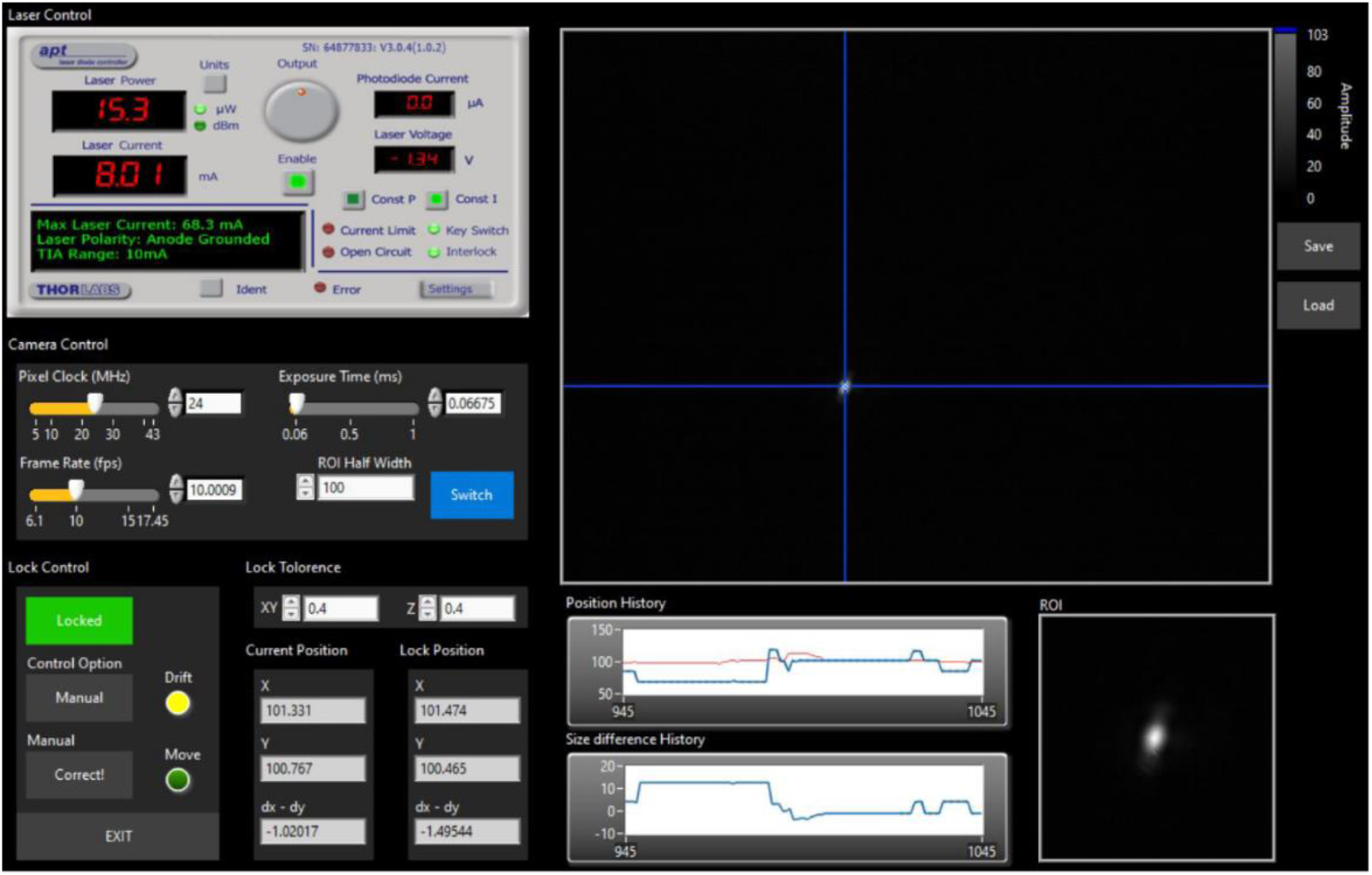
User interface of ‘Focus Lock’ control panel.

### 3.15. Test sample preparation ⬤ Timing: 3 days

126. Clean a 25-mm-diameter round precision cover glass (Thorlabs, cat # CG15XH) in 1 M KOH in a sonic bath for 15 minutes.

127. Rinse the cover glass three times with filtered water.

128. Rinse the cover glass in 100% ethanol.

129. Let the cover glass dry at room temperature.

130. Apply 200 μL 1 mg/mL BSA-biotin solution diluted in PBS to the cover glass and let it sit for 5 minutes.

131. Remove the solution and rinse three times with PBS.

132. Apply 200 μL 1 mg/mL neutravidin solution diluted in PBS. Let it sit on the cover glass for 5 minutes.

133. Remove the solution and rinse three times with 10 mM MgCl_2_ solution diluted in PBS. **! Note** This step is only required for using the GattaQuant beads, but not with other fluorescent beads described below.

134. Prepare beads solution containing: Gold Nanoparticles (Sigma-Aldrich, cat # 742058-25 mL, 150-nm diameter), 1:8 dilution; 10 mM MgCl_2_, final concentration; GattaQuant Bead R (Atto 647N conjugated, 23-nm diameter), 1:4 dilution. **! Note** We recommend making four samples, one with GattaQuant beads (for resolution measurement), as described, and, for alignment purposes, one each with crimson beads, red beads, and yellow-green beads replacing the GattaQuant beads. For the latter three, no MgCl2 is required, and we recommend dilutions of 1:10,000. If stored in the dark at room temperature with ProLong Diamond mountant, these samples can be used for a few months.

135. Apply 20-μL of the beads solution to the cover glass and let it sit on the glass for 5 minutes.

136. Remove the solution. Do not rinse.

137. Place the glass beads-side up in the customized sample holder.

138. Add a drop of ProLong Diamond Antifade Mountant on the beads side of the cover glass.

139. Apply another 25-mm round cover glass on top of the beads side of the prepared cover glass and then press it down gently.

140. Allow the sample to sit in the dark at room temperature for 48 hours.

141. Secure the cover glass sandwich to the customized sample holder using two-component silicone adhesive.

### 3.16. Detection path alignment ⬤ Timing: 1 day

142. Turn on LH1. Turn on LH2 and set its output power to 15%. In the nanoscope control software, set the 650-nm line approximately to the maximum.

143. Rotate HWP3 to achieve a roughly equal allocation of the excitation laser power between the upper and lower arms of the 4Pi interference cavity.

144. Insert a sample of crimson beads and gold nanoparticles between the two objectives. Image the gold nanoparticles with the PMT. Toggle between illuminating the sample with either the excitation or the depletion beam at power levels that create sufficient signal. In the nanoscope control software, optimize the ‘Tip’, ‘Tilt’, and ‘Defocus’ coefficients of the holograms displayed on the SLM to ensure the co-centered overlap of the depletion PSFs with the excitation ones in three dimensions (3D).

145. Image the beads with the upper and lower objectives sequentially to ensure that the two objectives image the same FOV. Replace the ND filter in front of the PMT with the bandpass filter to detect fluorescence from the crimson beads.

146. Activate the focus-lock module to maintain locked objective positions throughout the entire alignment procedure. This ensures that the objectives remain in their relative positions, providing stable and consistent imaging conditions.

147. Block the PMT with a cap to prevent over-exposure.

148. Switch the illumination beam from the excitation beam to the STED*_xy_* depletion beam. Set the depletion beam power to maximum. **! Note** Since a small portion of the depletion laser power can leak into the detection module due to the imperfect reflection of DI2 at 775 nm (**Supplementary Fig. S7**), we can use this leaked light for the alignment of the detection path.

149. Install the three rails (R8, R9, and R10) for the detection module on the base plate. Each rail corresponds to one detection channel. **! Note** Since the thick glasses of DI5 and DI6 result in a noticeable shift in the lateral direction when the beam passes through them, it is important to align the channels in the following order: (i) APD1 for the green channel, (ii) APD2 for the orange channel, and (iii) APD3 for the far-red channel.

150. Open both SH1 and SH2.

151. Mount DI5 on R8. Translate the DI5 mirror mount to roughly align the beam to hit the center of the front surface of DI5. Adjust the angle of the mount to ensure the beam is reflected parallel to rail R8. Translate the mount to center the reflected beam above R8. **! Note** It’s possible that two or more spots are observed with the alignment camera since the uncoated surface of the dichroic mirror can create ghost reflections. In this case, use only the brightest spot as the reference for alignment.

152. Mount L17 on R8. Translate L17 in 2D to align its center with the beam (analog to **Box 3**).

153. Connect the fiber head adapter (Thorlabs, SM1FC) with the fiber (Thorlabs, FG105LCA + FT05SS) and mount it on R8. Position a camera close to the other end of the fiber. Translate the adapter in 3D to maximize the brightness of the spot observed on the camera.

154. Turn off the depletion beam and turn on the excitation beam. Confirm that the sample is still in focus by imaging crimson beads with the PMT channel. If not, adjust the sample stage position. Repeat the brightness maximization of Step 153.

155. Mount the two bandpass filters F4 in a lens tube (Thorlabs, SM1S35), maintaining a gap of ∼50 mm between them to avoid possible interference. Screw the lens tube to the front of the fiber head adapter mount. This will reduce environmental light detection.

156. Remove the camera and connect the fiber end with APD3. Image the crimson sample with APD3 while gradually increasing the excitation laser power until the beads can be detected by the APD. Translate the fiber head adapter in 3D to maximize the photon count. ▲Critical Start from low laser power and increase gradually to avoid saturation and potential damage of the APDs.

157. Perform the same procedure for the orange (APD2) and far-red (APD3) detection channels using the corresponding optomechanical components and samples with yellow-green and red beads (Thermo Fisher, F8803 and F8801), respectively.

### 3.17. Setting up pulse gate/delay module ⬤ Timing: 1 hour

158. Insert a fluorescent bead sample between two objectives. Configure the system to use only one objective for imaging by opening one shutter (SH1 or SH2) while closing the other. **! Note** The system comprises three pulse gate/delay modules, each associated with a specific detection channel. Therefore, the selection of fluorescent beads should be based on their emission wavelengths, aligning with the passing band of the detection channel to be set up.

159. Turn on the excitation beam for imaging and move the beads into focus. On the ‘Timing’ control panel (**Fig. 20**) in the nanoscope control software, set ‘Window Width (ns)’ to a small value (∼0.1 ns). In continuous imaging mode, maximize the peak intensity observed in the collected images by adjusting the ‘Window Delay (ns)’.

**Fig. 20.**
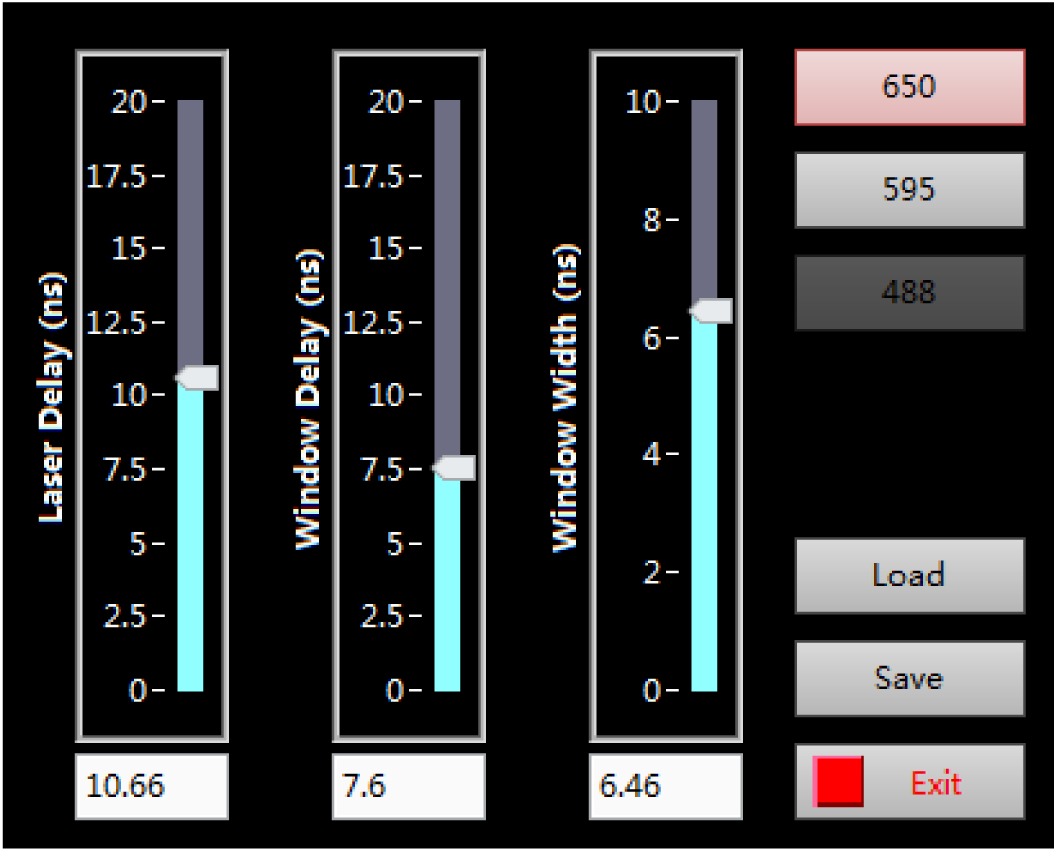
User interface of ‘Timing’ control panel.

160. Activate the blazed-grating hologram on the SLM. Turn on both excitation and depletion beams for imaging. Set ‘Window Width (ns)’ to its maximum value at 10. Minimize the fluorescence signal observed in the collected images by adjusting the ‘Laser Delay (ns)’ value. **! Note** It requires more than 60 mW depletion laser power measured at the back aperture of the objective to observe a substantial STED effect on crimson beams.

161. Set the ‘Window Width (ns)’ value to 5. Activate the ‘Vortex’ hologram on the SLM control panel of the nanoscope control software. Turn on both excitation and depletion beams for imaging. Fine tune ‘Laser Delay (ns)’ and ‘Window Delay (ns)’ values (±0.5 ns) as needed for the visually judged best resolution and contrast. The optimized ‘Window Delay (ns)’ value is expected to be ∼1 ns larger than the value found in Step 159.

162. Adjust the ‘Window Width (ns)’ value to maximize the contrast observed in the collected images. **! Note** The optimal ‘Window Width’ value is expected to be roughly the fluorescence lifetime of the imaged fluorophore.

163. Save these settings for future use. **! Note** The optimal combination of values depends on the spectral properties of the imaged fluorophores and therefore can vary when switching to a differently labeled sample.

164. Repeat this procedure for the other two channels.

### 3.18. Setting up and operating the AO module ⬤ Timing: 2 days

165. Read ‘**Aberration correction**’ for instructions. **! Note** The following steps describe the initial set up of the AO module, first correcting for system aberrations and then for sample-induced aberrations. Some of these steps will be revisited during daily operation of the instrument while others are only required during the initial setup. Additionally, we describe the procedure for the correction of aberrations in thick samples.

166. To correct the system aberrations caused by imperfections of the optical components and their alignment, turn on LH2 for excitation and image 100-nm-diameter crimson fluorescent beads as the target (refer to ‘**Initial test for sample imaging ⬤ Timing: 2 hours**’ for details). Confirm that the objective correction collar is set to the thickness of the cover glass. Rotate HWP6 to re-allocate a portion of the excitation laser power to the lower arm of the 4Pi interference cavity. Perform image-based aberration correction to optimize the Zernike coefficients of DM1 (‘Upper DM’ tab) and DM2 (‘Lower DM’ tab) by opening only one shutter at a time. To achieve this, open the ‘Deformable Mirrors’ control panel (**Fig. 21a**) and click the ‘Scan Modes’ button. In the subsequent popup panel (**Fig. 21b**), choose a single Zernike mode (*e.g.*, #7 for ‘horizontal coma’) for optimization. The nanoscope control software will automatically apply a series of different incremental corrections for the ‘horizontal coma’ coefficient while recording the corresponding value of the metric at each step. The control software will subsequently fit a Gaussian or quadratic function to the measured values of the metric (image brightness). The optimal correction value is determined as the value that maximizes the fitted function. Repeat this measurement for each required Zernike mode (typically, #5 to #11 and #22). Perform multiple sequential optimization cycles, covering all necessary Zernike modes, to achieve optimal aberration correction. Typically, 2-3 cycles are sufficient. Save these settings.

**Fig. 21.**
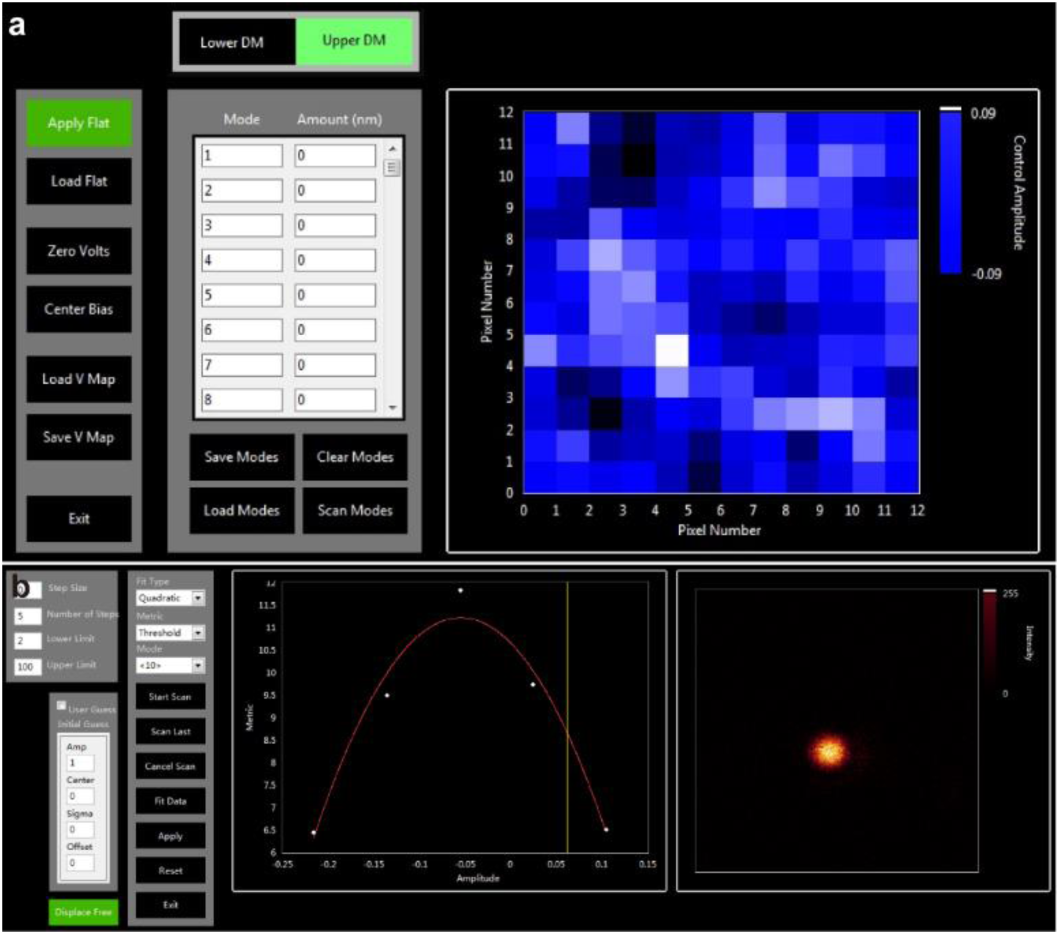
‘Deformable Mirror’ control interface. **a**, Main control panel. **b**, ‘Scan Modes’ selection pop-up.

167. Rotate HWP6 back to its original orientation and turn off LH2 after finishing this step.

168. Turn on LH1 for depletion. Display a ‘Top Hat’-shaped hologram on the SLM. Proceed with imaging the 150-nm diameter gold nanoparticles utilizing the *xz* scanning mode. Manually adjust each Zernike coefficient via the ‘SLM’ panel within the nanoscope control software until achieving a PSF with a central minimum approaching zero intensity and resembling the one shown in **Fig. 7b–c**. **! Note** Correcting system aberrations (Steps 166**–**168) is not a routine operation. It should only be performed during the initial system implementation or when any key optical components are re-aligned.

169. When imaging thick samples which induce large amounts of aberrations caused by the refractive index mismatch between sample, embedding medium and immersion liquid (silicone oil), proceed as follows: Image 150-nm diameter gold nanoparticles attached to the bottom coverslip of the thick sample using the depletion beam for illumination and the PMT for detection. Repeat Step 166 to optimize the Zernike coefficients for each DM individually by maximizing the brightness as the metric. To reduce noise contributions, apply a background pixel threshold by changing the ‘metric’ setting to ‘Threshold Mean’ in the popup panel (**Fig. 21b**). Adjust the ‘Lower Limit’ and ‘Upper Limit’ values to define the desired background threshold. These specific threshold values should be determined empirically based on the signal-to-noise ratio (SNR) observed in the acquired images.

170. In preparation for minimizing remaining aberrations, align and lock the objectives (refer to ‘**Objective alignment** ⬤ **Timing: 3 days**’ and ‘**Focus-lock module alignment** ⬤ **Timing: 4 hours**’ for details).

171. Turn on both LH1 for excitation and LH2 for depletion. Translate the sample to the region of interest. Activate the ‘Vortex’ hologram on the SLM for 2D STED imaging. Choose a weak depletion power (∼10% of the power level used for ordinary imaging) to limit photobleaching.

172. Compensate for the local sample-induced aberrations by simultaneously adjusting both DMs. ‘Scan Modes’ (**Fig. 21**) for automated DM adjustment is used in general while the manual adjustment is more reliable when the image SNR is less than 2. Proceed to optimize the 4Pi mode coefficients^16^ sequentially, using the image brightness as the metric for assessment. This iterative process is repeated as needed to ensure effective correction of aberrations induced by the sample. **! Note #1** This step is optional when imaging relatively thin cell samples, particularly if the imaging volume is close to the coverslip. However, for thick tissue samples, it becomes essential to compensate for local sample-induced aberrations to enable high-quality isoSTED imaging. **! Note #2** For manual compensation of sample-induced aberrations in isoSTED imaging, initiate live imaging in the nanoscope control software. Under the ‘Upper DM’ tab, introduce a small value (e.g., +0.1) into the second column of a chosen Zernike mode row (*e.g.*, #7 for ‘horizontal coma’). Repeat this adjustment with the identical value in the corresponding cell under the ‘Lower DM’ tab. Iterate with varying values to maximize photon counts. Subsequently, apply a small positive offset (e.g., +0.1) to the ‘Upper DM’ value and a corresponding negative offset (e.g., -0.1) to the ‘Lower DM’ value, again optimizing for maximum photon counts. Repeat this process for all relevant Zernike modes. Perform two to three sequential optimization cycles encompassing all necessary 4Pi modes to achieve optimal correction. Note that three specific 4Pi modes^16^ (*Q*_1_^−1^, *Q*_2_^+1^, and *Q*_3_^+1^) induce image shifts rather than aberrations, and thus require no adjustment in this step.

173. To establish bias values for 3D imaging of thick samples, begin with using the 2D isoSTED mode with a weak depletion power, set at approximately 10% of the power used for standard imaging. Next, optimize the 4Pi mode with the ‘z Comp’ panel in the nanoscope control software (**Fig. 17**), including ‘Piston’ for fringe-shift, ‘Defocus’, and ‘Spherical’ for the first-order spherical aberration, to manipulate the interference fringes underneath the PSF envelope until the effective PSF size is unchanged (See more details in Ref. [12] and Extended Data Fig. 3). Repeat this measurement every 1 μm in the *z* direction. Once a series of bias values corresponding to different depths is obtained, calculate the step size based on linear regression analysis applied to these discrete points. Activate the bias setting function and set the step sizes in the nanoscope control software. **! Note** Imaging large 3D volumes (>1 μm in the *z* direction) in thick tissues (>30 μm) necessitates this step.

### 3.19. Initial test for sample imaging ⬤ Timing: 2 hours

**! Note** The first time the steps in this section will be executed, the image size (scan range) has not been calibrated yet. The calibration steps are described in the following Section 3.20.

174. Install a sample to be imaged between two objectives.

175. Load the settings of pulse gate/delay modules from previously saved data.

176. Align the objectives following the procedure outlined in ‘Objective alignment ⬤ Timing: 3 days’.

177. Compensate for the sample-induced aberrations following the procedure outlined in Steps 169**–**173.

178. Lock the positions of the objectives following the procedure outlined in ‘**Focus-lock module alignment ⬤ Timing: 4 hours**’.

179. Use the ‘Image Acquisition Parameters’ panel (**Fig. 22**) within the nanoscope control software to preliminarily image the sample at relatively low laser power. Translate the sample to the region of interest. Configure the imaging parameters, *e.g.,* scan mode, image dimensions, step size and line accumulation values to assess imaging quality and performance without significantly bleaching it. Repeat the process outlined in Step 177 for local aberration compensation.

**Fig. 22.**
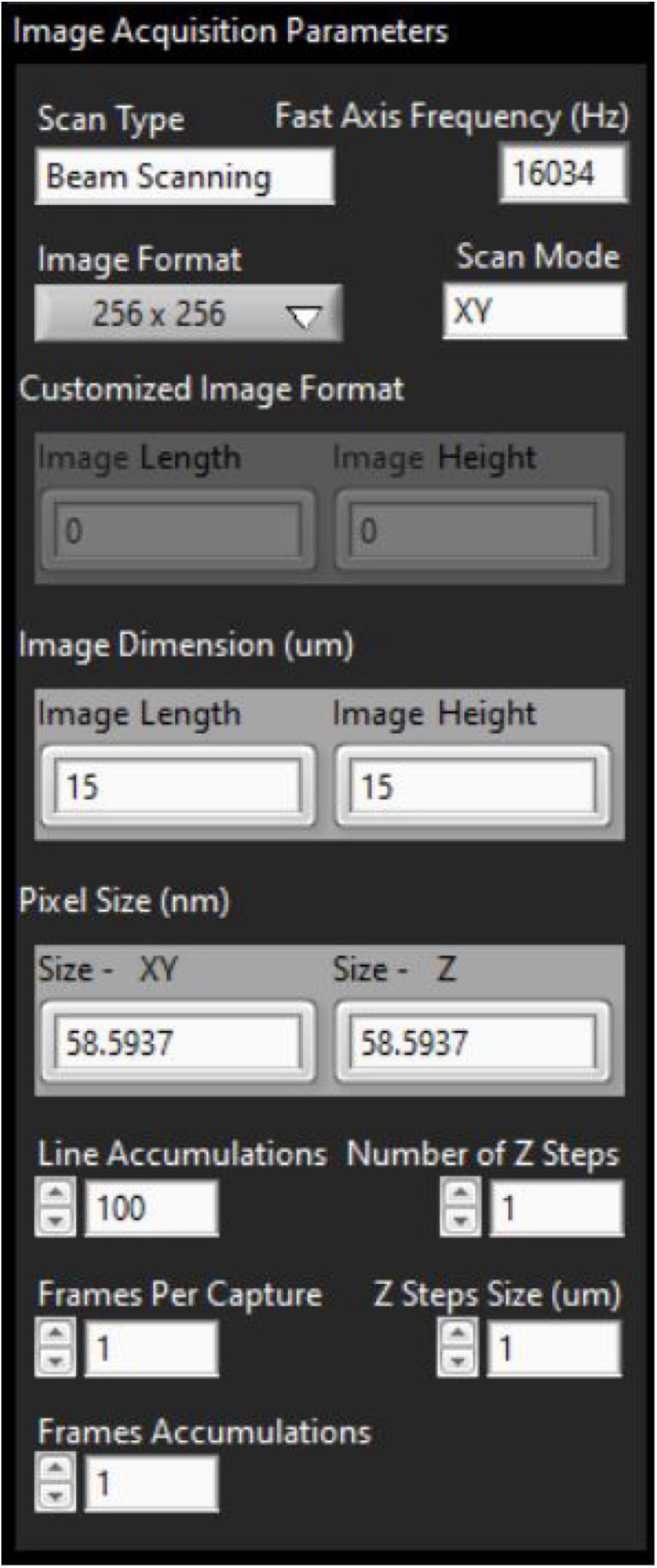
User interface of ‘Image Acquisition Parameters’ control panel.

180. Set the depletion laser power above 60 mW (measured at the back aperture of the objective) for high-resolution STED imaging. Typical parameters for high-quality isoSTED images are 10 × 10 µm image dimensions, 1024 × 1024 pixels, and 100 line accumulations.

181. Enable the preferred detection channel(s) by selecting them in the ‘Detectors’ panel and activate the corresponding laser(s) via the ‘Laser Controls’ panel. Click the ‘Live’ button to initiate live imaging mode.

182. To record data for later analysis, set the desired file path and click the ‘Capture’ button.

### 3.20. Image size (scan range) calibration ⬤ Timing: 2 days

183. Mount the Ronchi ruling (Edmund Optics, 38-566, 600 lp/mm, period length *T* = 1.667 μm). Open SH1 and close SH2 to utilize OBJ1 for imaging the sample with the PMT. Illuminate the sample with the STED*_xy_* beam configuration. Acquire images at a format of 1024×1024 pixels.

184. ▴Critical Calibrate the resonant mirror first. Carefully rotate the gratings to align the periodic structures parallel with the scanning direction of RM. **! Note** Scan a large range (e.g., 20 μm) to determine and adjust the orientation of the gratings as precisely as possible.

185. Apply different control voltages (*V*) to RM to acquire images of the Ronchi ruling grating at varying sizes of the FOV. Plot the values as shown in **Fig. 23a** to depict the relationship between the control voltage and the corresponding true image size measured by the Ronchi ruling. **! Note** The correlation between the control voltage and the image size becomes increasingly imprecise for smaller image sizes (<5 μm).

**Fig. 23.**
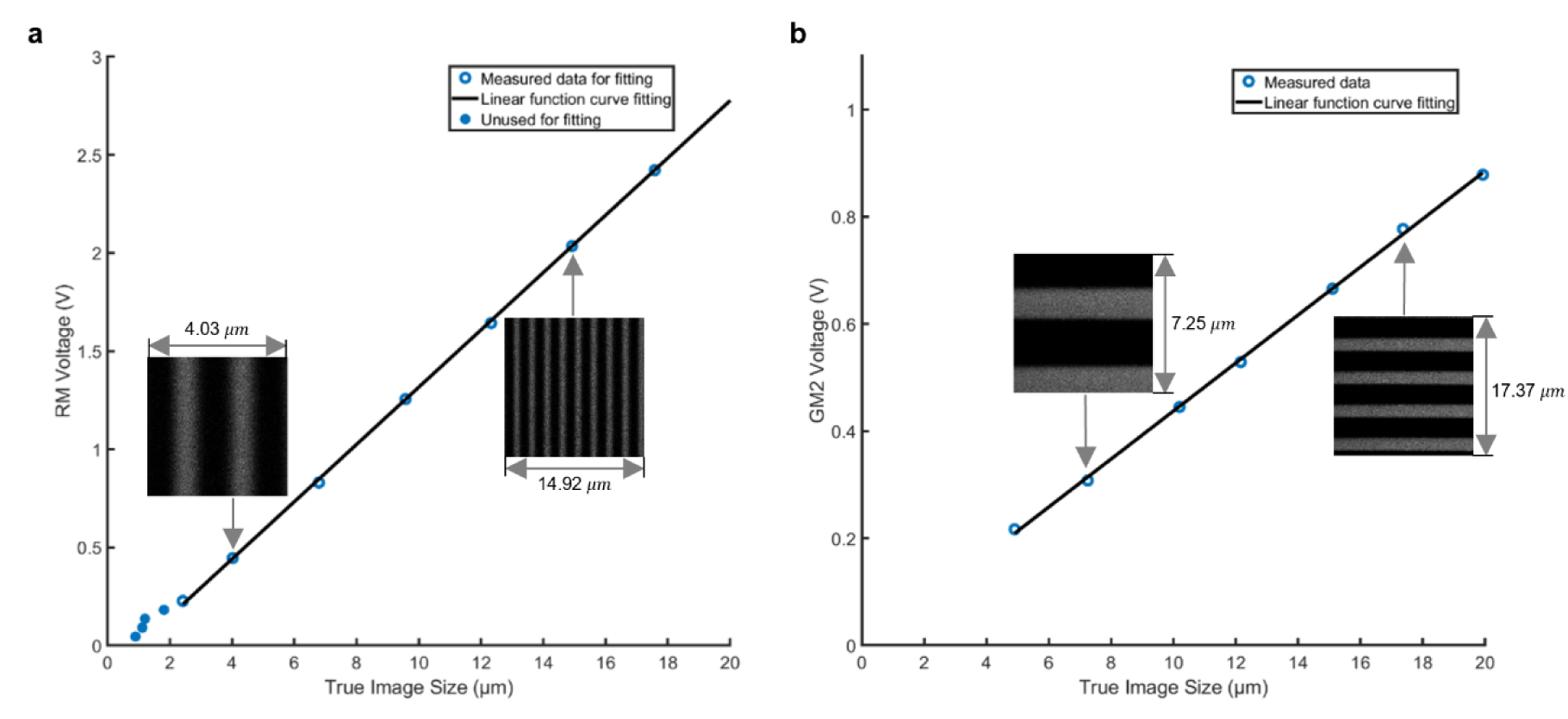
Image size calibration as a function of control voltage on scanning mirrors. **a**, resonant mirror. b, Galvanometer mirror. The image size is measured with Ronchi ruling gratings, and linear curve fitting is applied to identify the control parameters in the LabVIEW program.

186. Fit the raw data using a linear function *y* = *ax* + *b*. In the example shown in**Fig. 23a**, *a* = 0.146 and *b* = −0.1441.

187. Update the fitting parameters (*a* and *b*) in the ‘Re Mirror um to Volts.vi’ in the folder ‘Control Software > Host VIs > General Sub Vis’.

188. In preparation for calibrating the galvanometer mirrors, rotate the Ronchi ruling by 90°. **! Note** We opted to utilize a second Ronchi ruling grating (Edmund Optics, 66-346, 240 lp/mm), because the first one could not be mounted properly after undergoing a 90° rotation).

189. Repeat Steps 185-186 to obtain the fitting parameters for the galvanometer mirrors. In the examples shown in **Fig. 23b**, fitting with the function *y* = *ax* + *b* yields the *a* = 0.04481, *b* = −0.01072.

190. Update these parameters in the sub-VI called ‘Galvanometers um to Volts.vi’ in the folder ‘Control Software > Host VIs > General Sub Vis’.

### 3.21. Resolution measurement (bead samples) ⬤ Timing: 1 day

191. Acquire Gatta quant beads images referring to Section ‘**Initial test for sample imaging** ⬤ **Timing: 2 hours**’. Create a new folder and transfer the raw images with DBL file extension.

192. Open the analysis software named ‘exe.m’ written in MATLAB, located within the folder ‘resolution-quantification-beads’ (code available at https://zenodo.org/uploads/10574469).

193. Type ‘ROIr’, ‘pixelSize’, and ‘beadr’ in lines # 8, 9, and 10, respectively. ‘ROIr’ should be appropriate to crop a single bead. ‘pixelSize’ and ‘beadr’ represent the dimension of each pixel size and the diameter of the bead, respectively.

194. Run ‘exe.m’ and select the folder which includes bead data.

195. When an image is opened, proceed to click the center of each bead in the image. Press the enter key to finalize the selection.

196. Repeat Step 195 for all data.

197. The software will automatically export the resolution after the analysis.

### 3.22. Resolution measurement (biological samples) ⬤ Timing: 1 day

198. Save two different images with the same FOV sequentially.
199. Open ImageJ (National Institute of Health) and open two saved images to perform Fourier ring correlation.
200. Run the plugin following Plugins>BIOP>Image Analysis>FRC>FRC calculation.
201. Select two images and ‘Fixed 1/7’ for resolution criteria^18^. (Optionally, the method provided in Ref. [19] is recommended for resolution measurements with tubular biological structures).

## 4. User Operation

After completing the above steps, the isoSTED nanoscope should be properly aligned and calibrated and ready for biological applications. Users primarily operate the instrument through the nanoscope control software (the user interface is shown in **Fig. 17**). Steps 174-182 in Section ‘**Initial test for sample imaging ⬤ Timing: 2 hours**’ describe the standard operation procedure users should follow to image their samples.

To obtain images that fully take advantage of the optical resolution of the isoSTED nanoscope, the pixel size should be smaller than half of the expected resolution, in accordance with the Nyquist sampling theorem. The ‘Pixel Size (nm)’ is determined by the ‘Image Format’ and ‘Image Dimension (μm)’ settings in the ‘Image Acquisition Parameters’ control panel. In 3D *z*-stack imaging mode, it is additionally required to set the ‘Z Steps Size (μm)’ to a value smaller than half of the expected axial resolution. Based on our experience, optimal isoSTED imaging performance is achieved with a ‘Pixel Size (nm)’ and ‘Z Steps Size (μm)’ equal to 10 nm.

## 5. Troubleshooting

In complex optomechanical systems such as this isoSTED nanoscope, challenges are manifold and range from suboptimal resolution to a non-functional nanoscope. **Table 1** lists commonplace issues, but it is impossible to anticipate every potential problem. We invite our readers to contact us should they run into obstacles.

**Table 1.** Troubleshooting table.

| Problem | Possible reason | Solution |
| --- | --- | --- |
| Very low or no signal levels | The shutters in front of the DMs are closed. | Open the shutters. |
|  | The detection window width is set to 0. | Set the detection window width to greater than 3 ns. |
|  | Poorly labeled sample | Check the sample with a different microscope. |
|  | Lack of immersion oil | Add immersion oil until the droplet fills the gap between the objective and the specimen. Make sure no air bubble is introduced in the oil. |
|  | Poor cable connections. | Check the electronic connections. A frequently overlooked place is the real-time system integration (RTSI) bus between the two FPGA boards. |
| Heavily distorted PSF | Malfunction of DM(s) | Regular check on the membrane is encouraged. A quick check can be done by inspecting the beam reflected off the DM when the depletion laser is on. If one or two actuators are damaged, the illumination becomes non-uniform, which is observable directly by eye or using a camera. A thorough inspection is possible by re-calibrating the DM offline using a Twyman-Green interferometer (see section 'SLM and DM calibration' for details). |
|  | Beam misalignment | Misalignment of any of the beam paths disrupts the conjugation between key devices (SLM, resonant mirror, and DMs) with the back pupil of one or both objectives. A rapid assessment can be conducted by substituting the objective with a camera and capturing an image of its back pupil. Small misalignments can potentially be addressed by directly shifting the hologram pattern on the SLM. Larger misalignments require manual correction. |
| High background | Open instrument enclosure | Close the enclosure during imaging. |
|  | Incorrect polarization states of the depletion beam in the 4Pi interference cavity. | Rotate the wave plates inside the 4Pi interference cavity to allow the depletion beam leave the cavity from the empty (top-right) side of PBS2 after passing through both objectives<sup>12</sup>. |
| Low fluorescence intensity | Aberrated PSFs | If only the depletion PSF is affected, utilize the SLM for correcting the systematic aberrations in the depletion beam path. In cases where the excitation PSF is also impacted, opt for the use of DMs. |
|  | The sample is out of focus. | Translate the sample until it is in focus. |

## 6. Timing

We provide an approximate timeframe below for essential tasks during setup. Notably, the actual duration may vary depending upon part availability, as well as the experience of the personnel involved. Given the complex nature of the system, a generous allotment of time is advised for troubleshooting and potential repetition of tasks and to correct for errors in machined components or replace damaged elements. With careful planning and a focused effort, a collaboration between a biophysicist or optical engineer for instrument setup and biologists for specimen, isoSTED images can be acquired within 12 months. Upon gaining proficiency in constructing the nanoscope, coupled with the availability of all required components, the instrument can be set up in about 2-3 months!

Reagent setup, sample preparation: 1 hour for fluorescent bead samples, 1–2 days for cell samples, and 1 week for tissue samples.

Equipment setup, room preparation: 1 week to 2 years based on our experience, contingent upon facility considerations. This duration is highly variable and is intricately tied to the availability of suitable space and responsiveness of the local facilities department to fix problems.

Equipment setup, lead times on components: Certain pivotal components, such as paired objective lenses and deformable mirrors (DMs) often entail lead times of several months. Early ordering of these components is advisable to circumvent potential delays.

Equipment setup, software installation and familiarization: control and data collection software: 1 week; other custom code: 3 days

Steps 1**–**4, software installation and setup: 1 day

Steps 5**–**6, optomechanical assembly: 1 week

Steps 7**–**8, electronics assembly: 1 week

Steps 9**–**15, AOM setting and alignment: 2 hours

Steps 16**–**23, polarization separation module alignment: 4 hours

Steps 24**–**41, SLM installation and alignment: 1 day

Steps 42**–**62, excitation path alignment: 3 days

Steps 63**–**74, scanner installation and alignment: 1 day

Steps 75**–**88, PBS installation and angle alignment: 3 hours

Steps 89**–**91, power allocation wave plate alignment: 30 minutes

Steps 92**–**94, DM, scanners, and SLM conjugation alignment: 2 hours

Steps 95**–**112, 4Pi interference cavity alignment: 1 week

Steps 113**–**118, objective alignment: 3 days

Steps 119**–**125, focus-lock module alignment: 4 hours

Steps 126**–**141, Test sample preparation: 3 days

Steps 142**–**157, detection path alignment: 1 day

Steps 158**–**164, setting up pulse gate/delay module: 1 hour

Steps 165**–**173, setting up and operating AO module: 2 days

Steps 174–182, initial test for sample imaging: 2 hours

Steps 183**–**190, image size (scan range) calibration: 2 days

Steps 191**–**197, resolution measurement (bead samples): 1 day

Steps 198**–**201, resolution measurement (biological samples): 1 day

## 7. Anticipated results

The example presented here illustrates the expected performance of the system. When it is fully functional, the isoSTED nanoscope enables users to image fine subcellular structures in single or multi-color mode in cells and in tissues, for example, the Golgi apparatus in HeLa cells (**Fig. 24a**). If a sufficiently high 3D resolution is reached, the layered structure of the Golgi apparatus becomes clearly visible, and the cross sections at different depth are distinct (**Fig. 24b–d**).

**Fig. 24.**
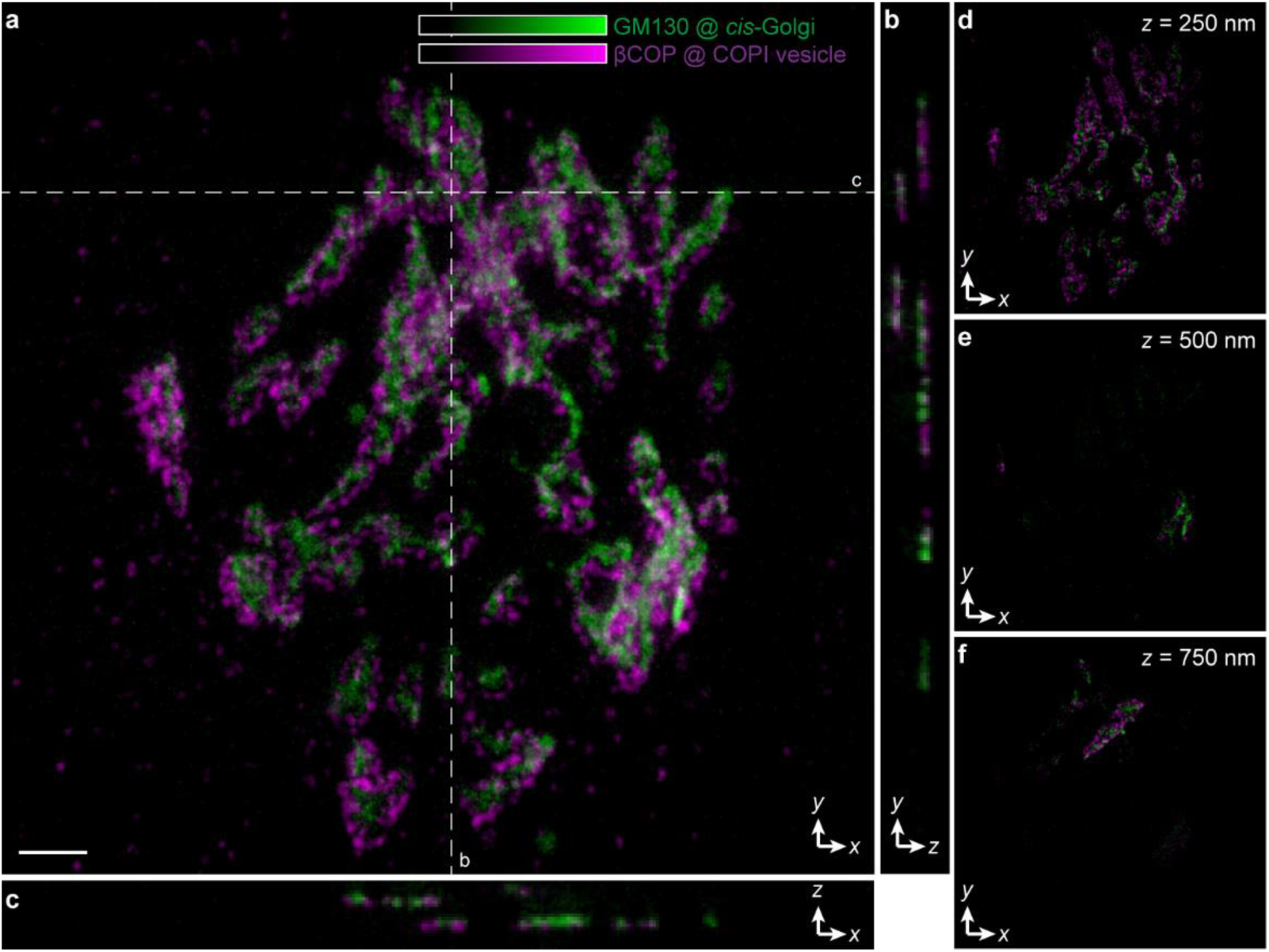
Example isoSTED images of coat protein complex I (COPI) vesicles (magenta) and Golgi apparatus (green) in a HeLa cell visualized by immunolabeling βCOP with ATTO 647N and GM130 with ATTO 594. **a**, Projection image in *z* direction. **b–c**, Cross sections at the positions denoted by the dashed lines in **a**. **b–d**, *xy* cross sections at different depths (z). Scale bar: 1 μm.

### 7.1. Reporting Summary

Further information on research design is available in the Nature Research Reporting Summary linked to this article.

### 7.2. Data availability

The CAD file repository (Dassault Systèmes SE, SolidWorks) can be found at https://zenodo.org/uploads/10574469. The complete part list is available as **Supplementary Table 1**. Datasets generated and/or analyzed in this study are available from X.H. or J.B. (the corresponding authors) upon request.

### 7.3. Code availability

The control and data collection software (National Instruments, LabVIEW) can be obtained from the Zenodo repository at https://zenodo.org/uploads/10574469 or by requesting it from the corresponding authors X.H. or J.B. The source code for SLM calibration is accessible from https://github.com/Hao-Laboratory/Diffraction-based-SLM-calibration, and the source code for DM calibration is found at https://github.com/jacopoantonello/dmlib. For other functions like data transformation and resolution quantification, the source code can be downloaded from https://zenodo.org/uploads/10574469.

## 8.1. Acknowledgements

We thank Jacopo Antonello for advice, discussions and help with adaptive optics. X.H. and S.T. were supported by the Zhejiang Provincial Natural Science Foundation of China (LR25F050002). J.B. acknowledges support by NIH grant U54 DK106857.

## 8.2. Author contributions

X.H., E.S.A. and D.-R.L. designed the instrument. Y. L., D.-R.L. and X.H. built the instrument. X.H., E.S.A., D.-R.L. and Y.L. developed the software. Y.L., D.-R.L. and S.T. calibrated, optimized, and tested the instrument. L.K.S., D.-R.L. and Y.L. designed and optimized the sample preparation protocols. X.H. and J.B. supervised this project. All authors contributed to writing the manuscript.

