## Supplementary File for "Implementation of an adaptive-optics assisted isoSTED nanoscope"

### Implementation of an isoSTED nanoscope

**Supplementary Note 1 | Quantification of microscope resolution in the presence of finite-sized objects.**

To assess the resolution of the microscope, fluorescent bead samples were imaged. The average full width at half maximum (FWHM) of the beads was determined by fitting a Gaussian or Lorentzian function to the intensity profile of multiple isolated beads. However, when the microscope's resolution approaches the bead size, the resulting FWHM of the bead images exceeds the width of the point spread function (PSF). This disparity arises from the convolution of the effective PSF with the object (in this case, a sphere with a specified diameter). While deconvolution offers a comprehensive approach to determine the PSF, it necessitates computationally intensive post-processing steps. Here, we propose a simple equation to estimate the FWHM of the effective PSF. We substantiate its validity through mathematical proof in one dimension, with applicability extending to two-dimensional images without loss of generality.

The PSF ($w(x)$) and an isolated (sparse) fluorescent bead ($o(x)$) can be approximately expressed as

$$w(x) = e^{-\frac{x^2}{2\sigma_{\mathrm{PSF}}^2}} \qquad \text{Eq. S1}$$

and

$$o(x) = e^{-\frac{x^2}{2\sigma_{\mathrm{bead}}^2}}, \qquad \text{Eq. S2}$$

where $\sigma_{\text{PSF}}$ and $\sigma_{\text{bead}}$ are the sizes of the effective PSF and the bead, respectively. The image of the bead ($v(x)$) is therefore a convolution of two Gaussian functions.

$$v(x) = w(x) \otimes o(x). \qquad \text{Eq. S3}$$

Based on the property of Gaussian convolutions, $v(x)$ is also a Gaussian function:

$$v(x) = \sqrt{2\pi} \cdot \frac{\sigma_{\text{PSF}}\sigma_{\text{bead}}}{\sigma_{\text{image}}} e^{-\frac{x^2}{2\sigma_{\text{image}}{}^2}}, \qquad \text{Eq. S4}$$

where $\sigma_{\text{image}}$ is the size of the image and can be expressed as

$$\sigma_{\text{image}} = \sqrt{\sigma_{\text{PSF}}^2 + \sigma_{\text{bead}}^2}. \qquad \text{Eq. S5}$$

As a result,

$$\sigma_{\text{PSF}} = \sqrt{\sigma_{\text{image}}^2 - \sigma_{\text{bead}}^2}. \qquad \text{Eq. S6}$$

For Gaussian functions, $\text{FWHM} = 2\sqrt{2\ln{(2)}}\sigma$. Substituting $\sigma$ with $\text{FWHM}$ in Eq. S6, we get

$$\text{FWHM}_{\text{PSF}} = \sqrt{\text{FWHM}_{\text{image}}^2 - \text{FWHM}_{\text{bead}}^2}. \qquad \text{Eq. S7}$$

**Supplementary Table 1 | Nanoscope component list.**

| Category | Part Number | Description | Vendor | Quantity |
|---|---|---|---|---|
| Optical table | 784-SPECIAL | 784-SPECIAL OPT.TOP/ 1,000 mm×1,500mm×SUPER 8 (nominally 200-mm thick) construction | TMC | 1 |
| Optical table | 14NP-417-89 | 14NP-417-89 SYSTEM 1/4000# CAP. 38×47×28H | TMC | 1 |
| Optical table | 14-OPTION | 14-OPTION OPTIONAL Retractable Casters in support stand above | TMC | 1 |
| Custom Parts | FAB-IS0001 | Goniometer Spacer | machine shop | 1 |
| Custom Parts | FAB-IS0002 | Goniometer Cube Holder | machine shop | 1 |
| Custom Parts | FAB-IS0003 | Filter Cube Emission | machine shop | 2 |
| Custom Parts | FAB-IS0005 | Upper Floating Plate | machine shop | 1 |
| Custom Parts | FAB-IS0006 | Jack Clamp Corner | machine shop | 2 |
| Custom Parts | FAB-IS0007 | Vertical Plate A - hole array | machine shop | 1 |
| Custom Parts | FAB-IS0008 | Lens Mount – 40 mm LOWER | machine shop | 2 |
| Custom Parts | FAB-IS0010 | BK7 Flat Mount | machine shop | 1 |
| Custom Parts | FAB-IS0011 | Shutter Mount | machine shop | 2 |
| Custom Parts | FAB-IS0012 | Cavity Half Inch Lens Holder LOWER | machine shop | 1 |
| Custom Parts | FAB-IS0013 | Cavity Half Inch Lens Holder UPPER | machine shop | 1 |
| Custom Parts | FAB-IS0014 | PRM1 Rotation Mount Holder | machine shop | 2 |
| Custom Parts | FAB-IS0015 | Stationary Wedge Holder | machine shop | 1 |
| Custom Parts | FAB-IS0016 | Table Fold Mirror | machine shop | 1 |
| Custom Parts | FAB-IS0017 | Galvo Mount | machine shop | 1 |
| Custom Parts | FAB-IS0018 | Mirror Base | machine shop | 1 |
| Custom Parts | FAB-IS0019p2 | Resonant Mirror Base | machine shop | 1 |
| Custom Parts | FAB-IS0019p3 | Resonant Mirror Mount Clamp | machine shop | 1 |
| Custom Parts | FAB-IS0020 | Resonant Mirror Mount Base | machine shop | 1 |
| Custom Parts | FAB-IS0021 | MTS25 Base Plate | machine shop | 1 |
| Custom Parts | FAB-IS0022 | Wedge Stage Connector | machine shop | 1 |
| Custom Parts | FAB-IS0023 | Mirror Cube Module LOWER | machine shop | 1 |
| Custom Parts | FAB-IS0024 | Mirror Cube Module UPPER | machine shop | 1 |
| Custom Parts | FAB-IS0025 | Piezo Mirror Holder | machine shop | 1 |
| Custom Parts | FAB-IS0026 | P841 Holder | machine shop | 1 |
| Custom Parts | FAB-IS0027 | Cube Holder Bracket (LOWER) | machine shop | 1 |
| Custom Parts | FAB-IS0028 | Quarter WP Rotation Mount | machine shop | 2 |

| | | | | |
|---|---|---|---|---|
| Custom Parts | FAB-IS0029 | Cavity Lens Translation Holder | machine shop | 2 |
| Custom Parts | FAB-IS0030 | Lens Trans Insert | machine shop | 2 |
| Custom Parts | FAB-IS0031 | Cube Holder Bracket UPPER | machine shop | 1 |
| Custom Parts | FAB-IS0032 | Custom Base Plate - hole array | machine shop | 1 |
| Custom Parts | FAB-IS0034 | SS100 Slot Mount | machine shop | 1 |
| Custom Parts | FAB-IS0037 | Vertical Support Cavity Leg Tall | machine shop | 1 |
| Custom Parts | FAB-IS0038 | Bridge Leg Connector | machine shop | 2 |
| Custom Parts | FAB-IS0040 | Thorlabs 66 Rail Holder STXY | machine shop | 3 |
| Custom Parts | FAB-IS0042 | SLM Assembly Plate - hole array | machine shop | 1 |
| Custom Parts | FAB-IS0043 | LM1XY Mount on SLM Plate | machine shop | 1 |
| Custom Parts | FAB-IS0044 | Right Angle Rail Mirror | machine shop | 1 |
| Custom Parts | FAB-IS0045 | Rail Lens Holder | machine shop | 1 |
| Custom Parts | FAB-IS0046 | OptoSigma Mount LP740RDC | machine shop | 1 |
| Custom Parts | FAB-IS0047 | OptoSigma Mount Custom RDC | machine shop | 1 |
| Custom Parts | FAB-IS0048 | Vertical Reference Plate | machine shop | 1 |
| Custom Parts | FAB-IS0049 | SLM Focus Rail | machine shop | 1 |
| Custom Parts | FAB-IS0050 | D Shaped Mirror Mount | machine shop | 1 |
| Custom Parts | FAB-IS0052 | Main Rail Carrier | machine shop | 1 |
| Custom Parts | FAB-IS0051 | Vertical Board Support Leg | machine shop | 2 |
| Custom Parts | FAB-IS0053 | Secondary Rail Carrier | machine shop | 2 |
| Custom Parts | FAB-IS0054 | Alignment Camera Connector | machine shop | 2 |
| Custom Parts | FAB-IS0057 | Camera Spacer | machine shop | 5 |
| Custom Parts | FAB-IS0058 | PMT Mount | machine shop | 1 |
| Custom Parts | FAB-IS0059 | Cavity Diag. Mirror Holder LOWER | machine shop | 1 |
| Custom Parts | FAB-IS0060 | Cavity Mirror Diag. Mirror Holder UPPER | machine shop | 1 |
| Custom Parts | FAB-IS0061 | Fixed Lens Trans Mount | machine shop | 1 |
| Custom Parts | FAB-IS0062 | Backside Mount | machine shop | 2 |
| Custom Parts | FAB-IS0063 | Pusher Mount with Cord Slot | machine shop | 1 |
| Custom Parts | FAB-IS0064 | Pusher Passive Cord | machine shop | 1 |
| Custom Parts | FAB-IS0065 | SLM Rotation Base Plate | machine shop | 1 |
| Custom Parts | FAB-IS0066 | SLM Rotation Mount | machine shop | 1 |
| Custom Parts | FAB-P0001 | Vertical Support Base | machine shop | 5 |
| Custom Parts | FAB-P0002 | Vertical Support Cavity Leg (Version 2) | machine shop | 1 |
| Custom Parts | FAB-P0003 | Support Bracket (with cut out center) | machine shop | 2 |

| | | | | |
|---|---|---|---|---|
| Custom Parts | FAB-P0004 | Lower Plate | machine shop | 1 |
| Custom Parts | FAB-P0005 | Upper Plate | machine shop | 1 |
| Custom Parts | FAB-P0006 | Foot Support | machine shop | 1 |
| Custom Parts | FAB-P0007 | Upper Objective Holder | machine shop | 1 |
| Custom Parts | FAB-P0008 | Jack Screw Clamp | machine shop | 1 |
| Custom Parts | FAB-P0009 | Sample Holder (updated cut outs) | machine shop | 1 |
| Custom Parts | FAB-P0011 | Piezo Pusher Base | machine shop | 2 |
| Custom Parts | FAB-P0012 | Piezo Pusher Plow | machine shop | 2 |
| Custom Parts | FAB-P0013 | Piezo Pusher Base, Passive | machine shop | 1 |
| Custom Parts | FAB-P0014 | Piezo Pusher Plow, Passive | machine shop | 1 |
| Custom Parts | FAB-P0015 | Lower Objective Holder | machine shop | 1 |
| Custom Parts | FAB-P0019 | Mirror Mount - SS100 | machine shop | 2 |
| Custom Parts | DM-Mock_01 | DM Main Body | machine shop | 2 |
| Custom Parts | AOM adapter | AOM adapter | machine shop | 1 |
| Custom Parts | AOTF adapter | AOTF adapter | machine shop | 1 |
| Custom Parts | MPB mounting plate | MPB mounting plate | machine shop | 1 |
| Optomechanics | 04 RDI 232 | Cavity Bi-Stable Shutter | CVI Melles Griot | 2 |
| Optomechanics | AJS100-0.5H-NL | Precision 100 TPI Hex Adjustment Screw, 12.7 mm Travel | Newport | 2 |
| Optomechanics | 9314-K | Hex Adjustment Screw, Unbraked, 25.4 mm Travel, Ball Tip, 1/4-80 | Newport | 2 |
| Optomechanics | SS100-F2H | Suprema Mirror Mount, 1.0 inch, (2) 100-TPI Locking Hex-Broach Actuators | Newport | 21 |
| Optomechanics | LP-2A | *xyz* $\theta_x\theta_y$ Lens Positioner, 2.0 in. (50.8 mm) Diameter | Newport | 2 |
| Optomechanics | SC100-R2H-LH | Suprema Clear Edge Rear Load Mount, 1.0 inch., 2 100-TPI Locking Hex, LH | Newport | 1 |
| Optomechanics | M-GON40-U | Upper Goniometric Stage, 40×40×20 $mm^3$, ±5° Travel, Metric | Newport | 1 |
| Optomechanics | CONEX-TRA12CC | TRA12CC Actuator, Integrated with CONEX-CC Controller | Newport | 2 |
| Optomechanics | CONEX-PS | Power Supply, 24 VDC, CONEX Motion Controllers | Newport | 2 |
| Optomechanics | M-RS65 | Aperture Platform Rotation Stage, 65 mm, 10° Fine, Metric | Newport | 1 |
| Optomechanics | M-423 | High-Performance Low-Profile Ball Bearing Linear Stage, 25.4 mm, M6 | Newport | 1 |

| | | | | |
|---|---|---|---|---|
| Optomechanics | SM-25 | Vernier Micrometer, 25 mm Travel, 23 lb Load Capacity, 50.8 TPI | Newport | 1 |
| Optomechanics | 9315 | Ball Tipped Nudger, 12.7 mm Travel, 1/4-80 | Newport | 3 |
| Optomechanics | FPR1-C1A | Fiber Optic Positioner, $xyz\theta_z$, Connectorized, FPH-CA Series | Newport | 2 |
| Optomechanics | M-UMR5.16 | Double-Row Ball Bearing Linear Stage, 16 mm, 600-N Load, Metric | Newport | 2 |
| Optomechanics | BM11.16 | Standard Resolution Micrometer, 16 mm Travel, 9-lb Load Capacity | Newport | 2 |
| Optomechanics | FPH-CA3 | Fiber Chuck, SMA Connectorized Fiber | Newport | 1 |
| Optomechanics | UPG-1 | Gimbaled Three-Axis Optic Tilt Mount | Newport | 2 |
| Optomechanics | FPH-CA4 | Fiber Chuck, FC Connectorized Fiber | Newport | 1 |
| Optomechanics | SS050-F3N | Suprema Optical Mirror Mount, 0.5 inch, (3) 100-TPI Allen-Key Actuators | Newport | 2 |
| Optomechanics | 9071-M | Four-Axis Tilt Aligner, 3 mm, 8°, M4 and M6 | Newport | 2 |
| Optomechanics | BSHL-25.4-2 | Compact Gimballed Beamsplitter Mounts / Compact Gimballed Beamsplitter | OptoSigma | 12 |
| Optomechanics | PRM05/M | High-Precision Rotation Mount for Ø1/2" (12.5 mm) Optics, Metric | Thorlabs | 2 |
| Optomechanics | RLA300/M | Dovetail Optical Rail, 300 mm, Metric | Thorlabs | 1 |
| Optomechanics | RLA600/M | Dovetail Optical Rail, 600 mm, Metric | Thorlabs | 1 |
| Optomechanics | PRM1/M | High-Precision Rotation Mount for Ø1" (25.4 mm) Optics, Metric | Thorlabs | 4 |
| Optomechanics | MTS25/M-Z8 | 25-mm (0.98") Motorized Translation Stage, M4 and M3 Taps | Thorlabs | 1 |
| Optomechanics | KDC101 | K-Cube Brushed DC Servo Motor Controller | Thorlabs | 1 |
| Optomechanics | KAP102 | Adapter Plate for KCH Series Hubs and 120-mm Wide T-Cubes | Thorlabs | 1 |
| Optomechanics | KCH301 | USB Controller Hub and Power Supply for Three K-Cubes or T-Cubes | Thorlabs | 1 |
| Optomechanics | LMR1/M | Lens Mount for Ø1" Optics, One Retaining Ring Included, M4 Tap | Thorlabs | 1 |
| Optomechanics | ST1XY-A/M | XY Translator with 100 TPI Drives, Metric | Thorlabs | 5 |

| | | | | |
|---|---|---|---|---|
| Optomechanics | XT66N | Rail Platform Locator for 66-mm Rails | Thorlabs | 4 |
| Optomechanics | LM1XY/M | Translating Lens Mount for Ø1" Optics, 1 Retaining Ring Included, Metric | Thorlabs | 2 |
| Optomechanics | KM100DL | Left-Handed Kinematic Mount for Ø1" D-Shaped Mirrors | Thorlabs | 1 |
| Optomechanics | CRM1P/M | Precision Cage Rotation Mount with Micrometer Drive, Ø1" Optics, M4 Tap | Thorlabs | 2 |
| Optomechanics | SM1Z | Z-Translator for Cage System | Thorlabs | 2 |
| Optomechanics | XT66SP-350-YALE-SP | 66-mm Single Dovetail Rail, L = 800 mm; custom hole positions | Thorlabs | 1 |
| Optomechanics | XT66SP-800-YALE-SP | 66-mm Single Dovetail Rail, L = 350 mm; custom hole positions | Thorlabs | 1 |
| Optomechanics | XT66C1 | 20-mm Long Double Dovetail Clamp for 66-mm Rails | Thorlabs | 10 |
| Optomechanics | B4CRP/M | 30-mm Cage Cube Precision Kinematic Rotation Platform (Metric) | Thorlabs | 1 |
| Optomechanics | B6C | 30-mm Cage Cube Clamp | Thorlabs | 1 |
| Optomechanics | C6W | 30-mm Cage Cube, Ø6 mm Through Holes | Thorlabs | 1 |
| Optomechanics | CF038C/M-P5 | Clamping Fork, 9.5-mm Counterbored Slot, M6×1.0 Captive Screw, 5 Pack | Thorlabs | 3 |
| Optomechanics | CP02/M | SM1-Threaded 30 mm Cage Plate, 0.35" Thick, 2 Retaining Rings, M4 Tap | Thorlabs | 1 |
| Optomechanics | CRM1P/M | Precision Cage Rotation Mount with Micrometer Drive, Ø1" Optics, M4 Tap | Thorlabs | 1 |
| Optomechanics | ER1-P4 | Cage Assembly Rod, 1" Long, Ø6 mm | Thorlabs | 3 |
| Optomechanics | KC05-T/M | SM05 Threaded Kinematic Cage Mount, Ø1/2" Optics, Metric | Thorlabs | 2 |
| Optomechanics | KCB05/M | Right-Angle Kinematic Mirror Mount, 16 mm Cage System and SM05 Compatible, M3 and M4 Mounting Holes | Thorlabs | 1 |
| Optomechanics | KMCP/M | Kinematic Mount Centering Plate, Metric | Thorlabs | 5 |
| Optomechanics | LM1XY/M | Translating Lens Mount for Ø1" Optics, 1 Retaining Ring Included, Metric | Thorlabs | 5 |
| Optomechanics | PRM1/M | High-Precision Rotation Mount for Ø1" (25.4 mm) Optics, Metric | Thorlabs | 1 |
| Optomechanics | RC1 | Dovetail Rail Carrier, 1.00" x 1.00" (25.4×25.4 $mm^2$), 1/4" (M6) Counterbore | Thorlabs | 10 |

| | | | | |
|---|---|---|---|---|
| Optomechanics | RLA150/M | Dovetail Optical Rail, 150 mm, Metric | Thorlabs | 1 |
| Optomechanics | RLA300/M | Dovetail Optical Rail, 300 mm, Metric | Thorlabs | 4 |
| Optomechanics | RS05P4M | Ø25.0 mm Pedestal Pillar Post, M4 Taps, L = 12.5 mm | Thorlabs | 3 |
| Optomechanics | RS10M | Ø25 mm Post Spacer, Thickness = 10 mm | Thorlabs | 3 |
| Optomechanics | RS19/M | Ø25.0 mm Pillar Post, M6 Taps, L = 19 mm, M4 Adapter Included | Thorlabs | 1 |
| Optomechanics | RS1P4M | Ø25.0 mm Pedestal Pillar Post, M4 Taps, L = 25 mm | Thorlabs | 1 |
| Optomechanics | RS2.5P4M | Ø25.0 mm Pedestal Pillar Post, M4 Taps, L = 65 mm | Thorlabs | 4 |
| Optomechanics | RS25/M | Ø25.0-mm Pillar Post, M6 Taps, L = 25 mm, M4 Adapter Included | Thorlabs | 6 |
| Optomechanics | RS2P/M | Ø25.0-mm Pedestal Pillar Post, M6 Taps, L = 50 mm | Thorlabs | 2 |
| Optomechanics | RS2P4M | Ø25.0 mm Pedestal Pillar Post, M4 Taps, L = 50 mm | Thorlabs | 7 |
| Optomechanics | RS38/M | Ø25.0-mm Pillar Post, M6 Taps, L = 38 mm, M4 Adapter Included | Thorlabs | 8 |
| Optomechanics | RS3M | Ø25-mm Post Spacer, Thickness = 3 mm | Thorlabs | 4 |
| Optomechanics | RS3P4M | Ø25.0-mm Pedestal Pillar Post, M4 Taps, L = 75 mm | Thorlabs | 1 |
| Optomechanics | RS4M | Ø25-mm Post Spacer, Thickness = 4 mm | Thorlabs | 2 |
| Optomechanics | RS5M | Ø25-mm Post Spacer, Thickness = 5 mm | Thorlabs | 6 |
| Optomechanics | RS7/M | Ø25-mm Post, M6 Tap, L = 7 mm | Thorlabs | 1 |
| Optomechanics | SB6C | 16-mm Cube Clamp | Thorlabs | 1 |
| Optomechanics | SC6W | 16-mm Cage Cube | Thorlabs | 1 |
| Optomechanics | SM1A3 | Adapter with External SM1 Threads and Internal RMS Threads | Thorlabs | 1 |
| Optomechanics | SM1FC | FC/PC Fiber Adapter Plate with External SM1 (1.035"-40) Thread | Thorlabs | 3 |
| Optomechanics | SP03 | Compact Cage Plate with 16-mm Aperture for a 16-mm Cage System | Thorlabs | 1 |
| Optomechanics | SPM2 | 16-mm Cage Prism Mount | Thorlabs | 1 |
| Optomechanics | SR1-P4 | Compact Cage Assembly Rod, 1" Long, Ø4 mm | Thorlabs | 1 |
| Optomechanics | SR2-P4 | Compact Cage Assembly Rod, 2" Long, Ø4 mm | Thorlabs | 2 |
| Optomechanics | SRM05 | 16 mm Cage Rotation Mount for Ø1/2" Optics | Thorlabs | 3 |

| | | | | |
|---|---|---|---|---|
| Optomechanics | SRSCA-P4 | Rod Adapter for Ø4-mm SR Rods, 4 Pack | Thorlabs | 1 |
| Optomechanics | ST1XY-S/M | XY Translator with Micrometer Drives, Metric | Thorlabs | 3 |
| Optomechanics | RP005/M | Ø1" Manual Rotation Stage, Metric | Thorlabs | 1 |
| Optomechanics | RS4M | Ø25 mm Post Spacer, Thickness = 4 mm | Thorlabs | 1 |
| Optomechanics | FM90/M | Flip Mount Adapter, Metric | Thorlabs | 1 |
| Optomechanics | KMCP/M | Kinematic Mount Centering Plate, Metric | Thorlabs | 1 |
| Optomechanics | KMS/M | Compact Kinematic Mirror Mount, M4 Taps for Post Mounting | Thorlabs | 1 |
| Optomechanics | MH25 | Mirror Holder for Ø1" Optics 2.5 - 6.1-mm Thick | Thorlabs | 1 |
| Optomechanics | CXY1 | 30 mm Cage System, XY Translating Lens Mount for Ø1" Optics | Thorlabs | 2 |
| Optomechanics | CP02/M | SM1-Threaded 30 mm Cage Plate, 0.35" Thick, 2 Retaining Rings, M4 Tap | Thorlabs | 1 |
| Optomechanics | SM1L20 | SM1 Lens Tube, 2.00" Thread Depth, One Retaining Ring Included | Thorlabs | 1 |
| Optomechanics | BD40 | Calcite Beam Displacer, 4.0 mm Beam Separation, Ø1" Housing | Thorlabs | 1 |
| Optomechanics | SM1T1 | SM1 (1.035"-40) Coupler, External Threads, 0.5" Long | Thorlabs | 1 |
| Optomechanics | SM1T2 | SM1 (1.035"-40) Coupler, External Threads, 0.5" Long | Thorlabs | 1 |
| Optomechanics | RS1M | Ø25 mm Post Spacer, Thickness = 1 mm | Thorlabs | 1 |
| Optomechanics | RS4M | Ø25 mm Post Spacer, Thickness = 4 mm | Thorlabs | 2 |
| Optomechanics | RS2M | Ø25 mm Post Spacer, Thickness = 2 mm | Thorlabs | 1 |
| Optomechanics | RS7M | Ø25 mm Post Spacer, Thickness = 7 mm | Thorlabs | 2 |
| Optomechanics | RC1 | Dovetail Rail Carrier, 1.00"×1.00" (25.4×25.4 $mm^2$), 1/4" (M6) Counterbore | Thorlabs | 6 |
| Optomechanics | RS38/M | 25.0 mm Pillar Post, M6 Taps, L = 38 mm, M4 Adapter Included | Thorlabs | 3 |
| Optomechanics | RS25/M | 25.0 mm Pillar Post, M6 Taps, L = 25 mm, M4 Adapter Included | Thorlabs | 1 |
| Optomechanics | ER4-P4 | Cage Assembly Rod, 4" Long, Ø6 mm, 4 Pack | Thorlabs | 1 |
| Optomechanics | MS1R/M | Mini-Series Optical Post, Ø6 mm, L = 25 mm | Thorlabs | 8 |

| | | | | |
|---|---|---|---|---|
| Optomechanics | MS1.5R/M | Mini-Series Optical Post, Ø6 mm, L = 38 mm | Thorlabs | 8 |
| Optomechanics | AE3M6M | Adapter with Internal M3×0.5 Threads and External M6×1.0 Threads | Thorlabs | 10 |
| Optomechanics | CP20S | 30 mm Cage System Iris, Ø20.0-mm Maximum Aperture | Thorlabs | 2 |
| Optomechanics | ER2-P4 | Cage Assembly Rod, 2" Long, Ø6 mm, 4 Pack | Thorlabs | 1 |
| Optomechanics | SP05/M | 30-mm to 16-mm Cage Adapter Plate, Metric | Thorlabs | 1 |
| Optomechanics | RS05P/M | Ø25.0-mm Pedestal Pillar Post, M6 Taps, L = 12.5 mm | Thorlabs | |
| Optomechanics | RS10M | Ø25-mm Post Spacer, Thickness = 10 mm | Thorlabs | 1 |
| Optomechanics | ST1XY-S/M | *xy* Translator with Micrometer Drives, Metric | Thorlabs | 3 |
| Optomechanics | RC4 | Dovetail Rail Carrier, 0.60"×1.00" (15.2×25.4 $mm^2$), #8 (M4) Counterbore | Thorlabs | 2 |
| Optomechanics | ST1XY-A/M | *xy* Translator with 100 TPI Drives | Thorlabs | 1 |
| Optomechanics | SM1Z | Z-Axis Translation Mount, 30-mm Cage Compatible | Thorlabs | 1 |
| Optomechanics | ER2-P4 | Cage Assembly Rod, 2" Long, Ø6 mm, 4 Pack | Thorlabs | 1 |
| Optomechanics | LM1XY/M | Translating Lens Mount for Ø1" Optics, 1 Retaining Ring Included, Metric | Thorlabs | 2 |
| Optomechanics | SM1A6 | Adapter with External SM1 Threads and Internal SM05 Threads, 0.15" Thick | Thorlabs | 1 |
| Optomechanics | RLA150/M | Dovetail Optical Rail, 150 mm, Metric | Thorlabs | 2 |
| Optomechanics | RLA300/M | Dovetail Optical Rail, 300 mm, Metric | Thorlabs | 1 |
| Optomechanics | RC1 | Dovetail Rail Carrier, 1.00"×1.00" (25.4×25.4 $mm^2$), 1/4" (M6) Counterbore | Thorlabs | 5 |
| Optomechanics | RC3 | Rail Carrier, Perpendicular Dovetail | Thorlabs | 5 |
| Optomechanics | RS25/M | Ø25.0-mm Pillar Post, M6 Taps, L = 25 mm, M4 Adapter Included | Thorlabs | 5 |
| Optomechanics | RS3M | Ø25-mm Post Spacer, Thickness = 3 mm | Thorlabs | 2 |
| Optomechanics | RS10M | Ø25-mm Post Spacer, Thickness = 10 mm | Thorlabs | 2 |
| Optomechanics | RS2M | Ø25-mm Post Spacer, Thickness = 2 mm | Thorlabs | 1 |
| Optomechanics | RS38/M | Ø25.0-mm Pillar Post, M6 Taps, L = 38 mm, M4 Adapter Included | Thorlabs | 3 |
| Optomechanics | RS2M | Ø25-mm Post Spacer, Thickness = 2 mm | Thorlabs | 1 |

| | | | | |
|---|---|---|---|---|
| Optomechanics | RS4P/M | Ø25.0-mm Pedestal Pillar Post, M6 Taps, L = 100 mm | Thorlabs | 2 |
| Optomechanics | RS8M | Ø25-mm Post Spacer, Thickness = 8 mm | Thorlabs | 2 |
| Optomechanics | RS05P/M | Ø25.0-mm Pedestal Pillar Post, M6 Taps, L = 12.5 mm | Thorlabs | 3 |
| Optomechanics | SM05L03 | SM05 Lens Tube, 0.30" Thread Depth, One Retaining Ring Included | Thorlabs | 2 |
| Optomechanics | CF038C/M-P5 | Clamping Fork, 9.5-mm Counterbored Slot, M6×1.0 Captive Screw, 5 Pack | Thorlabs | 1 |
| Optomechanics | ER1-P4 | Cage Assembly Rod, 1" Long, Ø6 mm, 4 Pack | Thorlabs | 1 |
| Optomechanics | RS1LM | Ø25-mm Laminated Post Spacer, 1-mm Thick, Pack of 5 | Thorlabs | 2 |
| Optomechanics | RS9M | Ø25 mm Post Spacer, Thickness = 9 mm | Thorlabs | 2 |
| Optomechanics | RM1G | 1" Construction Cube, Three 1/4" (M6) Counterbored Holes | Thorlabs | 1 |
| Optomechanics | SM1D12 | SM1 Lever-Actuated Iris Diaphragm (Ø0.8 - Ø12 mm) | Thorlabs | 2 |
| Optomechanics | TR50/M | Ø12.7-mm Optical Post, SS, M4 Setscrew, M6 Tap, L = 50 mm | Thorlabs | 1 |
| Optomechanics | SM1A3 | Adapter with External SM1 Threads and Internal RMS Threads | Thorlabs | 1 |
| Optomechanics | RLA150/M | Dovetail Optical Rail, 150 mm, Metric | Thorlabs | 1 |
| Optomechanics | RC1 | Dovetail Rail Carrier, 1.00"×1.00" (25.4×25.4 $mm^2$), 1/4" (M6) Counterbore | Thorlabs | 3 |
| Optomechanics | RS25/M | Ø25.0-mm Pillar Post, M6 Taps, L = 25 mm, M4 Adapter Included | Thorlabs | 3 |
| Optomechanics | LM1XY/M | Translating Lens Mount for Ø1" Optics, 1 Retaining Ring Included, Metric | Thorlabs | 2 |
| Optomechanics | SM1A6 | Adapter with External SM1 Threads and Internal SM05 Threads, 0.15" Thick | Thorlabs | 1 |
| Optomechanics | CF125C/M | Clamping Fork, 44.8-mm Counterbored Slot, M6×1.0 Captive Screw | Thorlabs | 1 |
| Optomechanics | RC3 | Rail Carrier, Perpendicular Dovetail | Thorlabs | 2 |
| Optomechanics | CRM1P/M | Precision Cage Rotation Mount with Micrometer Drive, Ø1" Optics, M4 Tap | Thorlabs | 1 |
| Optomechanics | SM1D12 | SM1 Lever-Actuated Iris Diaphragm (Ø0.8 - Ø12 mm) | Thorlabs | 1 |

| | | | | |
|---|---|---|---|---|
| Optomechanics | RC3 | Rail Carrier, Perpendicular Dovetail | Thorlabs | 1 |
| Optomechanics | CRM1P/M | Precision Cage Rotation Mount with Micrometer Drive, Ø1" Optics, M4 Tap | Thorlabs | 1 |
| Optomechanics | ER1-P4 | Cage Assembly Rod, 1" Long, Ø6 mm, 4 Pack | Thorlabs | 1 |
| Optomechanics | SM1L30 | SM1 Lens Tube, 3.00" Thread Depth, One Retaining Ring Included | Thorlabs | 2 |
| Optomechanics | SM1V10 | Ø1" Adjustable Lens Tube, 0.81" Travel Range | Thorlabs | 3 |
| Optomechanics | SM1L40 | SM1 Lens Tube, 4.00" Thread Depth, One Retaining Ring Included | Thorlabs | 1 |
| Optomechanics | RC3 | Rail Carrier, Perpendicular Dovetail | Thorlabs | 3 |
| Optomechanics | CRM1P/M | Precision Cage Rotation Mount with Micrometer Drive, Ø1" Optics, M4 Tap | Thorlabs | 1 |
| Optomechanics | RC1 | Dovetail Rail Carrier, 1.00"×1.00" (25.4×25.4 $mm^2$), 1/4" (M6) Counterbore | Thorlabs | 2 |
| Optomechanics | H45 | 45° Mirror Mount for Ø1" Optics | Thorlabs | 2 |
| Optomechanics | FMP1/M | Fixed Ø1" Mirror Mount, M4 Tap | Thorlabs | 2 |
| Optomechanics | RS3M | Ø25-mm Post Spacer, Thickness = 3 mm | Thorlabs | 2 |
| Optomechanics | RS38/M | Ø25.0-mm Pillar Post, M6 Taps, L = 38 mm, M4 Adapter Included | Thorlabs | 2 |
| Optics | ZT485/595/640/775-rpc-UF5 | Laser/STED polychroic mirror, R: 485, 595, 640 and 775 nm, T: 500-550, 600-635 and 665-740 nm, 25-mm diameter, 5-mm thick substrate, 1/4 wave RWF post-coat | Chroma | 2 |
| Optics | ZT1064rdc-sp | Reflects lasers between 950-1150 nm, 1-mm thick, 18×26 $mm^2$ | Chroma | 2 |
| Optics | T740lpxr | 25.4-mm Diameter, 5-mm Thick $\lambda$/8 RWD | Chroma | 1 |
| Optics | ZT568rdc | 25.4-mm Diameter, 3-mm Thick $\lambda$/8 RWD | Chroma | 1 |
| Optics | ZT640rdc | 25.4-mm Diameter, 3-mm Thick $\lambda$/8 RWD | Chroma | 1 |
| Optics | 49-331 | 12.5-mm Diameter 80-mm FL, VIS-NIR Coated, Achromatic Lens | Edmund Optics | 2 |
| Optics | 49-384 | 40-mm Diameter×250-mm FL, VIS-NIR Coated, Achromatic Lens | Edmund Optics | 1 |

| | | | | |
|---|---|---|---|---|
| Optics | 49-359 | 25-mm Diameter×85-mm FL, VIS-NIR Coated, Achromatic Lens | Edmund Optics | 1 |
| Optics | 48-777 | 25.0-mm Diameter×175.0-mm FL, NIR I Coated, Plano-Convex Lens | Edmund Optics | 3 |
| Optics | 49-354 | 25-mm Diameter×40-mm FL, VIS-NIR Coated, Achromatic Lens | Edmund Optics | 1 |
| Optics | 49-362 | 25-mm Diameter×150-mm FL, VIS-NIR Coated, Achromatic Lens | Edmund Optics | 1 |
| Optics | 49-381 | 40-mm Diameter×120-mm FL, VIS-NIR Coated, Achromatic Lens | Edmund Optics | 1 |
| Optics | 49-872 | 25-mm NIR, Polarizing Cube Beamsplitter | Edmund Optics | 2 |
| Optics | 49-228 | ½ wave 630-835 nm, Precise Polymer Achromatic Retarder | Edmund Optics | 2 |
| Optics | 49-361 | 25-mm Diameter×125-mm FL, VIS-NIR Coated, Achromatic Lens | Edmund Optics | 2 |
| Optics | DBL14017100 | Convex Lens, 90-mm focal length | JML | 2 |
| Optics | BPB-25.4SF2-550 | Broadband Polarizing Cube Beamsplitter | Lambda | 1 |
| Optics | 10Z40ER.2 | Broadband Metallic Mirror, Zerodur, 25.4 mm, $\lambda$/20, 480-20,000 nm | Newport | 26 |
| Optics | F-LA11 | Laser Diode Aspheric Objective Lens, 0.40 NA, 6.24-mm FL, 510-1550 nm | Newport | 1 |
| Optics | 05D620BD.2 | ValuMax Broadband Dielectric Mirrors | Newport | 1 |
| Optics | M-20X | 20×, 0.40 NA, 9.0-mm Focal Length | Newport | 1 |
| Optics | 1-U2B714 * | UPLSAPO100XS; UPLSAPO 100X SI OIL OBJ, NA 1.35,WD 0.20MM,W/CC | Olympus | 2 |
| Optics | BLP01-594R-25 | 594-nm EdgeBasic™ best-value long-pass edge filter | Semrock/AVR optics | 1 |
| Optics | BLP01-647R-25 | 647-nm EdgeBasic™ best-value long-pass edge filter | Semrock/AVR optics | 1 |
| Optics | FF02-685/40-25 | 685/40 BrightLine® Bandpass Filter, 25 mm | Semrock/AVR optics | 2 |
| Optics | FF01-624/40-25 | 624/40 BrightLine® Bandpass Filter, 25 mm | Semrock/AVR optics | 2 |
| Optics | FF03-525/50-25 | 525/50 BrightLine® Bandpass Filter, 25 mm | Semrock/AVR optics | 2 |
| Optics | AQWP05M-600 | Quarter wave plates; 1/2" unmounted zero order | Thorlabs | 2 |
| Optics | AHWP10M-600 | SM1-Mounted Achromatic Half-Wave Plate, Ø22.6 mm, 400 - 800 nm | Thorlabs | 2 |

| | | | | |
|---|---|---|---|---|
| Optics | AC254-075-B | f=75.0 mm, Ø1" Achromatic Doublet, ARC: 400-700 nm | Thorlabs | 1 |
| Optics | PBS10-780 | 10 mm Polarizing Beamsplitter Cube, 780 nm | Thorlabs | 1 |
| Optics | BB0511-E03 | Ø1/2" (Ø12.7 mm) Zerodur® Broadband Dielectric Mirror, 750-1100 nm | Thorlabs | 5 |
| Optics | WPQ05M-780 | Ø1/2" Mounted Zero-Order Quarter-Wave Plate, Ø1" Mount, 780 nm | Thorlabs | 2 |
| Optics | WPH05M-780 | Ø1/2" Mounted Zero-Order Half-Wave Plate, Ø1" Mount, 780 nm | Thorlabs | 1 |
| Optics | PBS251 | 1" Polarizing Beamsplitter Cube, 420 – 680 nm | Thorlabs | 1 |
| Optics | AQWP05M-600 | Ø1/2" Mounted Achromatic Quarter-Wave Plate, Ø1" Mount, 400 - 800 nm | Thorlabs | 1 |
| Optics | AHWP10M-600 | Ø1" Mounted Achromatic Half-Wave Plate, SM1-Threaded Mount, 400 - 800 nm | Thorlabs | 1 |
| Optics | BB1-E02 | Ø1" Broadband Dielectric Mirror, 400 - 750 nm | Thorlabs | 4 |
| Optics | AQWP10M-980 | SM1-Mounted Achromatic Quarter-Wave Plate, Ø22.6 mm, 690 - 1200 nm | Thorlabs | 1 |
| Optics | PFD10-03-P01 | 1" Protected Silver D-Shaped Mirror | Thorlabs | 1 |
| Optics | PFE05-P01 | 1/2" Protected Silver Elliptical Mirror, 450 nm - 20 µm | Thorlabs | 2 |
| Optics | AC254-030-B | f=30.0 mm, Ø1" Achromatic Doublet, ARC: 650 - 1050 nm | Thorlabs | 1 |
| Optics | AC254-060-A | f=60.0 mm, Ø1" Achromatic Doublet, ARC: 400 - 700 nm | Thorlabs | 1 |
| Optics | PS879-A | N-KZFS8 Mounted Prism Pair, ARC: 350 - 700 nm, Mag: 3.0 | Thorlabs | 3 |
| Optics | KM200V | Large Kinematic V-Clamp Mount | Thorlabs | 3 |
| Optics | A397TM-A | f = 11.00 mm, NA = 0.30, Mounted Rochester Aspheric Lens, AR: 350 - 700 nm | Thorlabs | 1 |
| Optics | AC254-150-A | f = 150.0 mm, Ø1" Achromatic Doublet, ARC: 400 - 700 nm | Thorlabs | 1 |
| Optics | NE10A-B | Ø25 mm AR-Coated Absorptive Neutral Density Filter, 650 - 1050 nm, SM1-Threaded Mount, OD:1.0 | Thorlabs | 1 |
| Optics | NE20A-B | Ø25-mm AR-Coated Absorptive Neutral Density Filter, 650 - 1050 nm, SM1-Threaded Mount, OD: 2.0 | Thorlabs | 1 |
| Optics | AC254-060-B | f = 60.0 mm, Ø1" Achromatic Doublet, ARC: 650 - 1050 nm | Thorlabs | 1 |

| | | | | |
|---|---|---|---|---|
| Optics | FG105LCA + FT05SS | 0.22 NA, 400 - 2400 nm, 105-µm core, Ø5.0-mm stainless steel tubing, 2-m long | Thorlabs | 3 |
| Optics | --- | BK-7 Wedge and Plate | UVISIR | 1 |
| Optics | AC127-025-B-ML | f = 25 mm, Ø1/2" Achromatic Doublet, SM05-Threaded Mount, ARC: 400 - 700 nm | Thorlabs | 1 |
| Optics | AC254-400-B | f = 400.0 mm, Ø1" Achromatic Doublet, ARC: 650 - 1050 nm | Thorlabs | 1 |
| Optics | LJ1144RM-B | f = 500 mm, Ø1", N-BK7 Mounted Plano-Convex Round Cyl Lens, ARC: 650 - 1050 nm | Thorlabs | 1 |
| Optics | 20Z40ER.2 | Zerodur Broadband Metallic Mirror, 50.8 mm, $\lambda$/20, 480 - 20,000 nm | Newport | 1 |
| Optics | BLP01-808R-25 | 808 nm EdgeBasic™ best-value long-pass edge filter | Semrock/AVR optics | 1 |
| Optics | ND40A | Reflective Ø25-mm ND Filter, SM1-Threaded Mount, Optical Density: 4.0 | Thorlabs | 4 |
| Optics | NE10A | Ø25-mm Absorptive ND Filter, SM1-Threaded Mount, Optical Density: 1.0 | Thorlabs | 4 |
| Optics | LL01-488-25 | 488 nm MaxLine® laser clean-up filter | Semrock/AVR optics | 1 |
| Optics | FF01-591/6-25 | 591/6 nm BrightLine® single-band bandpass filter | Semrock/AVR optics | 1 |
| Optics | LL01-647-25 | 647.1 nm MaxLine® laser clean-up filter | Semrock/AVR optics | 1 |
| Optics | ET575lp | 25-mm diameter, 575 nm long-pass filter | Chroma | 1 |
| Optics | ET750sp-2p8 | 25-mm diameter, 750 nm long-pass filter | Chroma | 1 |
| Optics | AC127-025-A-ML | f = 25 mm, Ø1/2" Achromatic Doublet, SM05-Threaded Mount, ARC: 400 - 700 nm | Thorlabs | 1 |
| Optics | AQWP05M-600 | Ø1/2" Mounted Achromatic Quarter-Wave Plate, Ø1" Mount, 400 - 800 nm | Thorlabs | 1 |
| Optics | BB0511-E02 | Ø1/2" (Ø12.7 mm) Zerodur® Broadband Dielectric Mirror, 400 - 750 nm | Thorlabs | 1 |
| Optics | PBSH-450-1300-100 | Visible and Near-IR Broadband Polarizing Cube Polarizers, 450-1300nm, 25.4×25.4×25.4, Broadband Dielectric | CVI laser optics | 1 |
| Optics | ZET442/514/561 | 25-mm diameter, multi-band-pass | Chroma | 1 |
| Optics | AHWP05M-600 | Ø1/2" Mounted Achromatic Half-Wave Plate, Ø1" Mount, 400 - 800 nm | Thorlabs | 1 |
| Optics | ET655lp | 25-mm diameter, 655 nm long-pass filter | Chroma | 1 |

| | | | | |
|---|---|---|---|---|
| Optics | AT600lp | 25-mm diameter, 600 nm long-pass filter | Chroma | 1 |
| Optics | RET493lp | 25-mm diameter, 493 nm long-pass filter | Chroma | 1 |
| Actuators | MS1+1 | Two Axis DC Servo Motor controller w/ single control knob Control electronics included ASI's proven antibacklash algorithm to increase bi-directional accuracy. | ASI | 2 |
| Actuators | LS-50-AMCLL | Two LS-50 Linear Stages, with sub-micron accuracy, and 50 mm of travel Closed loop positioning utilizing rotary encoders. Standard 1.59-mm pitch lead screws provide 5.5-nm encoder resolution and < 20 nm intrinsic motor movement resolution. | ASI | 4 |
| Actuators | LS-LS-ME | LS-50 Micro-E Linear Encoders | ASI | 4 |
| Actuators | U-751.24 | Precision linear stage with travel range of 25×25 $mm^2$, maximum velocity of 100 mm/s, minimum incremental motion of 0.3 μm, load capacity of 50 N, repeatability of 0.3 μm, pitch of 50 μrad. PILine® ultrasonic piezomotor, performance class 2 with integrated Incremental, Optical, Direct Measuring feedback. | PI | 1 |
| Actuators | C-867.2U2 | PILine(R) Controller, 2 channels, highly precise, DSUB connector | PI | 1 |
| Actuators | M-227.10 | High-Resolution Linear Actuator with Closed-Loop DC Servo Motor, 10 mm | PI | 3 |
| Actuators | M-219.10 | Spherical End Piece, M10×0.5 mm | PI | 3 |
| Actuators | C-863.11 | Mercury DC Motor Controller, 1 Channel, with Wide-Range Power Supply | PI | 3 |
| Actuators | P-541.ZCD | Vertical Nanopositioning Stage with Large Aperture, 100 μm, Direct Metrology, Capacitive Sensors | PI | 1 |

| | | | | |
|---|---|---|---|---|
| Actuators | E-621.CR | Piezo Amplifier / Servo Controller Module, 1 Channel, -30 to 130 V, Capacitive Sensor, USB, RS-232 | PI | 1 |
| Actuators | P-612.2SL | *xy* Piezo Flexure Nanopositioning System with 20×20-mm$^2$ Aperture, 100×100 μm, Closed-Loop, Strain Gauge Sensors | PI | 1 |
| Actuators | E-621.SR | Piezo Amplifier / Servo Controller Module, 1 Channel, -30 to 130V, Closed-Loop, SGS Sensor, USB, RS-232 | PI | 2 |
| Actuators | E-501.621 | 9.5''-Chassis for up to four E-621 Modules, Power Supply | PI | 1 |
| Actuators | N-565.360 | Precision linear stage with travel range of 52 mm, maximum velocity of 10 mm/s, minimum incremental motion of 0.002 μm, load capacity of 20 N, repeatability of 0.005 μm, pitch of 80 μrad, and flatness of 1.5 μm. NEXACT® piezo stepping drive with integrated Pione Linear Encoder feedback. | PI | 1 |
| Actuators | E-861.1A1 | NEXACT® Controller, 1 Channel, Linear Encoder | PI | 1 |
| Actuators | P-841.10 | Preloaded piezo actuator, 15-μm travel range, 1000 N / 50 N, strain gauge sensor | PI | 1 |
| Actuators | E-709.SRG | Digital piezo controller, 1 axis, -30 to 130 V, strain gauge sensor, benchtop device | PI | 1 |
| Light sources | S422-205-000 | SuperK EXTREME EXR-20 | NKT Photonics | 1 |
| Light sources | TLD001 | T-Cube Laser Diode Controller (Power Supply Not Included) | Thorlabs | 1 |
| Light sources | LP980-SF15 | 980 nm, 15 mW, E Pin Code, SM Fiber-Pigtailed Laser Diode, FC/PC | Thorlabs | 1 |
| Light sources | SR9A-DB9 | ESD Protection and Strain Relief Cables | ThorLabs | 1 |
| Light sources | PTLB1 | Ø1/2" Post-Mountable Bracket for Fiber-Pigtailed Laser Diodes | ThorLabs | 1 |
| Light sources | PFL-80-3000-775-B1R | 775 nm, 3 W laser | MPB Communications | 1 |
| Miscellaneous Parts | 90974A111 | Black-Oxide Steel Partial-Thread T-Slot Nut | McMaster | 1 |

| | | | | |
|---|---|---|---|---|
| Miscellaneous Parts | SPW603 | SM05 Spanner Wrench, Length = 1" | Thorlabs | 1 |
| Miscellaneous Parts | 93070A143 | Low-Profile Alloy Steel Socket Head Cap Screws, 4-mm head, 12-mm long | McMaster | 1 |
| Miscellaneous Parts | 93070A098 | Low-Profile Alloy Steel Socket Head Cap Screws, 2.8-mm head, 8-mm long | Mcmaster | 1 |
| Miscellaneous Parts | 95345A020 | Pan Head Phillips Machine Screws with Lock Washer, 2-56, 3/4" length | McMaster | 1 |
| Miscellaneous Parts | CBSS4-10 | Ultra Low profile M4 Screw | MiSUMi | 1 |
| Miscellaneous Parts | 66103-4 | Contact; Pin; 16; Brass; Signal; Matte tin over Nickel; Plating Gold (30); 24-20AWG; Cri | Allied electronics | 20 |
| Miscellaneous Parts | 770147-1 | .093 Soft-Shell Pin & Socket; Contacts(Pre-Tinned Brass); Pin; for 24-18 AWG | Allied electronics | 20 |
| Miscellaneous Parts | 770078-1 | Connector; Soft Shell; 250 VAC; 13 A (Max.); Plug; Nylon; Natural; 4; 0.093 in. | Allied electronics | 5 |
| Miscellaneous Parts | 770075-1 | Connector; Shell; 250 VAC; 13 A (Max.); 4; Receptacle; Nylon; Natural; In-Line | Allied electronics | 5 |
| Miscellaneous Parts | 770146-1 | .093 Soft-Shell Pin & Socket; Contacts(Pre-Tinned Brass); Socket; for 24-18AWG | Allied electronics | 20 |
| Miscellaneous Parts | PRC03-32A10-7M10.5 PRC03 | PRC03 Relay Adapter (One-touch Lock) | MiSUMi | 1 |
| Miscellaneous Parts | 93285A21 | 18-8 Stainless Steel Nylon-Tip Set Screw, M4×0.7-mm Thread, 5-mm Long, 5 pack | McMaster | 1 |
| Miscellaneous Parts | 93285A014 | 18-8 Stainless Steel Nylon-Tip Set Screw, M2×0.4-mm Thread, 4 mm Long | McMaster | 4 |
| Miscellaneous Parts | 28130-67-01 | Cord; Power; 5-15P Plug; Wire Leads; 3 Cond SJT Cbl; 6'7"; 18 AWG; Black | Allied electronics | 2 |
| Miscellaneous Parts | HW-KIT3/M | M4 Setscrew and Hardware Kit | Thorlabs | 1 |
| Miscellaneous Parts | TC3/M | 15-Piece Balldriver and Hex Key Kit with Stand, Metric | Thorlabs | 1 |
| Miscellaneous Parts | CBSS6-12 | Ultra Low profile M6 Screw | MiSUMi | 1 |
| Miscellaneous Parts | CT4098-100 | RF Cable Assemblies 40 IN BNC M TO M RG223/U CBL ASSMBLY | Mouser | 7 |
| Miscellaneous Parts | CT4098-200 | RF Cable Assemblies 80 IN BNC M TO M RG223/U CBL ASSMBLY | Mouser | 4 |
| Miscellaneous Parts | CMS002 | CMS002 50×50 $mm^2$ Black Cable Trunking, 1-m (3.2 ft) Long | ThorLabs | 1 |

| | | | | |
|---|---|---|---|---|
| Miscellaneous Parts | CCL-1 | Cable Clamps, Qty 5 Short and 5 Long | Newport | 3 |
| Miscellaneous Parts | 7432K69 | Spiral Bundling Wrap, UV-Resistant Polyethylene, 1/16" ID, 1/8" OD 25ft | McMaster Carr | 1 |
| Miscellaneous Parts | 7432K63 | Spiral Bundling Wrap, UV-Resistant Polyethylene, 1/4" ID, 3/8" OD 25ft | McMaster Carr | 1 |
| Miscellaneous Parts | 34076417 | StarTech.com Rack Shelf 1U x 10"d | Gov Connection | 1 |
| Miscellaneous Parts | 274987 | StarTech.com Rack Shelf 2U x 15"d | Gov Connection | 1 |
| Miscellaneous Parts | 88732-9000 | USB Cables / IEEE 1394 Cables USB A-TO-B Shielded 0.82m | Mouser | 5 |
| Miscellaneous Parts | PAA632 | APT DC Servo Motor Cable for Z8 Motors, DE15 Male to DE15 Female, 2.5 m | Thorlabs | 1 |
| Miscellaneous Parts | CPA1 | 30 mm Cage System Alignment Plate with Ø1-mm Hole | Thorlabs | 1 |
| Miscellaneous Parts | SCPA1 | 16-mm Cage Alignment Plate | Thorlabs | 1 |
| Miscellaneous Parts | SM1A7 | SM1 Series Alignment Disk | Thorlabs | 1 |
| Miscellaneous Parts | 91290A221 | Black-Oxide Alloy Steel Socket Head Screw, M5×0.8-mm Thread, 6-mm Long | McMaster | 1 |
| Miscellaneous Parts | WLH5-25 | Heat-Proof Wire Springs - WLH (50% Deflection)- | MiSUMi | 10 |
| Miscellaneous Parts | 70031714 | connector; rf coaxial; bnc crimp straightjack; for rg179; 187 cable; 75 Ω | Allied electronics | 10 |
| Miscellaneous Parts | 70090253 | Cable Assembly; SMA Plug; BNC Plug; RG-316 Cbl; 12" | Allied electronics | 1 |
| Miscellaneous Parts | 88000104-1.5 | Cable, RG-174 50 Ωcoaxial with SMA and BNC plugs, 1.5 ft. | pulse research lab | 1 |
| Miscellaneous Parts | 88001350-1.5 | Cable, RG-174 50 Ω coaxial with SMA plugs, 1.5 ft. | pulse research lab | 1 |
| Miscellaneous Parts | 17000003 | Termination, 50 Ω, SMA Male | pulse research lab | 1 |
| Miscellaneous Parts | 88000104-1.5 | Cable, RG-174 50 Ω coaxial with SMA and BNC plugs, 1.5 ft. | pulse research lab | 3 |
| Miscellaneous Parts | 88000071-1.5 | Cable, RG-174 50 Ω coaxial with BNC plugs, 1.5 ft. | pulse research lab | 1 |
| Electronic devices | 781103-01 | PCIe-7852R R Series Multifunction RIO Device, Virtex-5 LX50, 8 AI, 8 AO, 96 DIO, 750kS/sec | National Instruments | 1 |
| Electronic devices | 960680-100 | NI Standard Service Program for Hardware Contact Customer Service for | National Instruments | 1 |

| | | | | |
|---|---|---|---|---|
| | | Availability Duration: 3 Year(s) For PN 781103-01 | | |
| Electronic devices | 781100-01 | PCIe-7841R R Series Multifunction RIO Device, Virtex-5 LX30, 8 AI, 8 AO, 96 DIO, 200kS/sec | National Instruments | 1 |
| Electronic devices | 960680-100 | NI Standard Service Program for Hardware Contact Customer Service for Availability Duration: 3 Year(s) For PN 781100-01 | National Instruments | 1 |
| Electronic devices | 184749-02 | SH68-68-EP, Shielded Cable, 2 m | National Instruments | 1 |
| Electronic devices | 782536-01 | SCB-68A Noise Rejecting, Shielded I/O Connector Block | National Instruments | 3 |
| Electronic devices | 189588-02 | SHC68-68-RMIO Shielded Cable for the Reconfigurable MIO Connector, 68 pin DType to 68 pin VHDCI, 2 m | National Instruments | 2 |
| Electronic devices | 776249-02 | RTSI Bus Cables for 2 PCI or AT/ISA Devices | National Instruments | 1 |
| Electronic devices | 191667-01 | SHC68-68-RDIO Shielded Cable for DIO connector of the 7831R boards, 68pin DType to 68pin VHDCI, 1 m | National Instruments | 4 |
| Electronic devices | Multi-DM5.5 | Multi-DM MEMS Deformable Mirros w/4 meter cables | Boston MicroMachines | 2 |
| Electronic devices | SC-30 | Resonant scanner | EOPC | 1 |
| Electronic devices | Mirror | Silver coated mirror | EOPC | 1 |
| Electronic devices | Driver | AGC-CUSTOM driver includes TTL mute, (3) BNC connections | EOPC | 1 |
| Electronic devices | PRL-414B | 1:4 TTL/CMOS Fanout Buffer and Line Driver | Pulse Research Lab | 1 |
| Electronic devices | PRL-350TTL-NIM | 2-Channel Comparator, TTL Outputs, NIM-compatible Inputs | Pulse Research Lab | 2 |
| Electronic devices | PRL-420ND | 2-Channel Translator, TTL to NECL | Pulse Research Lab | 1 |
| Electronic devices | | Single Axis Scan Unit Set, | ScanLab America | 2 |
| Electronic devices | dynAXIS XS | Power Supply +15 V / -15 V, | ScanLab America | 1 |
| Electronic devices | H10682-01 | Photon Counting Module | Hamamatsu | 1 |
| Electronic devices | C8137 | Power Supply Unit | Hamamatsu | 1 |
| Electronic devices | X10468-02 | LCOS-SLM X10468-02 (750-850 nm) 800×600 px, DiElectric Mirror | Hamamatsu | 1 |

| | | | | |
|---|---|---|---|---|
| Electronic devices | SPCM-AQRH-13-FC | Excelitas SPCM, <250 c/sec w/FC Output pulse width 10 ns | Excelitas/Pacer | 3 |
| Electronic devices | SPCM-POWER SUPPLY | US plug power supply for SPCM 490-SDI18-5-U-P6/552-AC30UNA-R | Excelitas/Pacer | 3 |
| Electronic devices | 0005-00014 | MTS40-B2A3-750.850 Modulator/Shifter Te02-S, 40 +/- 1MHz, 3×3 $mm^2$, 750 - 850 nm | Quanta-Tech | 1 |
| Electronic devices | MPDS1C-B65-30-39.41 | MPDS1C-B65-30-39.41 Multi Purposes Digital Synthesizer, 1 channel, 39-41 MHz, 24VDC, MOD IN: 0-10V/10Kohms (or 0-5V), BLANKING (BLK): TTL/10Kohms, nom 1W, USB/Bluetooth | Quanta-Tech | 1 |
| Electronic devices | 1016-00003 | CBL-SAM150SAM-RG223 Coaxial cable SMAm/RG223/SMAm, Length 1.5 m | Quanta-Tech | 1 |
| Electronic devices | 0012-00033 | AOTFnC-400.730-CPCh-TN Polychromatic Modulator 400-730 nm, Frequency range 46-111 MHz, Aperture 2.5×2.5 $mm^2$, STAB-TN | Quanta-Tech | 1 |
| Electronic devices | MPDS8C-B65-22-46.111 | MPDS8C-B65-22-46.111 Multi Purpose Digital Synthesizer, 8 channels, 46-111 MHz, 24VDC, MOD IN+BLANKING 0-10 V/10 KΩ(or 0-5 V), nom RF power 22 dBm/channel | Quanta-Tech | 1 |
| Electronic devices | 1016-00005 | CBL-SAM200SAM-RG223 Coaxial cable SMAm/RG223/SMAm, Length 2 m | Quanta-Tech | 1 |
| Electronic devices | 1016-00009 | CBL-SCF200SCF-RG316 Coaxial cable SMCf/RG316/SMCf, Length 2 m | Quanta-Tech | 1 |
| Electronic devices | DCC1545M | USB 2.0 CMOS Camera, 1280×1024, Monochrome Sensor | Thorlabs | 1 |
| Electronic devices | 4001-00000 | PS-50-24 Power supply Output: 24 VDC +/- 1%, max 2.2 A Input: 88-264 VAC | Quanta-Tech | 2 |

| | | | | |
|---|---|---|---|---|
| Electronic devices | DCC1545M | USB 2.0 CMOS Camera, 1280×1024, Monochrome Sensor | Thorlabs | 1 |

**Supplementary Table 2 | Connectors on FPGA cards.**

| Function | Connector | Terminal | Pinout | Range (V) |
|---|---|---|---|---|
| FPGA for system control (National Instruments PCIe-7841R) | | | | |
| galvanometer mirror #1 – set position | 0 | 50 | AO 5 | -5 to +5 |
| galvanometer mirror #1 – temperature OK | 0 | 37 | DIO 1 | TTL input |
| galvanometer mirror #1 – power OK | 0 | 38 | DIO 2 | TTL input |
| galvanometer mirror #1 – position OK | 0 | 39 | DIO 3 | TTL input |
| galvanometer mirror #2 – set position | 0 | 49 | AO 6 | -5 to +5 |
| galvanometer mirror #2 – temperature OK | 0 | 40 | DIO 4 | TTL input |
| galvanometer mirror #2 – power OK | 0 | 41 | DIO 5 | TTL input |
| galvanometer mirror #2 – position OK | 0 | 42 | DIO 6 | TTL input |
| resonant mirror – amplitude | 0 | 48 | AO 7 | 0 to 5 |
| resonant mirror – mute | 0 | 43 | DIO 7 | TTL output |
| resonant mirror – synchronization | 1 | 35 | DIO 0 | TTL input |
| PI P541 piezo stage – *z* position | 0 | 52 | AO 3 | 0 to 10 |
| AOTF – channel #1 transmittance | 0 | 53 | AO 2 | 0 to 10 |
| AOTF – channel #2 transmittance | 0 | 54 | AO 1 | 0 to 10 |
| AOTF – channel #3 transmittance | 0 | 55 | AO 0 | 0 to 10 |
| AOTF – blanking | 1 | 43 | DIO 8 | TTL output |
| AOM – transmittance | 0 | 51 | AO 4 | 0 to 5 |
| AOM - blanking | 0 | 36 | DIO 0 | TTL output |
| upper shutter (SH1) – power supply | 1 | 28 | +5 V | +5 |
| upper shutter (SH1) – open/close | 1 | 29 | DIO 28 | TTL output |
| upper shutter (SH1) – status | 1 | 62 | DIO 27 | TTL input |
| lower shutter (SH2) – power supply | 1 | 27 | +5 V | +5 |
| lower shutter (SH2) – open/close | 1 | 61 | DIO 26 | TTL output |
| lower shutter (SH2) – status | 1 | 60 | DIO 25 | TTL input |
| FPGA for signal collection (National Instruments PCIe-7852R) | | | | |
| PMT | 1 | 35 | DIO 0 | TTL input |
| APD #1 | 1 | 36 | DIO 1 | TTL input |
| APD #2 | 1 | 37 | DIO 2 | TTL input |
| APD #3 | 1 | 38 | DIO 3 | TTL input |

Abbreviations: AO, analog output; AOTF, acousto-optic tunable filter; DIO, digital input/output; FPGA, field programmable gate array; TTL, transistor-to-transistor logic.

**Supplementary Table 3 | Filters in chromatic dispersion compensation module.**

| Vendor | Part number |
|---|---|
| Chroma | REP493lp |
| Chroma | AT600lp |
| Chroma | ET655lp |

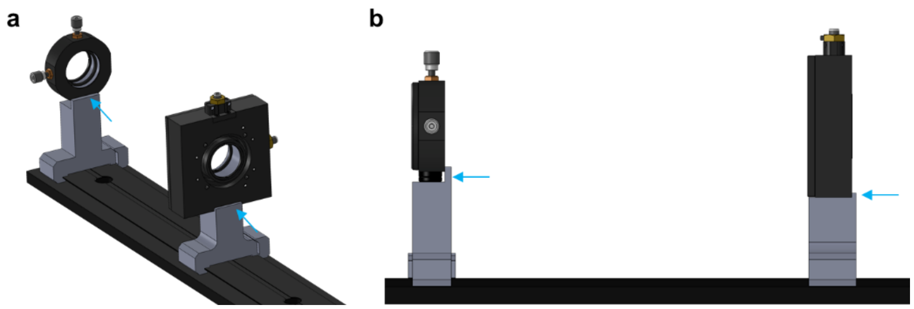


**Supplementary Fig. S1 | An example design of datum plane in lens holder. a**, Overview of CAD rendering. **b**, Side view**.** Blue arrows indicate the datum planes, where the optomechanical component (black) touches the holder (gray).

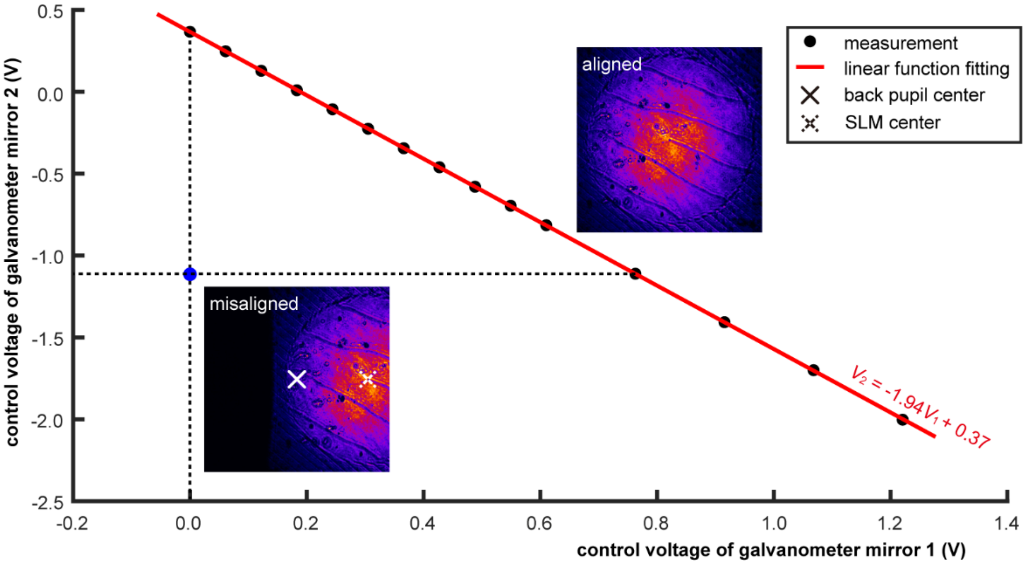


**Supplementary Fig. S2 | Control voltage of galvanometer mirror 1 (*$V_1$*) versus that of galvanometer mirror 2 (*$V_2$*).** The difference between the actual fitted function (red line) and the theoretical one ($V_2 = -2V_1$) reflects errors in the mechanical installation. Any other combination of control voltage signals beyond the actual fitted function will result in the center of hologram on SLM not coinciding with that of the back pupil of the objective lens.

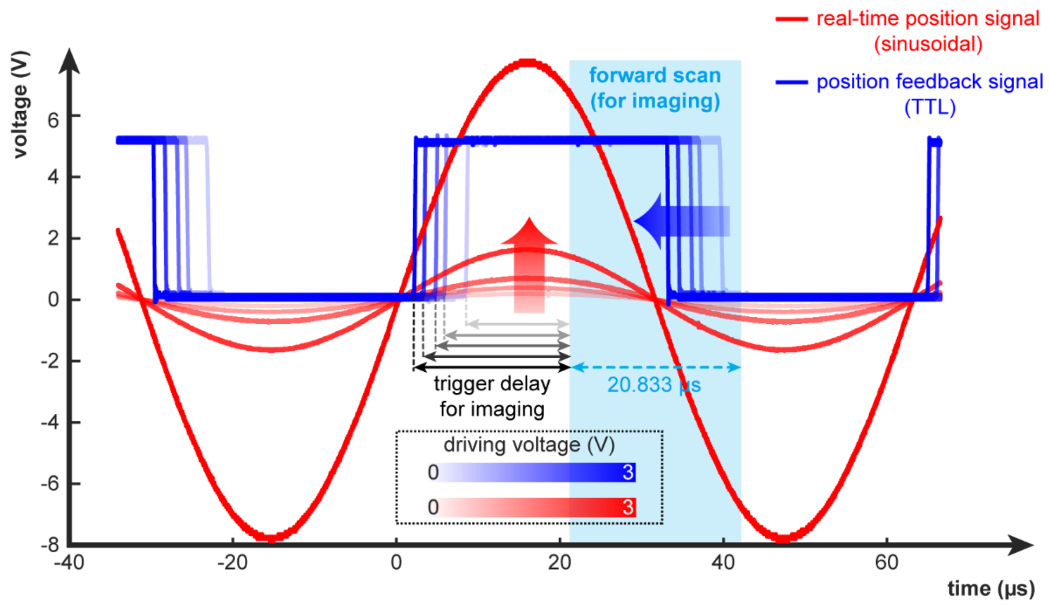


**Supplementary Fig. S3 | Outputs of resonant mirror (RM) during scanning.** By driving the RM with a higher voltage, its resonance frequency remains constant, while the electrostatic force acting upon it intensifies. Consequently, the RM undergoes larger oscillations, leading to a wider FOV. This expanded FOV facilitates applications that require a broader scanning range. Simultaneously, there is a subtle yet significant leftward shift along the temporal axis of the position feedback signal. This temporal shift alludes to a shift in phase between these signals, requiring calibration of the trigger delay for different FOVs (see **Supplementary Fig. S4**). Within our nanoscope control software, the trigger delay is controlled by the internal parameter ‘Trigger Delay (Ticks)’. The calibration data was fitted with a fifth-order polynomial function. The resulting fitting coefficients, which define the polynomial relationship between resonant mirror amplitude and ‘Trigger Delay (Ticks)’, are stored in the file ‘Res Mirror Amplitude to Trigger Delay.vi’ located in the ‘Control Software > Host VIs > General Sub Vis’ directory. This file is essential for automated trigger delay adjustments, ensuring accurate temporal alignment during image acquisition across different FOVs.

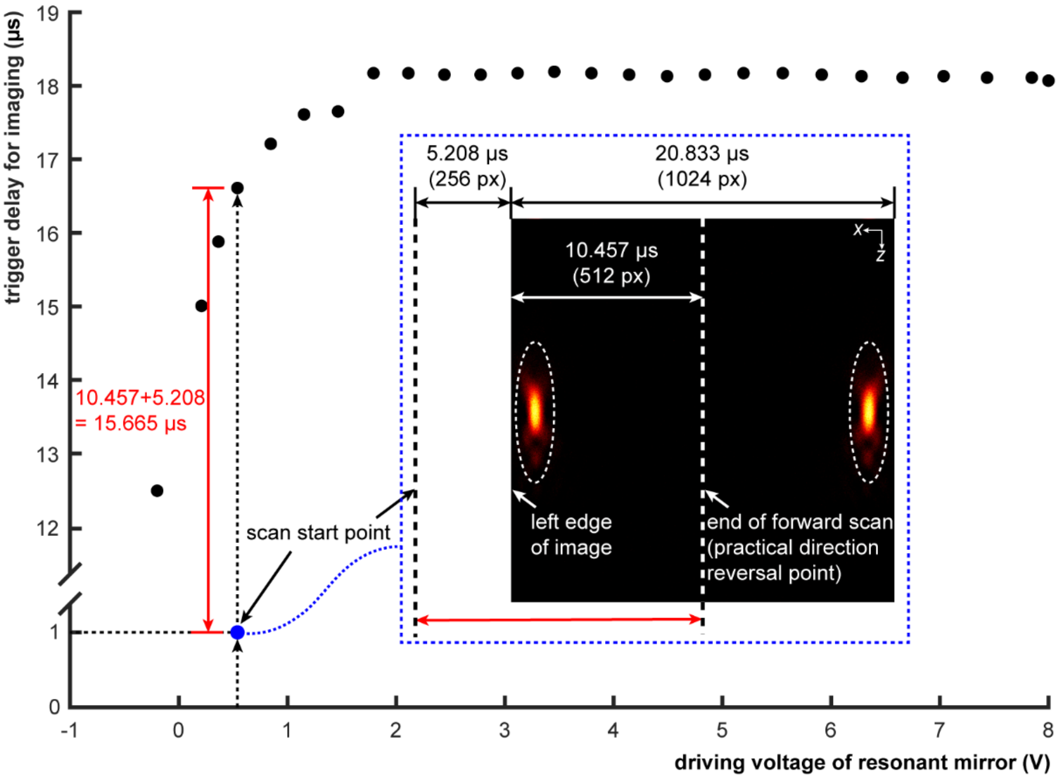


**Supplementary Fig. S4 | Calibration of trigger delay based on imaging.** The imaging process with a 1-μs acquisition trigger delay reveals symmetrical patterns in the gold nanoparticle images, suggesting that the scanning direction of the galvanometer is reversed during acquisition. This direction reversal delineates a symmetry axis, *i.e.* the end of forward scan or the direction reversal point, between the mirrored gold nanoparticle images. To identify this axis precisely, the image is divided into two subgraphs. One subgraph is flipped horizontally and correlated with the other to find the maximum correlation. Utilizing known data about the resonant mirror steering position and pixel dwell time, it becomes feasible to determine the time taken for the galvanometer to move to the left edge of the image. By comparing this to the ideal galvanometer position (1/12 of the galvanometer's cycle time, *i.e.* 5.208 μs), the discrepancy between the scan start point and the direction reversal point is calculated. Incorporating the 1-μs initial bias delay yields the trigger delay as a function of the drive voltage. Notably, the trigger delay escalates with increasing drive voltage (within the 0-2 V range), yet stabilizes beyond 2 V. Further increases in drive voltage yield marginal alterations in trigger delay.

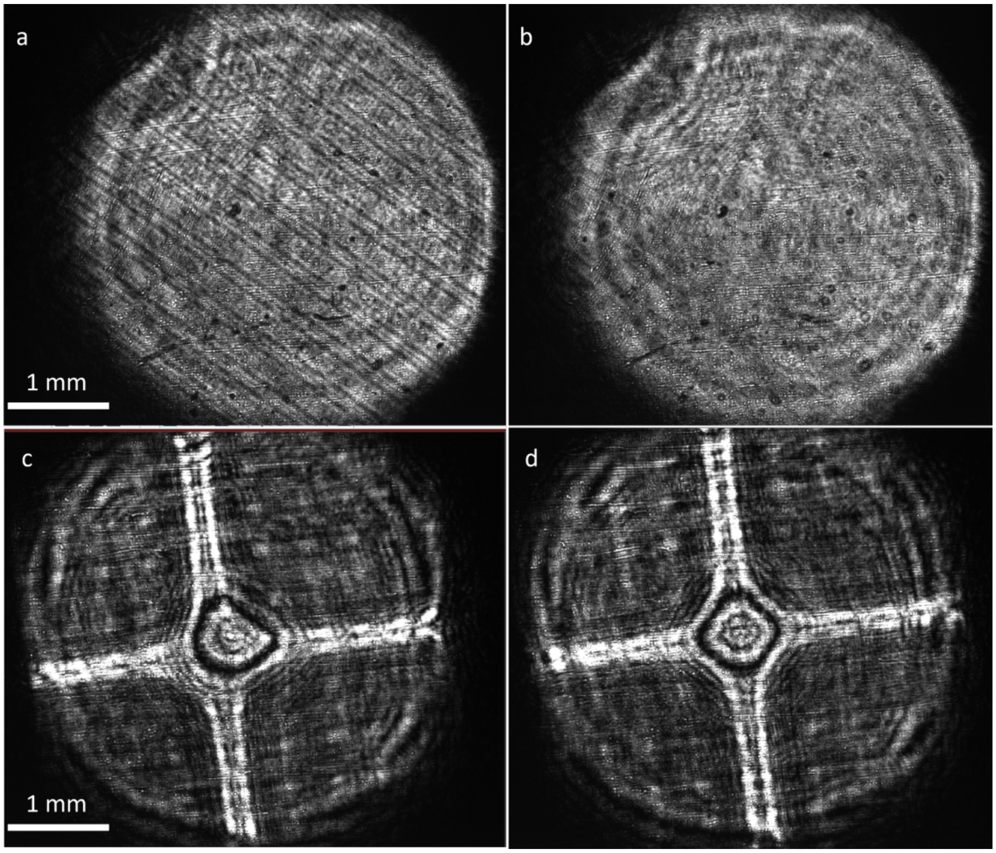


**Supplementary Fig. S5 | DM conjugation to objective back aperture (a, b) and co-centering to SLM holograms (c, d). a-b**, images of DM surface at back aperture of objective. When unconjugated, a grid pattern is visible (**a**), which disappears upon conjugation (**b**). **c-d,** images of both DM and SLM at the position beyond the back aperture of objective. To highlight their corresponding centers, a top-hat and a cross pattern are loaded onto the SLM and DM, respectively. An asymmetric image is evident when the co-centering relationship is not satisfied (**c**), whereas symmetry is restored upon achieving co-centering (**d**).

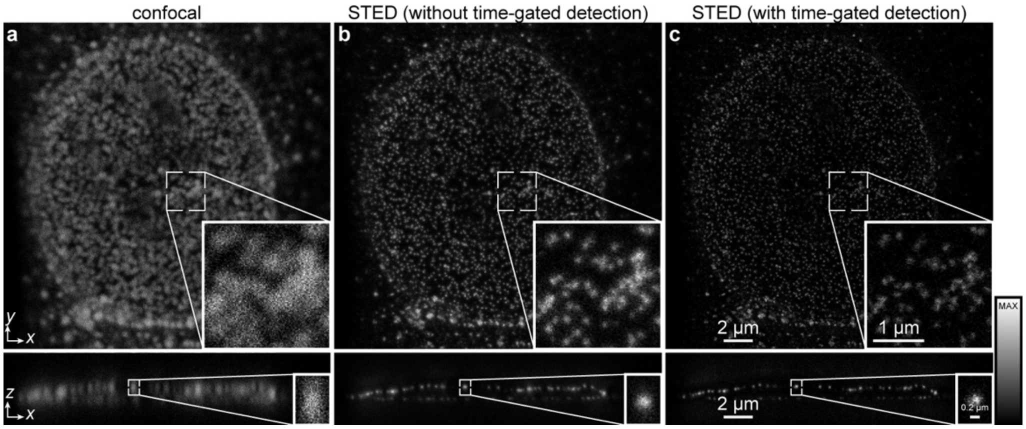


**Supplementary Fig. S6 | Effects of time-gated detection on imaging performance. a**, confocal image of nuclear pore complexes in a vero-B4 cell immunolabelling MAB414 with STAR RED. **b**-**c**, STED imaging of the same field of view without (**b**) or with (**c**) time-gated detection. The figures in the top and the bottom rows show $xy|_{z=0}$ (top) and $xz|_{y=0}$ (bottom) cross sections, respectively. Each figure is normalized to its own peak intensity.

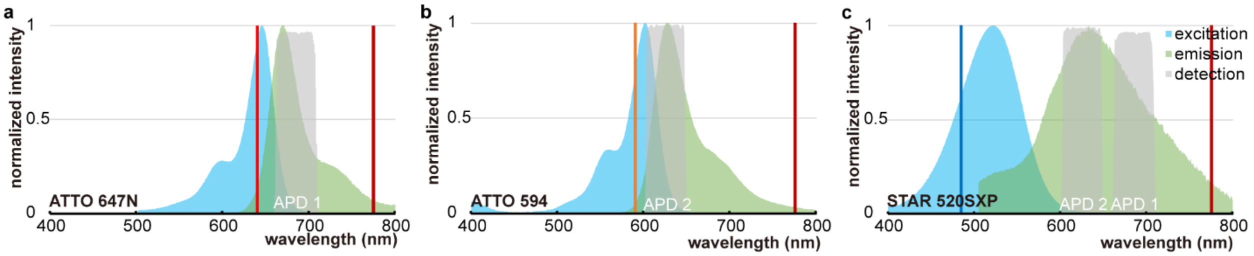


**Supplementary Fig. S7 | Excitation (blue) and emission (green) spectra of fluorophores recommended for isoSTED nanoscopy. a**, ATTO 647N. **b**, ATTO 594. **c**, STAR 520XP.

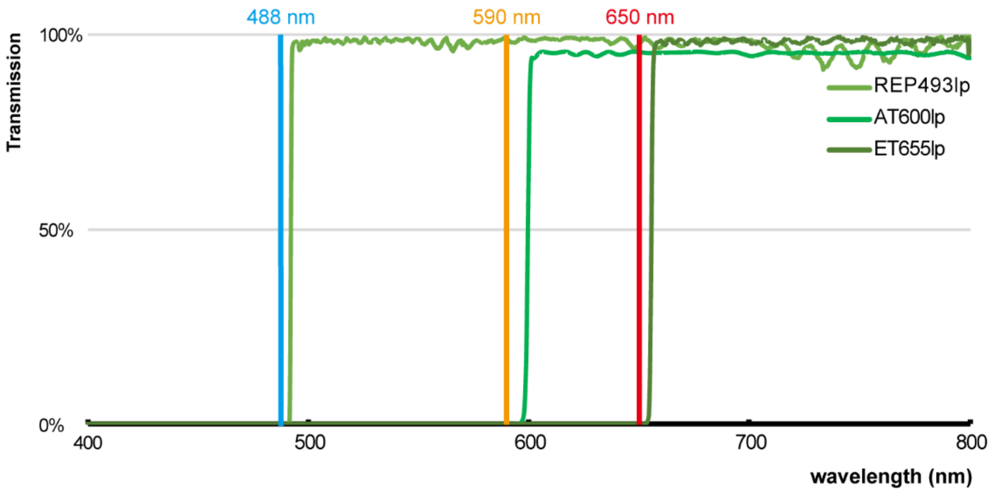


**Supplementary Fig. S8 | Spectral transmission of filters in chromatic dispersion compensation module.**

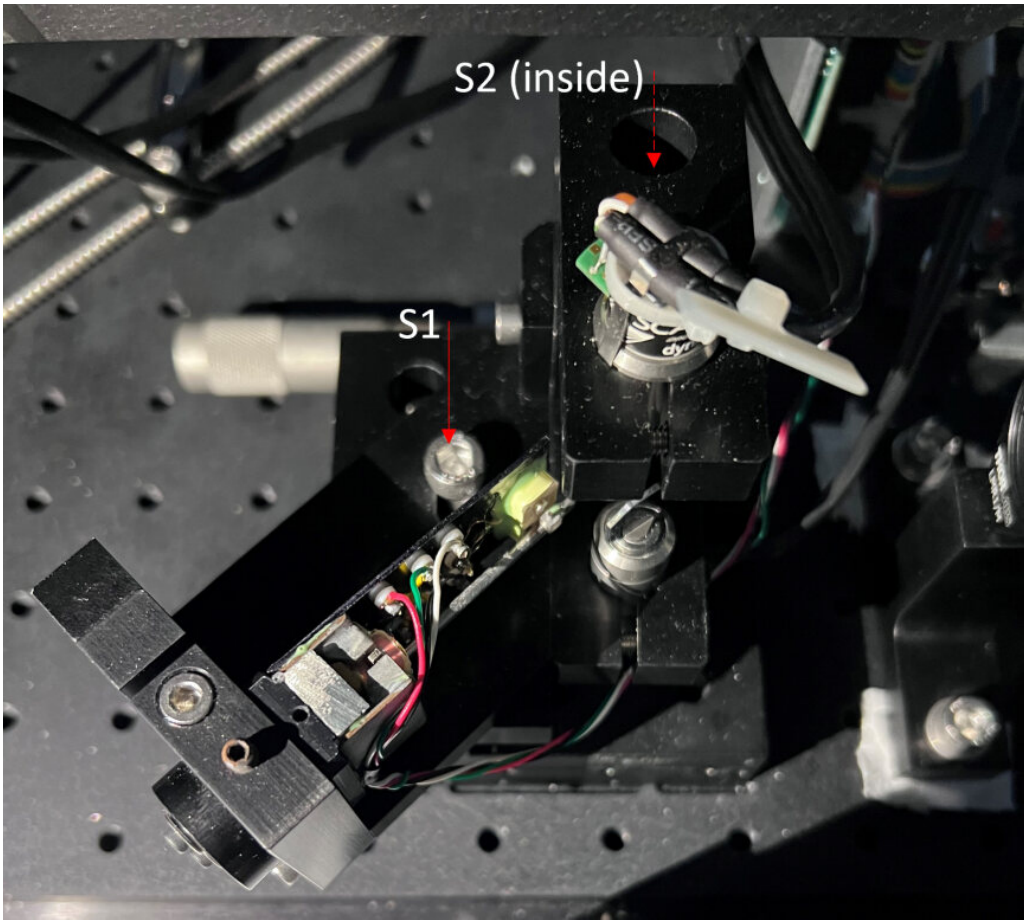


**Supplementary Fig. S9 | Customized holder for scanner.** Loosening screws (S1 and S2) enables translation of all the three scan mirrors together.

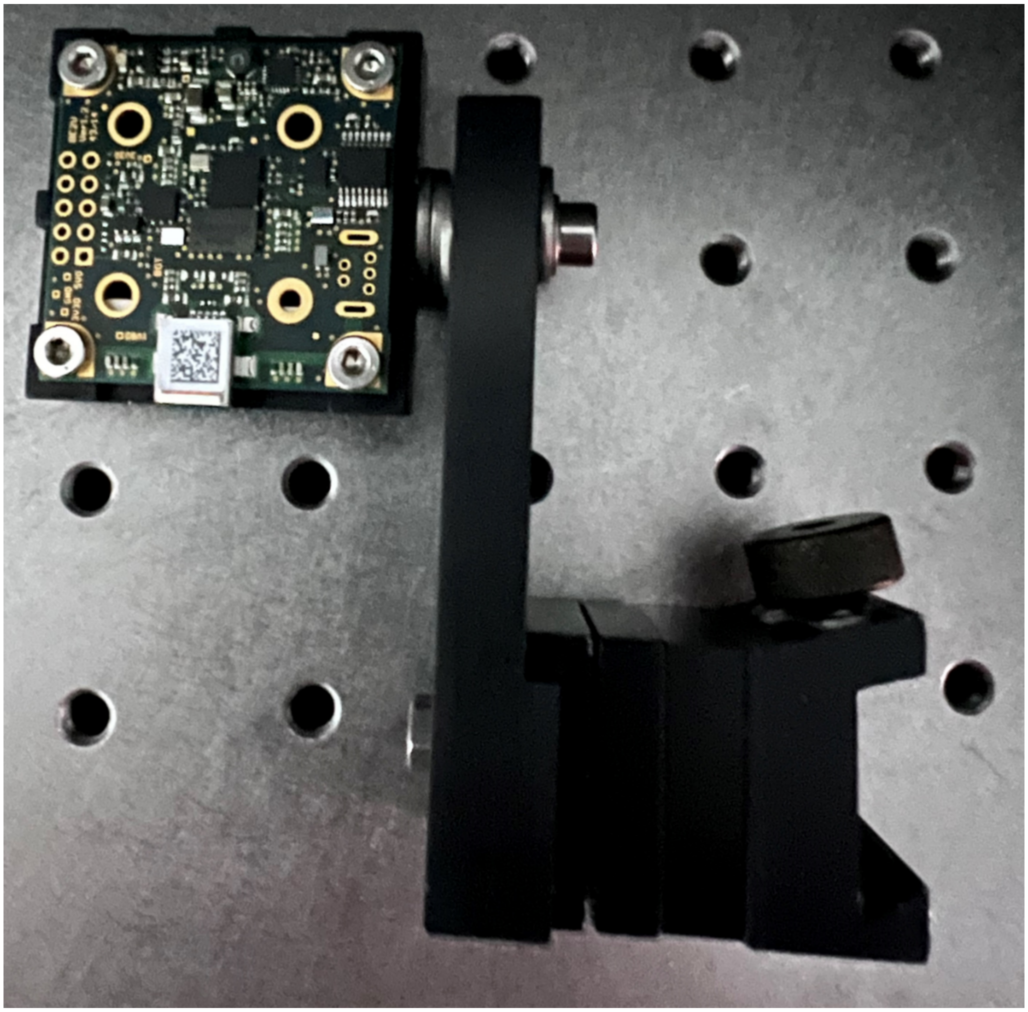

**Supplementary Fig. S10 | Alignment camera module for rails (R5, R6) in 4Pi interference cavity.** The camera is assembled with a kinematic base (Thorlabs, KB1X1), a dovetail carrier (Thorlabs, RC1), and a customized connector (FAB-IS0054). The kinematic base provides flexible and repeatable mounting of the camera when sliding RC1 on the rails.

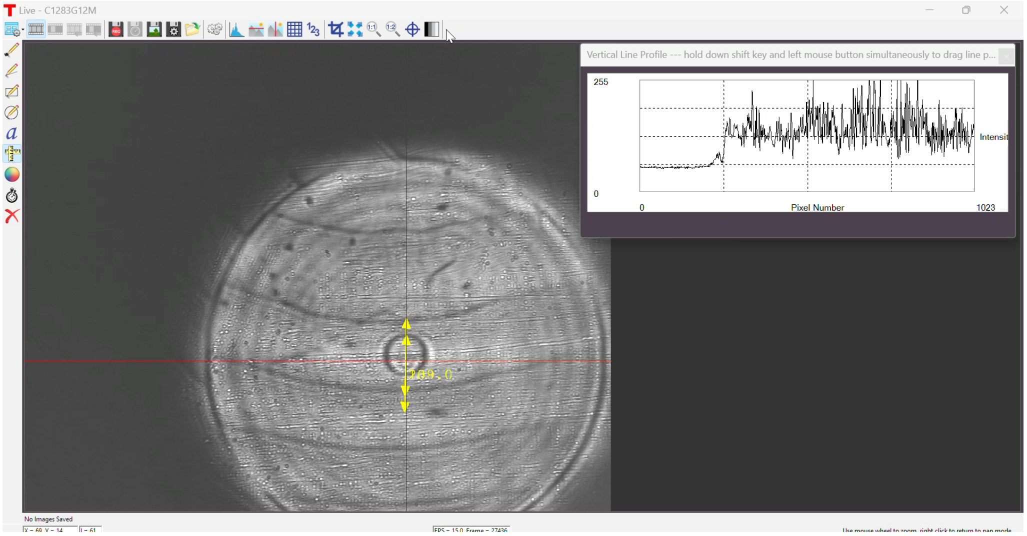


**Supplementary Video S1 | Image of Galvanometer mirror at back pupil of objective when the mirror is unconjugated with the DM.** Scanning scale of the Galvanometer mirror corresponds to an image size of ~30 μm at the focal plane in the sample.

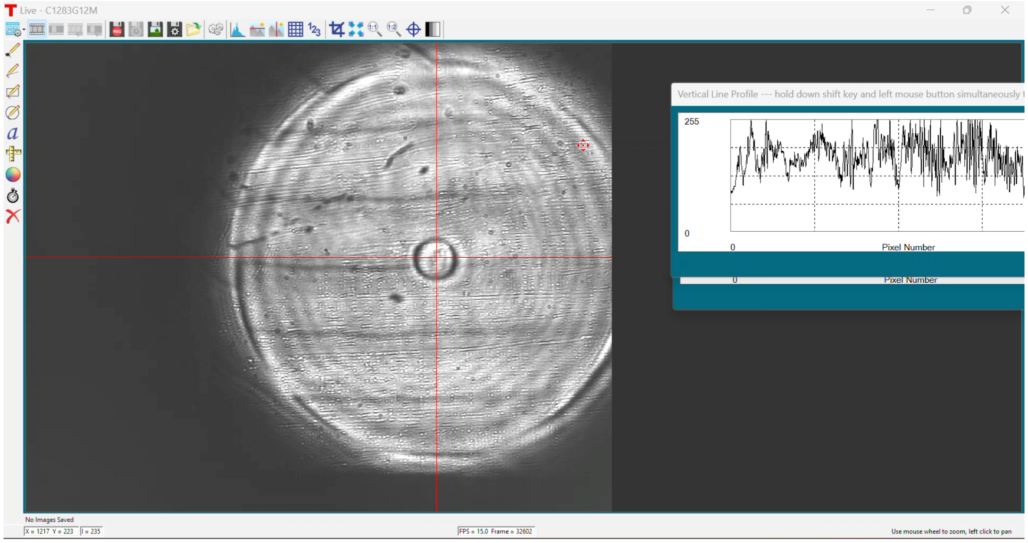


**Supplementary Video S2 | Image of Galvanometer mirror at back pupil of objective when the mirror is conjugated with the DM.** Scanning scale of the Galvanometer mirror corresponds to an image size of ~30 μm at the focal plane in the sample.
